# Interchromosomal Colocalization with Parental Genes Is Linked to the Function and Evolution of Mammalian Retrocopies

**DOI:** 10.1101/2023.07.12.548661

**Authors:** Yubin Yan, Yuhan Tian, Zefeng Wu, Kunling Zhang, Ruolin Yang

## Abstract

Retrocopies are gene duplicates arising from reverse transcription of mature mRNA transcripts and their insertion back into the genome. While long being regarded as processed pseudogenes, more and more functional retrocopies have been discovered. How the stripped-down retrocopies recover expression capability and become functional paralogs continually intrigues evolutionary biologists. Here, we investigated the function and evolution of retrocopies in the context of three-dimensional (3D) genome organization. By mapping retrocopy-parent pairs onto sequencing-based and imaging-based chromatin contact maps in human and mouse cell lines and onto Hi-C interaction maps in five other mammals, we found that retrocopies and their parental genes show a higher-than-expected interchromosomal colocalization frequency. The spatial interactions between retrocopies and parental genes occur frequently at loci in active subcompartments and near nuclear speckles. Accordingly, colocalized retrocopies are more actively transcribed and translated, and are more evolutionarily conserved than noncolocalized ones. The active transcription of colocalized retrocopies may result from their permissive epigenetic environment and shared regulatory elements with parental genes. Population genetic analysis of retroposed gene copy number variants (retroCNVs) in human populations revealed that retrocopy insertions are not entirely random in regard to interchromosomal interactions and that colocalized retroCNVs are more likely to reach high frequencies, suggesting that both insertion bias and natural selection contribute to the colocalization of retrocopy-parent pairs. Further dissection implies that reduced selection efficacy, rather than positive selection, contributes to the elevated allele frequency of colocalized retroCNVs. Overall, our results hint a role of interchromosomal colocalization in the “resurrection” of initially neutral retrocopies.

## Introduction

Gene duplication is a major process leading to the birth of new genes, which contributes significantly to the adaptive evolution and phenotypic novelty of species in diverse clades (Ohno 1970; Zhang 2003; Chen et al. 2013). Gene retrotransposition is an RNA-based gene duplication mechanism whereby mRNA derived from protein-coding genes are reverse transcribed to DNA and inserted back into the genome (Kaessmann et al. 2009). In mammals, gene retroduplication is mainly mediated by the non-LTR retrotransposon LINE-1s (L1s), and occasionally by LTR-retrotransposons (Esnault et al. 2000; Tan et al. 2016).

Although RNA-based gene duplication is less common than DNA-based duplication (i.e., segmental duplication and whole genome duplication) (Kaessmann 2010), thousands of retrotransposed gene copies (retrocopies) have been identified in humans and other mammalian species (Navarro and Galante 2015; Carelli et al. 2016; Rosikiewicz et al. 2017). However, to what extent this large amount of retrocopies contribute to new gene origination and genome evolution is still an open question. On the one hand, retrocopies have been stripped of introns and promoters, rendering them to be “dead on arrival”, and were frequently dismissed as “processed pseudogenes” (Mighell et al. 2000; Zhang et al. 2003). On the other hand, increasing number of protein-coding retrocopies (retrogenes) with explicit functions have been reported in various organisms (Casola and Betran 2017). For example, a recent study reported that a mammalian retrogene *HAPSTR2* encodes a protein that retains the biochemical features of its parental gene *HAPSTR1* and acts as a safeguard in stress signaling and resilience (Amici et al. 2023). Moreover, accumulating evidence indicates that many retrocopies are robustly expressed, often in a tissue-specific manner (Carelli et al. 2016; Qian et al. 2022). These retrocopy-derived RNAs may function as natural antisense transcripts (NATs), long noncoding RNAs (lncRNAs), or sponging microRNAs to regulate their progenitors or host genes (Kubiak and Makałowska 2017; Ciomborowska-Basheer et al. 2021). Ribosome profiling data further suggest that a sizable proportion of retrocopies may be actively translated (Kim et al. 2014; Ji et al. 2015; Qian et al. 2022). However, the specific proteins they produce and their functions remain subjects for further investigation.

How some retrocopies gain their expression capability and “awake” as functional paralogs continually attracts evolutionary biologists (Vinckenbosch et al. 2006). Previous research discovered that retrocopies may exploit preexisting promoters, evolve new promoters from scratch, or recruit proto-promoters from their genomic vicinity (Carelli et al. 2016). For example, the promoter of the aforementioned retrogene *HAPSTR2* originated from a nearby proto-promoter (Amici et al. 2023). However, current studies mostly focused on the role of local genomic context in the expression of retrocopies (Carelli et al. 2016; Machado and Antunes 2020), while the influences of high-order chromatin organization on retrocopy expression and functionality are largely unexplored.

In eukaryotes, high-order chromatin organization and transcription are tightly linked, with reciprocal influence on each other (Bonev and Cavalli 2016; van Steensel and Furlong 2019). Apart from the well-studied intrachromosomal (*cis*) structures, such as A/B compartments, topologically associating domains (TADs), and chromatin loops, interchromosomal (*trans*) interactions also play a critical role in gene regulation (Quinodoz et al. 2018; Maass et al. 2019) and genome evolution (Dai et al. 2014; Xie et al. 2016). For example, human interchromosomal paralogous gene pairs generated by whole genome duplication (i.e., ohnolog pairs) tend to be colocalized in the nucleus, which may associate with stronger dosage balance in those ohnologs (Xie et al. 2016). Likewise, neighboring genes in yeast exhibit interchromosomal spatial proximity after their separation (Dai et al. 2014), which indicates a role for interchromosomal interactions in the coregulation and evolution of functionally related genes. Inspired by these previous studies and the fact that most retrocopies and their parental genes are located on different chromosomes, we sought to ask whether retrocopy-parent pairs show interchromosomal colocalization in the nucleus, and if so, how spatial proximity links to the function and evolution of retrocopies.

Here, we used chromatin contact maps obtained by orthogonal techniques in human and mouse cell lines and Hi-C interaction matrices in chimpanzees, rhesus macaques, marmosets, cows, and dogs to detected significant interchromosomal interactions between retrocopies and their parental genes. Through extensive simulations, we demonstrated that mammalian retrocopy-parent pairs exhibit a higher colocalization frequency than random chromatin fragment pairs under various null models. Compared to noncolocalized retrocopies, colocalized retrocopies are enriched in active subcompartments and speckles, and are more actively transcribed and translated, supporting a connection between spatial proximity and functionality of retrocopies. Such differences might be resulted from the higher number of shared regulatory elements with parental genes and the more active epigenetic landscape of colocalized retrocopies. Furthermore, by investigating retroposed gene copy number variants (retroCNVs) in human populations, we discovered that insertion bias and nonadaptive evolution shaped the colocalization between retrocopies and parental genes. Our results shed lights on the function and evolution of retrocopies in the context of 3D genome organization.

## Results

### Human Interchromosomal Retrocopy-Parent Pairs Exhibit Widespread Nuclear Colocalization

To determine the extent of spatial colocalization between retrocopies and their parental genes, we retrieved 5,159 human retrocopy-parent pairs from Carelli et al. (2016).After filtering steps (Materials and Methods), 4,427 interchromosomal pairs were retained for subsequent analyses. Using Hi-C contact maps at 250-kb resolution of seven human cell lines (GM12878, HMEC, HUVEC, IMR90, K562, KBM7, and NEHK) (Rao et al. 2014), we found that between 738 (16.7%; in IMR90) and 1,019 (23.0%; in GM12878) retrocopy-parent pairs exhibit significant interchromosomal interactions, i.e., spatial colocalization (**fig. 1A**, **supplementary fig. S1** and **table S1**). More than half (2,511; 56.7%) retrocopy-parent pairs are colocalized in at least one cell lines, and 1,465 (33.1%) pairs exhibit spatial proximity in two or more cell lines (**fig. 1B**). We reasoned that if the spatial proximity between retrocopies and parental genes had biological significance, then a higher-than-expected colocalization frequency should be observed. Therefore, we performed various simulations to determine the statistical significance of retrocopy-parent colocalization (**supplementary fig. S2**). First, we simulated 1,000 datasets, in each of which 4,427 random fragment pairs with matching chromosomes and sequence lengths as the retrocopy-parent pairs were sampled. Notably, we found that the colocalization frequency of true retrocopy-parent pairs is significantly higher than simulated chromatin pairs (*p* < 0.001; **fig. 1C** and **supplementary fig. S3**), suggesting nonrandom colocalization between retrocopies and parental genes. Second, we kept the information of retrocopies unchanged while sampling random coding and noncoding genes with matching chromosomes as parental genes. Third, we kept the information of parental genes unchanged while sampling random fragments with matching chromosomes and sequence lengths as retrocopies. Forth, similar to the second simulation, except that we constrained the randomly sampled genes to exclusively protein-coding genes. For these simulations, we also observed a significant higher colocalization frequency in true retrocopy-parent pairs (*p* < 0.05, except for simulation 4 of NHEK; **fig. 1C** and **supplementary fig. S3**). However, compared to the first null model, the colocalization frequencies in these simulated data are higher, indicating that some properties (e.g., local genomic and epigenomic environments) of retrocopies or parental genes may contribute to interchromosomal colocalization.

**Figure 1.**
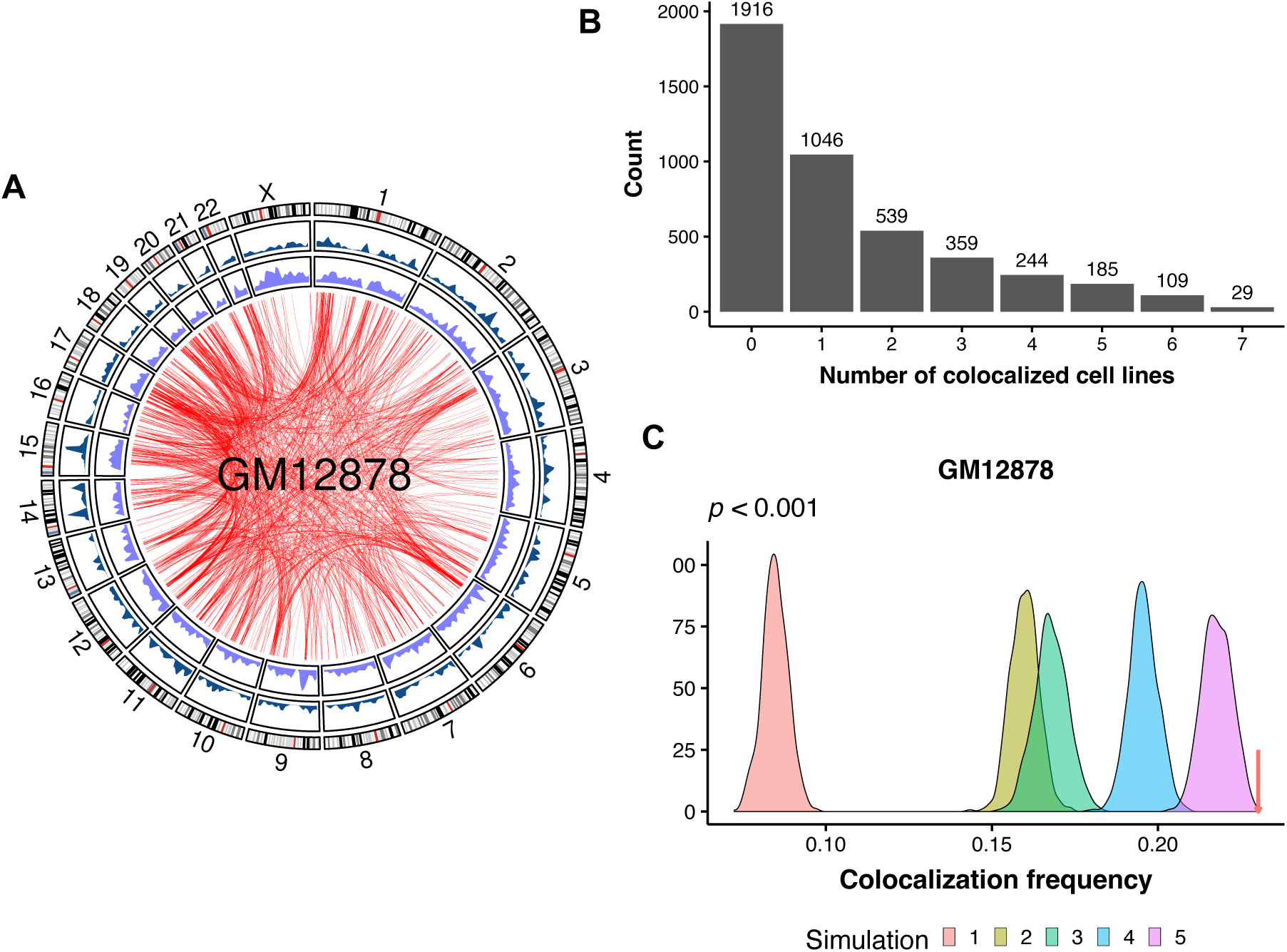
Spatial colocalization between retrocopies and their parental genes. (*A*) Circos plot highlighting the chromatin interactions between retrocopies and parental genes in the GM12878 cell line. From outer to inner are the ideogram of human chromosomes, the density of parental genes, the density of retrocopies, and significant chromatin interactions between retrocopies and parental genes. (*B*) Bar plot showing the number of human interchromosomal retrocopy-parent pairs as a function of the number of cell lines wherein the spatial colocalization is detected. (*C*) Distribution of colocalization frequency between chromatin pairs under different null models, while the red arrow denotes the colocalization frequency of true retrocopy-parent pairs. Simulations and true frequency are based on Hi-C data of GM12878. *P*-value < 0.001 in all cases.

Lastly, to further investigate the influence of retrocopies and parental genes on spatial proximity, we randomly shuffled the relationship between retrocopies and cognate genes for 1,000 times. Particularly, the observed colocalization frequencies in GM12878, HMEC, HUVEC, K562, and NHEK are still significantly higher than expected (*p* < 0.05), while the significance is marginal in IMR90 and KBM7 (*p* = 0.082 and 0.078, respectively; **fig. 1C** and **supplementary fig. S3**). To further evaluate the significance of interchromosomal colocalization between retrocopies and their parental genes, we analyzed contact maps derived from two orthogonal methods: 1) split-pool recognition of interactions by tag extension (SPRITE), which is a sequencing-based but ligation-free technique that enables detection of multiway genome-wide chromatin interactions; and 2) multiplexed error-robust fluorescence in situ hybridization (MERFISH), which is an imaging-based approach to map chromatin structure at genome scale. In GM12878, we observed a significant overlap in the colocalization status of retrocopy-parent pairs derived from Hi-C (Rao et al. 2014) and SPRITE (Quinodoz et al. 2018) when assessed at the same resolution (1 Mb) and significance level (q < 0.05). (**supplementary fig. S4A**). Furthermore, when employing significant contacts identified by SPRITE, retrocopies and their parental genes exhibit a colocalization frequency higher than expected (*p* < 0.05 in all simulations; **supplementary fig. S4B**). Notably, in IMR90, retrocopy- parent pairs identified as colocalized by Hi-C display shorter median spatial distances and higher proximity frequencies, as measured by MERFISH (Su et al. 2020), compared to noncolocalized pairs. (*p* = 3.5 × 10^-6^ and 2.0 × 10^-13^, respectively; **supplementary fig. S5**).

### Colocalized Retrocopies Are Enriched in Active Subcompartments and Are Close to Nuclear Speckles

We next asked where did retrocopies and parental genes colocalize with respect to the 3D genome organization in the nucleus. By clustering interchromosomal contact matrix of the ultra-deep sequenced GM12878 cell line, Rao et al. (2014) divided the active A and inactive B compartments into five primary subcompartments (A1, A2, B1, B2, and B3), which were associated with distinct genomic and epigenomic features. We found that in GM12878, retrocopies are slightly enriched in A2 subcompartment while depleted in B2 and B3 subcompartments (**fig. 2A**). Such enrichment of active subcompartments and depletion of inactive subcompartments is more apparent for parental genes. In addition, compared to protein-coding genes as a whole, retrocopies’ parental genes show stronger enrichment of active subcompartments and depletion of inactive subcompartments, which is consistent with previous findings that highly expressed genes are more prone to generate retrocopies (Zhang et al. 2003; Podlaha and Zhang 2009; Sisu et al. 2014).

**Figure 2.**
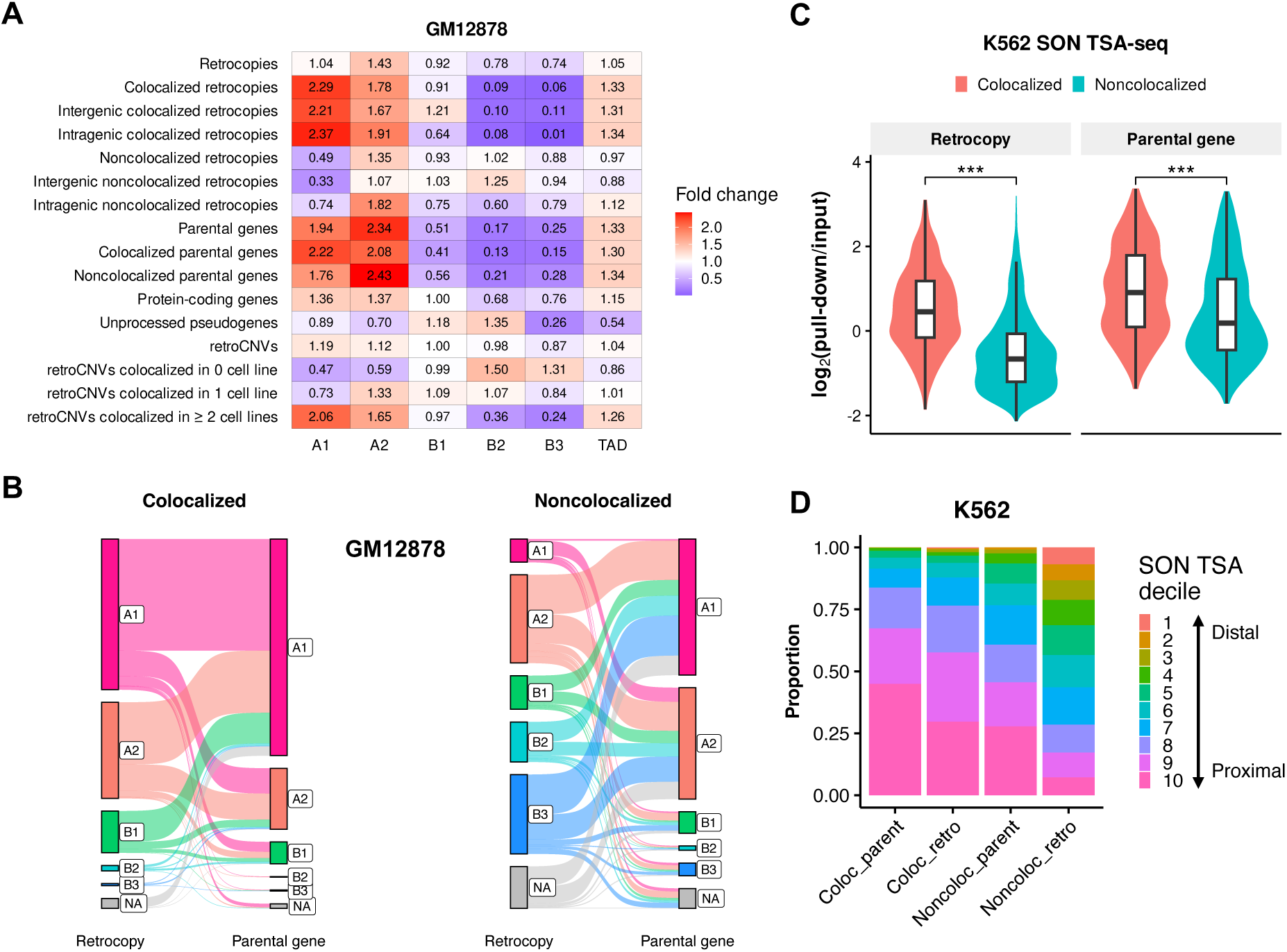
Chromatin positioning of retrocopies and parental genes in the 3D genome. (*A*) Enrichments of retrocopies, parental genes, unprocessed pseudogenes, and retroCNVs in five primary subcompartments and TAD in the GM12878 cell line. (*B*) Sankey diagram showing the connection of colocalized (left panel) and noncolocalized (right panel) retrocopy-parent pairs in the context of subcompartment annotations in GM12878. The height of bar indicates the proportion of loci that fall in respective categories. Lines connect retrocopies and their corresponding parent genes. NA denotes a fragment is not annotated as any of the five subcompartments. (*C*) Comparison of SON TSA-seq signal between colocalized and noncolocalized retrocopies/parental genes in K562 (\*\*\**p* < 0.001; two tailed Wilcoxon test). (*D*) Proportion of colocalized and noncolocalized retrocopies/parental genes that overlapped with different SON TSA-seq deciles in K562. Decile 1 (estimated distance: 0.868-1.020 μm) and decile 10 (estimated distance: 0.054-0.399 μm) represent the top 10% of chromatin that are farthest and closest to SON, respectively, and other deciles are those that fall between. Coloc_retro: colocalized retrocopies; noncoloc_retro: noncolocalized retrocopies; coloc_parent: colocalized parental genes; noncoloc_parent: noncolocalized parental genes.

Remarkably, colocalized retrocopies show strong enrichment of A1 and A2 subcompartments and strong depletion of B2 and B3 subcompartments, resembling the pattern of parental genes (**fig. 2A**). In contrast, noncolocalized retrocopies are depleted in active A1 subcompartment and show little depletion of inactive subcompartments. Dividing colocalized and noncolocalized retrocopies into intergenic and intragenic subclasses do not affect the general pattern (**fig. 2A**), indicating that the subcompartment association is not affected by host genes. Besides, parental genes and colocalized retrocopies are slightly enriched in TADs. Contrary to retrocopies, unprocessed pseudogenes are depleted in A1 and A2 subcompartments and TAD while slightly enriched in B1 and B2 subcompartments (**fig. 2A**), reflecting different 3D chromatin environments between processed and unprocessed “pseudogenes”. Using subcompartment annotations inferred by deep learning on relatively low resolution Hi-C maps (Xiong and Ma 2019), we obtained similar results in HMEC, HUVEC, IMR90, and K562 (**supplementary fig. S6**).

Of note, for colocalized retrocopy-parent pairs, both ends tend to be overlapped with A1 or A2 subcompartments, whereas for noncolocalized pairs, a substantial proportion of retrocopies are overlapped with B2 or B3 subcompartments, despite the fact that the majority of parental genes still overlap with active subcompartments (**fig. 2B** and **supplementary fig. S7**).

Subcompartments are strongly associated with subnuclear structures, such as nuclear lamina, nucleolus, and speckles, which are associated with genome regulation (Chen et al. 2018). Thus, we compared the spatial positioning relative to subnuclear structures of colocalized and noncolocalized retrocopies using TSA-seq data in K562 (Chen et al. 2018). TSA-seq is a technique that combines tyramide signal amplification (TSA) and high-throughput sequencing to map chromosome organization relative to subnuclear structures genome-wide. We found that the log2(pull-down/input) signals of SON and pSC35 TSA-Seq of colocalized retrocopies or parental genes are significantly higher than their noncolocalized counterparts, whereas an opposite trend is observed for LaminAC and LaminB TSA-Seq signals (*p* < 0.001; **fig. 2C** and **supplementary figs. S8A-C**). SON and pSC35 are two marker proteins for nuclear speckles while LaminAC and LaminB are markers for nuclear lamina, and high signal means shorter cytological distance to the respective subnuclear structure. Thus, our results indicate that colocalized retrocopies and parental genes are closer to speckles, while their noncolocalized counterparts are closer to lamina.

SON TSA-Seq signals can be further converted to actual distance (in μm) between chromatin and nuclear speckles and loci can be assigned to different TSA deciles based on their distance to speckles (Chen et al. 2018). Leveraging these data, we found that colocalized retrocopies have an average distance of 0.48μm to speckles, longer than that of colocalized parental genes but significantly shorter than that of noncolocalized retrocopies (*p* < 0.001; **supplementary fig. S8D**). Consistently, colocalized parental genes have the highest proportion of fragments falling in the 9^th^ and 10^th^ SON TSA deciles (chromatin that is mostly close to speckle), followed by colocalized retrocopies and noncolocalized parental genes, while only a small proportion of noncolocalized retrocopies fall in the 9^th^ and 10^th^ deciles (**fig. 2D**). Furthermore, by integrating TSA-Seq, DamID and Hi-C data in the K562 cell line, Wang et al. (2021) partitioned the genome into 10 different spatial compartmentalization states relative to nuclear speckles, lamina, and nucleolus. We found that colocalized retrocopies are enriched in interior active regions and speckle-associated chromatin, while depleted in interior repressive regions and lamina-associated chromatin, similar to the pattern of parental genes (**supplementary fig. S9**). Since both ends of colocalized retrocopy-parent pairs tend to have shorter spatial distances to speckles (**supplementary fig. S10A**) in comparison with noncolocalized pairs, we randomly sampled protein-coding genes with matching speckle distance (± 0.01 μm away) as true parental genes to form simulated chromatin pairs with retrocopies for 1,000 times. We found that after controlling speckle distances, true retrocopy-parent pairs still exhibit higher-than-expected colocalization frequency (*p* < 0.001; **supplementary fig. S10B**), suggesting that speckle proximity cannot fully explaining retrocopy-parent colocalization. However, because there are 10-50 speckles in a nucleus (Spector and Lamond, 2011), we cannot exclude the possibility that retrocopies and their parental genes tend to be close to the same nuclear speckle.

### RNA Biogenesis and Shared Regulatory Elements Are Linked to the Interchromosomal Colocalization between Retrocopies and Parental Genes

Although retrocopies are often thought to be processed pseudogenes, most of them show evidence of transcription (Carelli et al. 2016). The distinct positioning of colocalized and noncolocalized retrocopies in the nucleus suggests they may have different transcription profile, we then asked whether the spatial colocalization with parental genes is associated with the expression of retrocopies. Using RNA-seq data of GM12878, HUVEC, K562, and NHEK from the ENCODE project (Djebali et al. 2012), we have also detected evidence of transcription for most retrocopies (**supplementary table S1**), with a substantial proportion of them showing robust expression. For instance, when assessing polyadenylated (polyA+) RNAs in whole cells, there are 772 to 1,350 (17.4% to 30.5%) retrocopies exhibiting an FPKM (Fragments Per Kilobase per Million mapped fragments) ≥ 1 (**supplementary fig. S11A**). The proportion of robustly expressed (FPKM ≥ 1) retrocopies for polyA+ RNAs is larger than 15% and 20% in the nucleus and cytosol, respectively (**supplementary figs. S11B and C**). There are also substantial non-polyadenylated (polyA-) RNAs transcribed from retrocopies in the nucleus and whole cell, but relatively few in the cytosol (**supplementary figs. S11D-F**). More importantly, colocalized retrocopies and parental genes have significantly higher expression level than their noncolocalized counterparts in most cell lines, for RNAs in different compartments (whole cell, nucleus, and cytosol), and for both ployA+ and polyA- RNAs (*p* < 0.01; **fig. 3A** and **supplementary figs. S12 and S13**). In addition, the GRO-Seq and PolII TSA-Seq data in K562 (Chen et al. 2018) shows that colocalized retrocopies and parental genes generate more nascent RNAs and are closer to RNA polymerase II than their noncolocalized counterparts (*p* < 0.001; **fig. 3B**), suggesting that RNA biogenesis is more active at colocalized loci.

**Figure 3.**
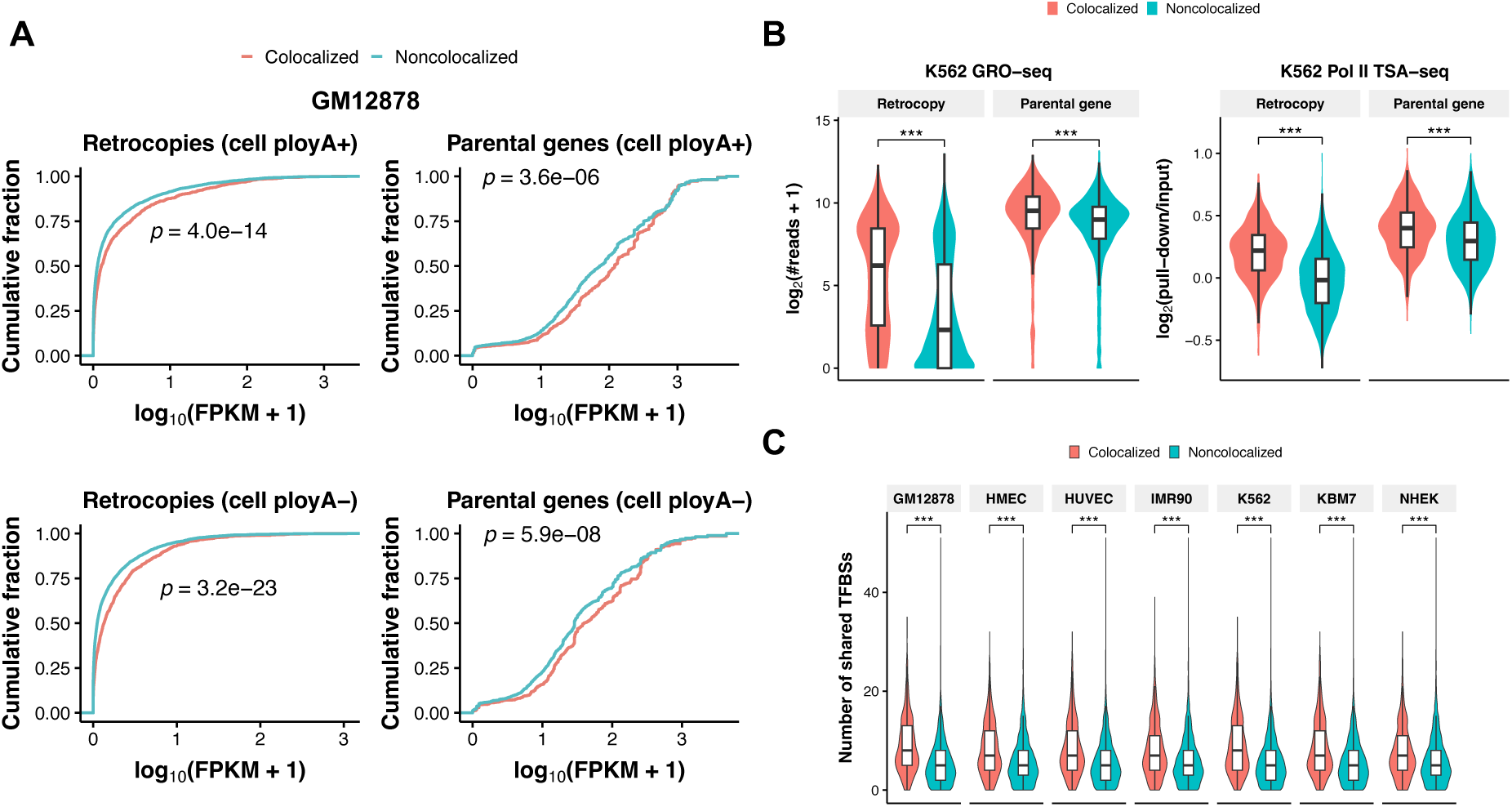
Expression and regulatory element sharing of retrocopies and parental genes. (*A*) Plots of cumulative fraction of expression of retrocopies (left panels) and parental genes (right panels) in GM12878. The expression levels of polyadenylated (ployA+; top panels) and non- polyadenylated (ployA-; bottom panels) RNAs are plotted separately. (*B*) Comparison of GRO-seq signal (left panel) and Pol II TSA-seq signal (right panel) between colocalized and noncolocalized retrocopies/parental genes in K562. (*C*) Comparison of the number of shared TFBSs between colocalized and noncolocalized retrocopy-parent pairs. \*\*\**p* < 0.001; two tailed Wilcoxon test.

Enhancer sharing has been hypothesized to account for the spatial colocalization and coregulation of paralogous genes located on the same chromosome (Ibn-Salem et al. 2017). Also, transcription factors (TFs) play a part in interchromosomal interactions (Kim and Shendure 2019). We thus interrogated the relationship between shared regulatory elements and spatial colocalization. As expected, we found that the flanking regions (putative transcription start site, i.e., TSS ± 3 kb) of colocalized retrocopies and parental genes harbor more transcription factor binding sites (TFBSs) than noncolocalized ones in most cell lines (*p* < 0.05; **supplementary fig. S14**). In particular, the number of shared TFBSs between colocalized retrocopies and parental genes is significantly greater than that of noncolocalized retrocopy- parent pairs (*p* < 0.001; **fig. 3C**). It should be noted that the choice of flaking regions (e.g., TSS ± 1 kb or upstream 3 kb) does not affect our results (**supplementary fig. S15**). Significantly, colocalized retrocopies show a higher correlation coefficient of expression with their parental genes across 17,282 GTEx samples in comparison with noncolocalized pairs (**supplementary fig. S16**), supporting the involvement of shared TFBSs in retrocopy-parent coexpression. Interestingly, the corresponding TFs of shared motifs are mostly zinc finger proteins, such as ZNF263, ZNF384, ZNF460, and ZNF135, consistent with a recent report in which zinc finger factors are found to play a role in speckle-associated interchromosomal interactions (Joo et al. 2023).

Colocalized and noncolocalized retrocopies are enriched in different subcompartments, which are associated with distinct epigenomic features and may affect the expression of retrocopies. We then explicitly examined the signals of 12 epigenomic features (including DNase I sensitivity, the histone variant H2A.Z, and 10 histone modifications) for retrocopies. Generally, colocalized retrocopies and their flanking regions have stronger signals of activating histone modifications (e.g., H3K4me1, H3K27ac, H3K36me3, and H3K79me2) in comparison with noncolocalized retrocopies (**fig. 4** and **supplementary figs. S17-S21**), indicating an association between epigenetic landscape and interchromosomal colocalization. Notably, retrocopies, even those noncolocalized ones, harbor stronger activating histone modifications than unprocessed pseudogenes (**fig. 4** and **supplementary figs. S17-S21**), further supporting the delimitation between processed and unprocessed “pseudogenes” in the context of 3D genome environment.

**Figure 4.**
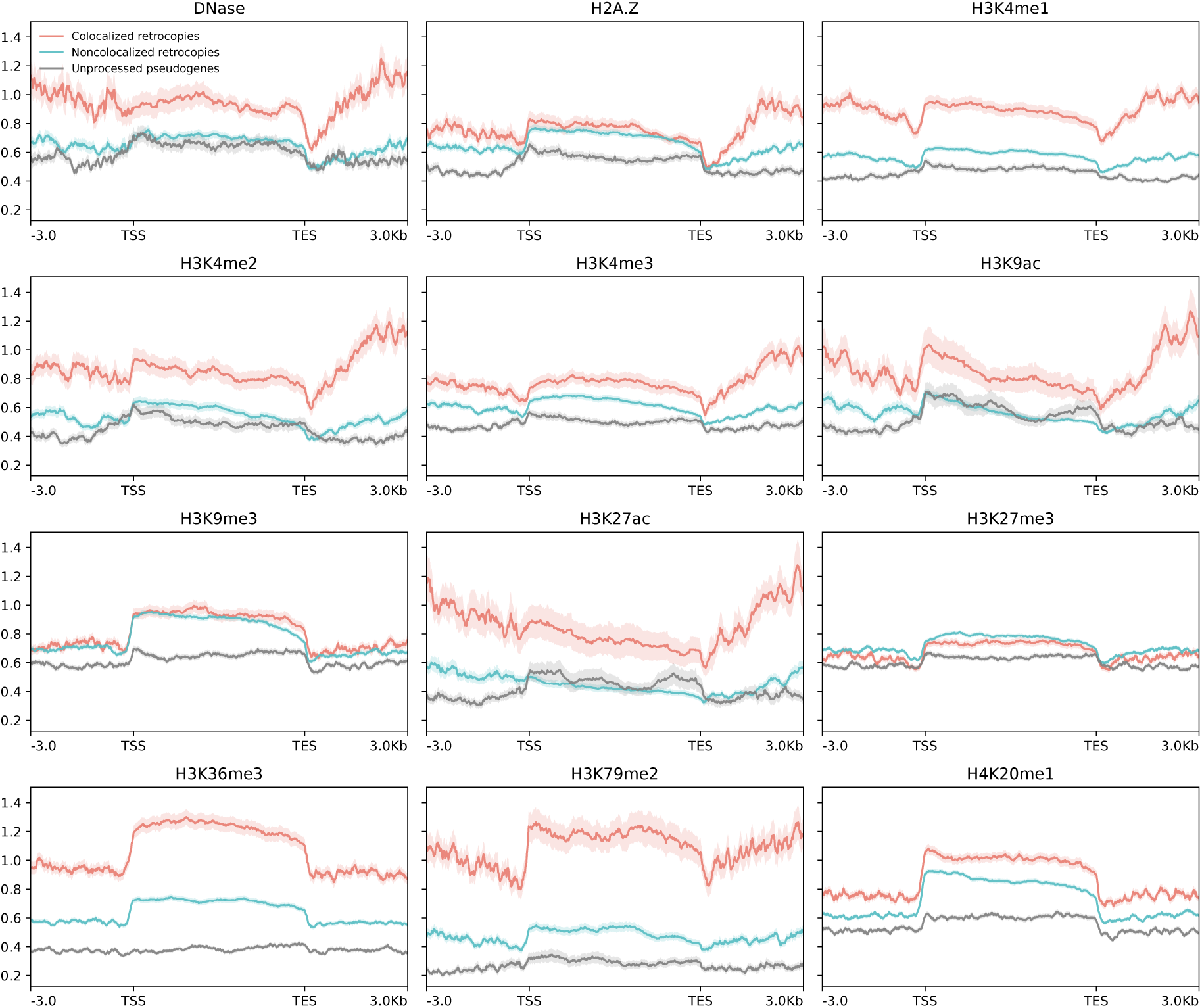
Epigenetic signals for DNase I hypersensitivity, histone variant H2A.Z, and 10 histone modifications along colocalized retrocopies (red lines), noncolocalized retrocopies (blue lines), and unprocessed pseudogenes (gray lines) and their flanking regions (± 3 kb) in the GM12878 cell line. Solid lines represent the mean fold change signal, shadowed areas denote the standard error. TSS: putative transcription start site; TES: putative transcription end site.

To evaluate the relative importance of different features in the colocalization between retrocopies and their parental genes, we trained a logistic regression model using 35 genetic and epigenetic features of retrocopies and parental genes as predictors (**supplementary table S2**) in each cell line where matched data were available. For GM12878, the machine learning model is able to predict retrocopy-parent colocalization with an accuracy of 0.798 and an area under the receiver operating characteristic curve (AUC) of 0.785 (**supplementary fig. S22A**). Among the features included, the number of shared motifs between retrocopies and parental genes is the most important, with various histone modifications of retrocopies also contribute significantly (**supplementary fig. S22B**). The number of shared motifs and epigenetic features of retrocopies are also top predictors in models of HUVEC, K562, and NHEK, but the performance is worse for these cell lines (**supplementary figs. S23-25**).

### Colocalized Retrocopies Are More Likely to Be Functional and Evolutionarily Conserved

Retrocopies may preserved the coding potentials of their parental genes before degenerative mutations disrupting the open reading frames (ORFs), therefore, gaining expression is the first and most important step of being functional for a retrocopy. The resulting RNAs may serve as functional units *per se*, such as NATs, lncRNAs, or microRNAs (Kubiak and Makałowska 2017). There are also sporadic cases that retrocopies act as important protein-coding genes in development and disease (Casola and Betran 2017). To further assess the functional relevance of retrocopies at the translation level, we leveraged the ribosome profiling (i.e., ribo-seq) data across 101 studies curated on RPFdb (Wang et al. 2019), and found that colocalized retrocopies have significantly higher RPKM (Reads Per Kilobase per Million mapped reads) values than noncolocalized retrocopies in five of the seven cell lines (*p* < 0.001; **fig. S26**). The difference in ribo-seq signals is statistically marginal between retrocopies that are colocalized in zero and one cell line (*p* = 7.8 × 10^-2^), while retrocopies with recurrent colocalized in two or more cell lines are covered by significantly more reads than the other two categories of retrocopies (*p* = 5.5 × 10^-13^ and 3.7 × 10^-6^, respectively; **fig. 5A**). Because ribo-seq reads could be derived from genomic fragments randomly co-purified with ribosomes, we further screened retrocopy transcripts with actively translated ORFs. These ORFs were detected using RibORF (Ji et al. 2015), which makes use of the 3-nt periodicity and uniformity of read distribution across codons of translated regions, thus reducing spurious signals of translation. When only transcripts with at least one detected ORFs were considered, colocalized retrocopies still present higher ribo-seq signals than noncolocalized ones (**supplementary fig. S27**). If the higher levels in transcription and translation of colocalized retrocopies did confer any functional significance, they would be more evolutionarily conserved than noncolocalized ones. As expected, we observed that the average phastCons conservation scores of colocalized retrocopies are significantly higher than noncolocalized ones (**fig 5B** and **supplementary fig. S28**). This pattern is retained when we use LINSIGHT scores (Huang et al. 2017) to measure the conservation of retrocopies (**supplementary fig. S29**). In addition, colocalized retrocopies and their putative promoters (1 kb upstream of TSSs) are more likely to overlap with NHGRI-EBI GWAS single nucleotide polymorphisms (SNPs) (Buniello et al. 2019) than noncolocalized ones (*p* < 0.01 in six out of the seven cell lines, Fisher exact test; **fig. 5C**), further supporting their higher potential of functionality. We found that the phastCons scores of retrocopies are positively correlated with their levels of RNA-seq (R = 0.45, *p* < 2.2 × 10^−16^) and ribo-seq (R = 0.28, *p* < 2.2 × 10^−16^; **fig. 5D**). However, after accounting for the variation of RNA expression level, the partial correlation coefficient between phastCons score and ribo-seq signal is only 0.088 (*p* = 5.5 × 10^-8^), while the partial correlation coefficient between phastCons score and RNA expression level is 0.374 (*p* = 1.8 × 10^-12^) when controlling for ribo-seq signal, suggesting that retrocopies may be of more functional relevance at the RNA level. Of particular note is that for two sister retrocopies derived from the same parental gene, the colocalized one generally has higher levels of expression, motif sharedness, and conservation than the noncolocalized one in GM12878 (*p* = 1.4 × 10^-2^, 1.5 × 10^-5^, and 3.0 × 10^-2^, respectively; **supplementary fig. S30**). But the trends are not apparent in other cell lines (data not shown), possibly due to the small sample size of desired retrocopies and the nosier nature of colocalization in cell lines other than GM12878.

**Figure 5.**
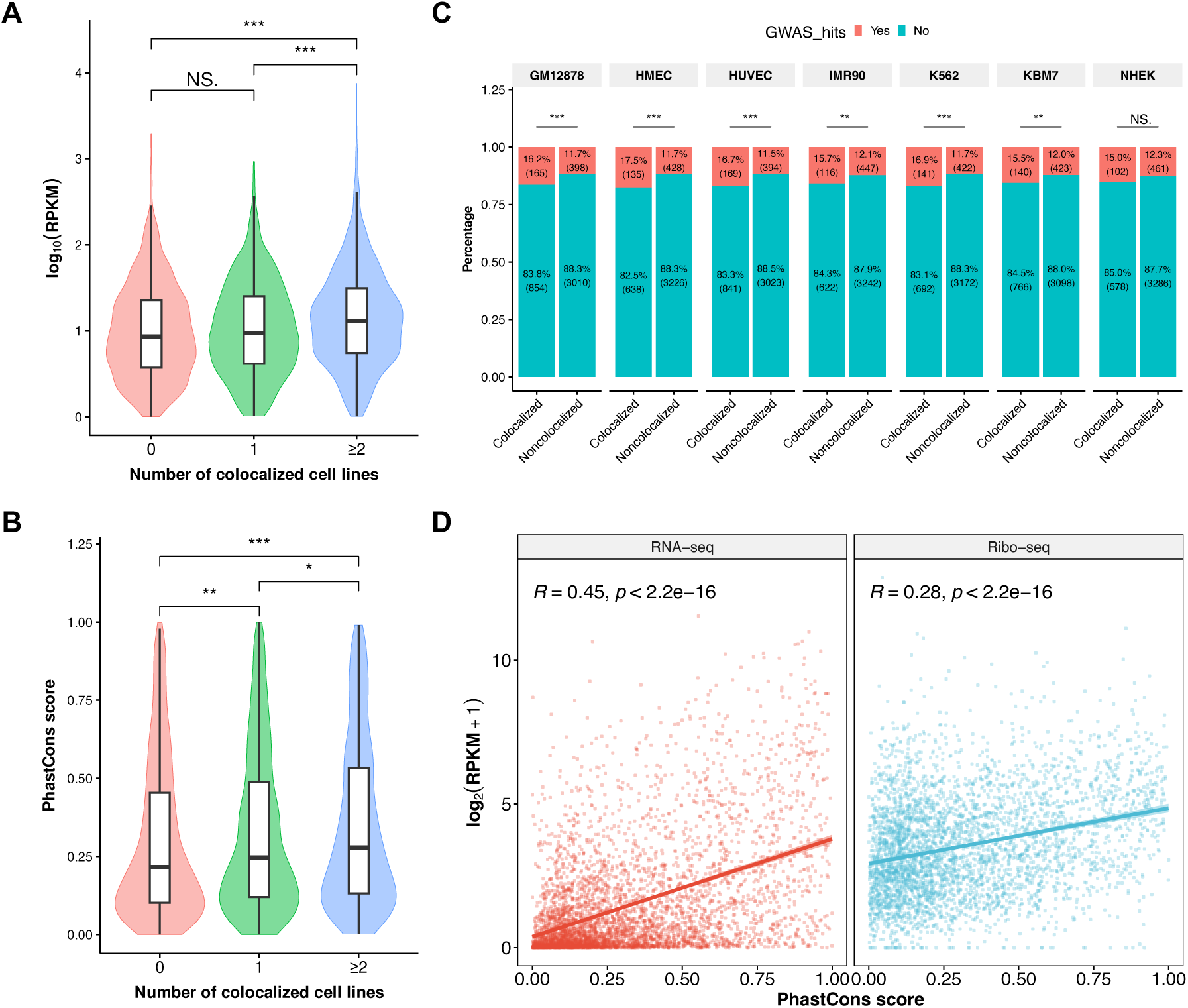
Coding potential and functional importance of colocalized and noncolocalized retrocopies. (*A-B*) Comparison of ribo-seq RPKM values (*A*) and phastCons scores (*B*) among retrocopies that are colocalized in 0, 1, and ≥ 2 cell lines. For ribo-seq data, only retrocopies with RPKM ≥ 1 were included. (*C*) Percentage of colocalized and noncolocalized retrocopies in the seven cell lines that are overlapped with at least one NHGRI-EBI GWAS SNPs. (*D*) Correlation between phastCons score and RNA expression level (left panel) and between phastCons score and ribo-seq signal (right panel) of retrocopies. Solid line and shadowed area in each plot denote the linear regression line and its confident interval. \*\*\**p* < 0.001, \*\**p* < 0.01, \**p* < 0.05, NS. not significant; two tailed Wilcoxon test in A and B, Fisher’s exact test in C.

In addition, to ask whether the colocalization pattern of retrocopy-parent pairs and its functional significance is conserved across mammalian species, we performed similar analyses with retrocopy-parent pairs of mice, chimpanzees, rhesus macaques, marmosets, cows, and dogs (**supplementary table S3**) (Carelli et al. 2016). Using Hi-C maps of corresponding species at 250-kb resolution (Rao et al. 2014; Zhang et al. 2019; Luo et al. 2021; Li et al. 2022), we found that retrocopy-parent pairs in five out of these six species exhibit significantly higher colocalization frequencies than randomly chromatin pairs under various null models (**supplementary fig. S31**). The result in dogs is an exception (*p* = 0.132 for simulation 5), possibly due to the large number and short length of dog’s chromosomes (Li et al. 2022). In mouse embryonic stem cells (mESCs), we found that colocalized retrocopy-parent pairs determined by Hi-C (Bonev et al. 2017) and SPRITE (Quinodoz et al. 2018) have a great proportion of overlap (**supplementary fig. S32A**). Leveraging high-resolution genome-wide seqFISH+ imaging data in mESCs (Takei et al. 2023), we also observed a significant shorter mean spatial distance for retrocopy-parent pairs identified as colocalized by Hi-C in comparison with noncolocalized pairs (*p* = 6.1 × 10^-94^, **supplementary fig. S33A**). More importantly, when significant interactions (*q* < 0.05) identified by SPRITE or the top 10% interchromosomal chromatin pairs with the shortest spatial distances detected by seqFISH+ were used to determine interchromosomal colocalization in mESCs, we found a higher-than-expected colocalization frequency for true retrocopy-parent pairs (**supplementary figs. S32B and S33B**).

In addition, we observed that colocalized retrocopies and parental genes also expressed at a higher level than noncolocalized ones in mouse CH12-LX cells (**supplementary fig. S34**). Comparison of ribo-seq signals and phastCons scores indicate mouse colocalized retrocopies are more actively translated and tend to be more conserved than noncolocalized ones (*p* = 0.002 and 0.012, respectively; **supplementary fig. S35**). Besides, the conservation of retrocopies is positively correlated with their levels of transcription and translation in mouse, with a greater correlation coefficient between conservation and RNA expression (**supplementary fig. S36**).

### Biased Insertion and Natural Selection Shaped the Interchromosomal Colocalization Between Retrocopies and Parental Genes

Lastly, we explored what molecular and evolutionary mechanisms might have shaped the spatial colocalization between retrocopies and parental genes. To this end, we identified 625 polymorphic retrocopies (retroCNVs) in five human populations (CEU, CHS, LWK, PEL, and YRI; 491 individuals) using sideRETRO (Miller et al. 2021) (**supplementary table S4**; Materials and Methods). Similar to Zhang et al. (2021), we did not distinguish homozygotes and heterozygotes, only accounted for the presence or absence of each allele in different individuals. Our list captures most retroCNVs identified in other studies (Abyzov et al. 2013; Ewing et al. 2013; Schrider et al. 2013; Zhang et al. 2017; Feng and Li 2021; Batcher et al. 2022) (**supplementary table S5**) and the neighbor-joining tree based on retroCNV presence frequency is topologically the same as the tree built from genome-wide SNPs (**supplementary fig. S37**), indicating the overall accuracy of retroCNV identification. The presence frequency spectrum of retroCNVs is more skewed toward low-frequency in comparison with SNPs (**supplementary fig. S38**), congruent with the highly deleterious effect of retroCNV insertions. Ninety-eight retroCNVs are present in only one individual, which may be enriched in *de novo* retrocopy insertions. We then performed the same simulations for these presumably *de novo* retrocopies and their parental genes as did for putative “fixed” retrocopy-parent pairs (**supplementary fig. S2**), and found that the colocalization frequency between *de novo* retrocopy insertion sites and their parental genes is only significantly higher than expected in some simulations (**fig. 6A** and **supplementary fig. S39**). This result suggests that the insertion of retrocopies is not random, but nonrandom insertion cannot fully explain the current colocalization pattern between retrocopies and parental genes.

**Figure 6.**
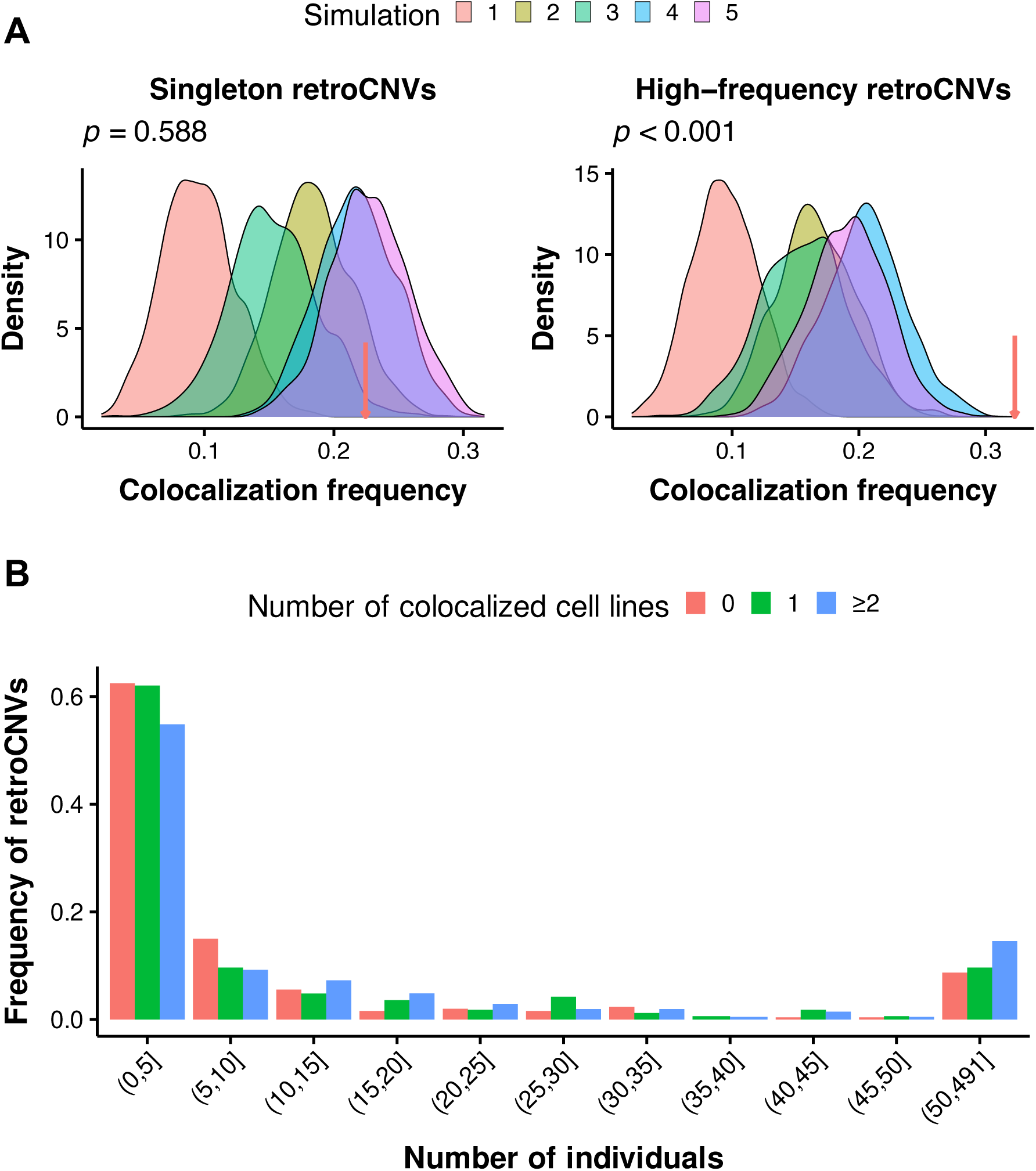
Colocalization and evolution of retroCNVs in humans. (*A*) Density plots showing the distribution of colocalization frequency between chromatin pairs under different null models for singleton retroCNVs (left panel) and high-frequency retroCNVs (right panel). Red arrows denote the colocalization frequency between true retroCNVs and their parental genes. Simulations and true frequencies are based on Hi-C data in GM12878. *P*-values are indicated for simulation 5. (*B*) The presence frequency spectra for retroCNVs that are colocalized in 0, 1, and ≥ 2 cell lines.

Then we investigated the potential role of post-insertion selection in determining retrocopy- parent colocalization. In contrast to *de novo* retrocopy insertions, high-frequency (occurs in ≥ 28 individuals; n = 96, a sample size that is comparable to singletons) retroCNVs and their parental genes exhibit a higher-than-expected colocalization frequency in all simulations (*p* < 0.01 except for simulations 4 and 5 in HUVEC. **fig. 6A** and **supplementary fig. S40**), indicating the action of post-insertion selection on retrocopy survival in the context of interchromosomal interactions. Furthermore, we found that the presence frequency spectrum of retroCNVs exhibiting colocalization in two or more cell lines is less skewed toward low frequency compared to retroCNVs showing colocalization in zero or one cell line (**fig. 6B**), suggesting relaxed negative selection on colocalized retroCNVs. The difference in frequency spectrum holds true for retroCNV subclasses base on colocalization status in individuals cell lines (**supplementary fig. S41**). With respect to the 3D genome organization, recurrently colocalized retroCNVs are enriched in the active A1 subcompartment (**fig. 2A** and **supplementary fig. S4**) and interior active regions and speckles (**supplementary fig. S6**). These active chromatin regions may have higher density of selected sites and/or more intense selection, which are negatively correlated with local effective population size (Gossmann et al. 2011), leading to reduced selection efficacy on recurrently colocalized retroCNVs. Indeed, the density of conserved sites (phastCons score > 0.5) flanking colocalized retroCNVs (insertion site ± 50 kb) is significantly higher than noncolocalized retroCNVs (**supplementary fig. S42**). Also, the unfolded site frequency spectrum of SNPs in the flanking regions (insertion site ± 50 kb) of recurrently colocalized retroCNVs is more skewed toward low frequency (**supplementary fig. S43**).

Because there is also a higher proportion of intermediate-to-high-frequency alleles for recurrently colocalized retroCNVs, we sought to detect whether the elevated frequency of colocalized retroCNVs is driven by positive selection. If a retroCNV insertion was under strong positive selection, selective sweep would: 1) reduce the nucleotide diversity (*π*) among sequences near the insertion for retroCNV carriers compared to noncarriers; 2) reduce the Tajima’s D among sequences near the insertion for retroCNV carriers compared to genomic background; 3) increase the linkage disequilibrium (LD) between SNPs near the insertion for retroCNV carriers compared to genomic background (Zhang and Tautz 2022). Based on these principles, we scanned signals of positive selection for each retroCNV with allele frequency ≥ 0.5 in individual populations using SNPs from the 1000 Genomes Project (Altshuler et al. 2015) (Materials and Methods). We found that neither colocalized nor noncolocalized high-frequency retroCNVs are under positive selection using the above criteria (**supplementary table S6**). Therefore, our results indicate that the intermediate-to-high-frequency of both colocalized and noncolocalized retroCNVs is probably the result of nonadaptive process or weak positive selection that left untraceable signal.

## Discussion

In the present study, we revealed that retrocopy-parent pairs exhibit higher frequency of interchromosomal colocalization than expected under various null models and that colocalized retrocopies are more active in terms of transcription and translation. We further explored the potential mechanisms, functional consequences, and evolutionary implications of the nonrandom colocalization between retrocopies and their parental genes.

We focused exclusively on interactions between retrocopies and parental genes that are located on different chromosomes, i.e., interchromosomal contacts. Compared to intrachromosomal structures (e.g., TADs and chromatin loops), the mechanisms and functional consequences of interchromosomal interactions are less well understood. Although often being thought as noise or technical artifacts, emerging evidence suggests that interchromosomal interactions are relevant to fundamental cellular activity, such as transcription and splicing (Maass et al. 2019). Using CRISPR/Cas9 live-cell imaging, Maass et al. (2018) showed that interchromosomal contacts are as stable and frequent as intrachromosomal interactions. In another study, Quinodoz et al. (2018) detected two hubs of interchromosomal interactions that shape 3D genome organization in the nucleus using SPRITE. Along with accumulating evidences of *trans* gene regulation in health and in disease (Patel et al. 2014; Bashkirova and Lomvardas 2019; Maass et al. 2019; Lei et al. 2023; Moon et al. 2023), recent progress indicates that interchromosomal interactions play structural and functional roles in cells. Our finding that retrocopy function is linked to *trans* interactions with parental genes also hints nonrandom organization of interchromosomal contacts.

Interchromosomal contacts often occur at farther spatial distances than *cis* interactions (Maass et al. 2018), rendering their detection difficult for regular ligation-based methods (e.g., Hi-C). In this study, our main analyses of interchromosomal interactions rely on Hi-C contact maps at 250-kb resolution, given the constraints of sequencing depth. Leveraging the ultra-high-depth Hi-C data in GM12878, we further detected spatial colocalization between retrocopies and parental genes at 100-kb and 500kb resolutions. Under the same criterion (*q*- value < 0.05 in FitHiC2), the number of colocalized pairs of retrocopy-parent pairs decreases with the increase of resolution. Nevertheless, the colocalization pairs based on high-resolution data is almost a subset of pairs detected using low-resolution contact maps (**supplementary fig. S44**). Furthermore, retrocopies and their parental genes always possess a higher-than-expected colocalization frequencies at different resolutions (**supplementary fig. S45**). More importantly, chromatin contact maps derived from orthogonal methods (SPRITE, MERFISH, and seqFISH+) in human and mouse also support the significant colocalization between retrocopy-parent pairs. However, a limitation in our assessment of interchromosomal interactions is the low resolution of certain contact maps and the scarcity of cell lines with complementary data simultaneously available.

Subsequently, we established that colocalized and noncolocalized retrocopies exhibit distinct properties in terms of subcompartment annotations and cytological distance to subnuclear structures (**fig. 2 and supplementary figs. S4-7**). For subcompartments, both A1 and A2 harbor highly expressed genes and activating chromatin marks, and are depleted in nucleolus-associated domains (NADs), but they show difference in replication timing and GC content. B1 represents facultative heterochromatin that exhibits intermediate level of transcriptional activity and high level of H3K27me3. B2 and B3 are associated with nuclear lamina, but B2 is enriched in NADs whereas B3 is strongly depleted in NADs (Rao et al. 2014; Xiong and Ma 2019). Using TSA-seq data, Chen et al. (2018) further revealed that chromatin in A1 subcompartment is close to nuclear speckles while A2 subcompartment corresponds to region with non-speckle associated transcription. Therefore, our observations (**fig. 2 and supplementary figs. S4-7**) imply that many interchromosomal interactions between retrocopies and their parental genes may be mediated by nuclear speckles. Nuclear speckles are domains enriched in pre-mRNA splicing factors (Spector and Lamond 2011) and are also hubs of interchromosomal interactions where active transcription takes place nearby (Galganski et al. 2017; Quinodoz et al. 2018; Chen and Belmont 2019; Kim et al. 2020).

Remarkably, zinc finger proteins are found to facilitate the establishment of speckle-associated interchromosomal contacts (Joo et al. 2023), in accordance with our findings that the number of shared regulatory motifs is the most importance feature in predicting retrocopy-parent colocalization and that most shared motifs are bind by TFs in the zinc finger family. Thus, our findings further support a role of TFs in interchromosomal interactions (Kim et al. 2019; Kim and Shendure 2019).

Speckle-associated interchromosomal interactions, along with more active chromatin marks and higher number of shared motifs associated with colocalized retrocopies, signify colocalized retrocopies should be more actively transcribed. The implications of higher expression of colocalized retrocopies could be twofold. First, retrocopy-derived RNAs could function as NATs, lncRNAs, or microRNAs in diverse cellular activities, including regulation of their parental genes (Kubiak and Makałowska 2017). Besides, RNAs transcribed from retrocopies may be processed and translated, and function at the protein level (Qian et al. 2022). Second, the action of transcription and the resulting RNAs can orchestrate both intra- and inter-chromosomal chromatin organization (van Steensel and Furlong 2019; Bertero 2021; Quinodoz and Guttman 2021). In the nucleus, 3D genome organization and transcription are intermingled, with reciprocal influence on each other. Recent single-cell Hi-C and RNA-seq data discovered widespread chromatin interaction rewiring before transcription activation, strongly supporting a regulatory role of chromatin organization on gene expression (Liu et al. 2023). Meanwhile, TFs can bring DNA fragments on the same or different chromosomes to spatial proximity through direct oligomerization or cofactor oligomerization, leading to transcription-mediated interactions in *cis* or *trans* (Kim et al. 2019; Kim and Shendure 2019). In addition, non-coding RNAs in the nucleus may serve as chromosome scaffolds or seeds to guide chromatin organization (Mele and Rinn 2016; Quinodoz et al. 2021; Bouwman et al. 2022). However, the actual impact of retrocopy transcription and retrocopy-derived RNAs on interchromosomal interactions needs additional investigations.

We further showed that both biased insertion and natural selection contribute to the nonrandom colocalization between retrocopies and parental genes (**fig. 6**). Mechanistically, new L1 insertions are linked to DNA replication, with a preference in early-replicating regions of the genome (Flasch et al. 2019; Sultana et al. 2019). Because early-replicating regions are enriched with active subcompartments and speckles and are more like to be involved in interchromosomal interactions (Vouzas and Gilbert 2021), if retrocopy insertions have the same property as L1s, this insertion bias would partly explain the higher-than-expected colocalization frequency between *de novo* retroCNVs and parental genes.

Our population genetic analysis indicates that colocalized retroCNVs are more likely to reach high frequency, but this elevated frequency is probably driven by nonadaptive forces. Then, what is the evolutionary significance of retrocopy-parent colocalization? Given the fact that colocalized “fixed” retrocopies are more actively transcribed and have higher potential of functionality than noncolocalized ones, we argue that the spatial proximity with parental genes may facilitate the “resurrection” of initially neutral retrocopies. This is similar to the situation where new genes frequently originated in the promiscuous testis (Kaessmann 2010). Colocalized retrocopies are also residing in chromatin with activating epigenetic marks, are cytologically close to speckles, and extensively share regulatory elements with their progenitors, all of which may facilitate their exploitation of cellular transcription machinery and the exaptation of these seemingly neutral retroposed gene copies.

However, we are aware of some potential caveats should be considered in interpreting the population genetic analysis of retroCNVs. First, our search for retroCNVs is not exhaustive. Only five representative human populations were used to identify retroCNVs, and only 98 singletons were identified to represent potential *de novo* retroCNVs, which may reduce the statistical power of our analysis. Second, our approach ignored retroCNVs that are present in the reference genomes, but absent in some other individuals. However, such retroCNVs may represent a relatively small proportion of total retroduplication variants (Zhang et al. 2017). Third, we were unable to derive the ancestral state of retroCNVs. Some low-frequency retroCNVs may arise from losses of ancestral retrocopy insertions. Nevertheless, our investigation exemplifies a first yet valuable attempt to understand the evolutionary dynamics of retrocopies in the context of 3D genome organization. In the future, more thorough survey of retroCNVs at large scale, coupled with long-read sequencing (e.g., Feng and Li 2021; Troskie et al. 2021), should give us a deeper understanding of the link between 3D genome organization and the function and evolution of retrocopies.

In conclusion, we demonstrated that retrocopies and their parental genes displayed significant colocalization in the nucleus, and characterized the spatial positioning and functional profiling of colocalized and noncolocalized retrocopies in great detail. Our results highlight the importance of 3D genome organization in the function and evolution of stripped-down retrocopies.

## Materials and Methods

### Retrocopy-Parent Pair Datasets

The lists of retrocopies and their cognate parental genes of human (GRCh37), mouse (NCBIM37), chimpanzee (CHIMP2.1.4), rhesus macaque (MMUL_1) were retrieved from the Supplementary Table S1 of Carelli et al. (2016). The retrocopy information of marmoset (C_jacchus3.2.1), cow (UMD3.1), and dog (CanFam3.1) were downloaded from RetroGeneDB2 (Rosikiewicz et al. 2017). Retrocopy-parent pairs fit any of the following criteria were removed from further analyses: 1) a retrocopy or its parental gene was not located on autosomes or the X chromosome; 2) a retrocopy and its parental gene were located on the same chromosome; 3) the information of parental gene of a retrocopy was inferred from other species (Carelli et al. 2016).

### Identification of Colocalized Retrocopy-Parent Pairs

The raw interchromosomal contact matrices at 250-kb resolution for seven human cell lines and one mouse cell line were downloaded from NCBI (GEO accession: GSE63525) (Rao et al. 2014). These contact matrices were constructed using Hi-C read pairs that were uniquely mapped to the respective reference genomes (mapping quality ≥ 30). The contact matrices in hic format of mESC (4DNESDXUWBD9; mm10), chimpanzee (panTro6) and marmoset (calJac3) induced Pluripotent Stem Cell (iPSC) (GSE116862), and macaque cortex plate (GSE163177; rheMac8) were downloaded from 4DN portal or NCBI and were converted to FitHiC2 format using HiCExplorer v3.7.2 (Wolff et al. 2020). For cow and dog, we downloaded the raw reads of ear skin Hi-C sequencing from NCBI (GSE167581) and constructed contact matrices using HiC-Pro v3.1.0 (Servant et al. 2015). Only contacts that were supported by high-mapping-quality read pairs (mapping quality ≥ 30) were retained for downstream analyses.

Significant interchromosomal interactions at 250-kb were identified using FitHiC2 v2.0.7 (Ay et al. 2014; Kaul et al. 2020). The Knight-Ruiz matrix balancing algorithm (Knight and Ruiz 2013) was applied to normalized Hi-C data during identification. Unlike intrachromosomal interactions where the observed contacts can be compared with the null distribution of contacts based on the linear distances between fragments to determine statistical significance, *p*-values of interchromosomal interactions in FitHiC2 were calculated using a uniform probability model (Duan et al. 2010). At the level of *q*-value (Benjamini-Hochberg adjusted *p*-value) < 0.05, 5,152,588 significant interchromosomal interactions at 250-kb were identified. Due to the much shallow depth of Hi-C sequencing in the other cell lines and species, far fewer significant interactions at *q*-value < 0.05 were detected. In order to perform comparable analyses, we selected the top-5 million fragment pairs with lowest *p*-values as significant interactions for all samples, as suggested by the authors of FitHiC2 (Kaul et al. 2020).

The colocalization status between retrocopies and their parental genes were determined from the interaction of their residing genomic fragments using the findOverlaps function of the R package GenomicInteractions v1.30.0 (Harmston et al. 2015). We converted the genomic coordinates of retrocopies to that of Hi-C matrices using the UCSC liftover tool when there were genome assembly mismatches. If a retrocopy and its parental gene were overlapped respectively with two ends of an interchromosomal fragment pair that was identified as significant interaction, then they were denoted as “colocalized”, otherwise they were “noncolocalized”. Visualization of significant interaction between retrocopies and parental genes was plotted with the R package circlize v0.4.14 (Gu et al. 2014).

### Statistical Tests on Chromatin Spatial Colocalization of Retrocopy-Parent Pairs

To assess the statistical significance of interchromosomal interactions between retrocopies and parental genes, we performed five different simulations. 1) Random fragment pairs with matching chromosomes and sequence lengths as the retrocopy-parent pairs were generated to represent simulated chromatin pairs. 2) The information of retrocopies was kept unchanged while sampling random genes (including coding and noncoding ones) with matching chromosomes as parental genes to form simulated chromatin pairs. 3) The information of parental genes was kept unchanged while sampling random fragments with matching chromosomes and sequence lengths as retrocopies to be simulated chromatin pairs. 4) Similar to the second simulation, but the randomly sampled genes were restricted to protein-coding genes only. 5) The pairing relationship between retrocopies and parental genes was randomly shuffled. For each sampling, the same number of chromatin pairs as true retrocopy-parent pairs was generated, and 1,000 samplings were carried out for each simulation. Colocalization of simulated chromatin pairs was determined using Hi-C data of respective cell lines, and the *p*- value was calculated as the fraction of simulated data sets that exhibited a colocalization frequency higher than that of true retrocopy-parent pairs. See also **supplementary fig. S2** for graphic illustrations.

### Analysis of SPRITE Data

The SPRITE interchromosomal interaction matrices of GM12878 and mESC were downloaded from NCBI (GSE114242). Similar to the original paper (Quinodoz et al. 2018), we applied FitHiC2 v2.0.7 (Ay et al. 2014; Kaul et al. 2020) to detect significant interactions using contact maps binned at 1Mb resolution based on SPRITE clusters containing 2 to 1000 reads without down-weighting for cluster size.

### Analysis of MERFISH and Two-layer seqFISH+ Imaging-based Data

The physical coordinates (x, y, and z) of 1,041 uniformly distributed genomic loci detected by MERFISH in IMR90 cells (Su et al. 2020) were downloaded from Zenodo (doi: 10.5281/zenodo.3928890). The spatial distance between any pair of loci was calculated as the Euclidean distance between their fitted 3D Gaussian centers using the scripts provided by the authors (https://github.com/ZhuangLab/Chromatin_ Analysis_2020_cell). The median distance of a given pair of loci was calculated from ∼5400 cells where both ends were detected. The proximity frequency between two loci was computed as the number of cells in which their spatial distance was less than 1 μm divided by the total number of detected cells. The coordinates of DNA spots were extended to 1.5 Mb before overlapping with retrocopies and parental genes.

For two-layer seqFISH+ data in mESC (Takei et al. 2023), we downloaded the physical coordinates (x, y, and z) of 100,049 genomic loci at 25-kb resolution in 1,076 cells from Zenodo (doi: 10.5281/zenodo.7693825). The spatial distance between any pair of loci binned at 200-kb resolution was calculated using the scripts provided by the authors (https://github.com/CaiGroup/dna-seqfish-plus-multi-omics). The mean distance between any retrocopy-parent pair was obtained by overlapping the mean distance matrix at 200-kb resolution calculated across detected cells.

### Genome-wide Enrichment with Subcompartments, TADs, and SPIN States

Genome-wide annotations of subcompartments in GM12878 and TADs in the seven human cell lines were downloaded from NCBI (GEO accession: GSE63525) (Rao et al. 2014). Subcompartment annotations for HMEC, HUVEC, IMR90, and K562 were retrieved from https://cmu.box.com/s/n4jh3utmitzl88264s8bzsfcjhqnhaa0. In cell lines other than GM12878, subcompartment annotation was achieved through the SNIPER computational approach, in which interchromosomal interactions were imputed and subcompartments were inferred by denoising autoencoder and multilayer perceptron classifier using typical Hi-C datasets with moderate depth (Xiong and Ma 2019).

The annotation of 10 spatial compartmentalization states for K562 was downloaded from https://github.com/ma-compbio/SPIN.

Enrichment or depletion of retrocopies, parental genes, unprocessed pseudogenes, and retroCNV insertion sites in subcompartments, TAD, and SPIN states were computed using the Genomic Association Tester (GAT) (Heger et al. 2013). Only mappable regions of the hg19 genome were included for the sampling of random regions to determine the enrichment or depletion of test genomic elements. Retrocopies and parental genes were divided into “colocalized” and “noncolocalized” ones based on Hi-C data of individual cell lines. To assess the impact of host genes, retrocopies were further divided into “intragenic” and “intergenic” subcategories, based on whether they overlap with protein-coding genes. Protein-coding genes and unprocessed pseudogenes (GENCODE release 39; lifted to GRCh37) were included for comparison. Note that “transcribed unprocessed pseudogenes” annotated by GENCODE were not included. For retroCNVs, we used the estimated coordinates of insertion sites for enrichment analysis.

### TSA-seq and Related Data

Genomic tracks of SON, pSC35, lamin B, lamin A/C, and pol II TSA-seq signals, GRO-seq signal, and speckle distance in bigwig format for the K562 cell line were downloaded from NCBI (GEO accession: GSE81553) (Chen et al. 2018). The signal strength in 20 kb non-overlapping bins were calculated using deepTools (Ramirez et al. 2016) and intersected with retrocopies and parental genes using bedtools (Quinlan and Hall 2010).

To assess the influence of speckle distance on colocalization, we carried out an additional simulation, in which the information of retrocopies were kept unchanged, while sampling random protein-coding genes with matching speckle distance (± 0.01 μm away) as true parental genes to form simulated chromatin pairs. The *p*-value was calculated as the number of occurrences where the colocalization frequency of simulated pairs is higher than true retrocopy-parent pairs over 1,000 samplings.

### RNA-seq, TFBS, and Histone Modification Analyses

The genomic alignment files in bam format of RNA-seq for GM12878, HUVEC, K562, and NHEK were downloaded from the ENCODE RNA Dashboard website (https://public-docs.crg.eu/rguigo/Data/jlagarde/encode_RNA_dashboard//hg19/). RNA-seq data of mouse CH12-LX cell line was downloaded from ENCODE (ENCSR000AJV). Only uniquely mapped reads were kept for downstream analysis. The FPKM values of retrocopies and other genes were estimated using StringTie v2.2.1 (Pertea et al. 2016) and are averaged across replicates.

The position frequency matrices (PFMs) of 949 human TFs were downloaded from the JASPAR2022 database (https://jaspar.genereg.net) (Castro-Mondragon et al. 2022), and FIMO v5.4.1 (Grant et al. 2011) was used to scan the occurrences of TFBSs in the flanking sequences (± 3 kb, ± 1 kb, or upstream 3 kb of putative TSSs) of retrocopies and parental genes. Only TFBSs with *q*-value < 0.01 were retained for further analysis. The TPM (Transcripts Per Million) values of human genes and transcripts across 17,382 samples were downloaded from GTEx portal (release V8).

Genome-wide fold change signals of DNase I sensitivity, histone variant H2A.Z and 10 histone modifications (H3K4me1, H3K4me2, H3K4me3, H3K9ac, H3K9me3, H3K27ac, H3K27me3, H3K36me3, H3K9me2, and H4K20me1) in bigwig format for GM12878, HMEC, HUVEC, IMR90, K562, and NHEK were obtained from the Roadmap Epigenomics Project (https://egg2.wustl.edu/roadmap/web_portal/). The signals of retrocopies and unprocessed pseudogenes and their flanking regions (± 3 kb) were computed and visualized with DeepTools v3.5.1 (Ramirez et al. 2016).

### Logistic Regression and Evaluation of Feature Importance

For GM12878, HUVEC, K562, and NHEK, we built a logistic regression model to predict the colocalization status of retrocopy-parent pairs using 35 genetic and epigenetic features related to retrocopies and parental genes as predictors (**supplementary table S2**). The model was trained using 70% randomly selected retrocopy-parent pairs, and the remaining 30% of data was used to evaluate the performance of trained model. The R package Boruta v7.0.0 (Kursa and Rudnicki 2010) was applied to assess the importance of individual features.

### Coding Potential and Evolutionary Importance of Retrocopies

The RPKM values and ORF information of human and mouse genes inferred from ribosome profiling data were downloaded from RPFdb v2.0 (Wang et al. 2019). For human and mouse, 101 and 52 ribo-seq studies were included in the RPFdb database. The maximum RPKM of a given gene among all ribo-seq samples was used to compare the coding potential between colocalized and noncolocalized retrocopies, similar to Qian et al. (2022). The ORFs were detected with RibORF (Ji et al. 2015), which combines the 3-nt periodicity and uniformity of read distribution across codons of translated regions to identify translated ORFs from ribo-seq data.

The human 46-way phastCons scores and the mouse 60-way phastCons scores were downloaded from UCSC Genome Browser. For both species, the phastCons score of the placental mammal subset was used. For human, we also downloaded the precomputed LINSIGHT score from http://compgen.cshl.edu/LINSIGHT/LINSIGHT.bw. LINSIGHT leverages both functional and population genomic data to detect deleterious noncoding variants. The average phastCons/LINSIGHT scores of retrocopies were computed with the bigWigAverageOverBed tool of UCSC.

A total of 516,945 GWAS SNPs associating with various diseases were downloaded from the National Human Genome Research Institute’s (NHGRI) GWAS catalog (Buniello et al. 2019). Duplicated recodes were removed, leaving 254,286 unique SNPs for further analysis.

### Identification and Population Genetic Analysis of retroCNVs

To investigate the insertion bias and evolutionary forces of newly inserted retrocopies in relation to interchromosomal interactions, we used sideRETRO v1.1.3 (Miller et al. 2021) to identify retroCNVs using whole genome sequencing (WGS) data from the 1000 Genomes Project. sideRETRO utilize abnormal alignments (i.e., discordantly aligned paired-end reads and split reads) in WGS or whole exome sequencing (WES) to identify unfixed retrocopies absent in the reference genome, but present in the sequenced individual. Considering the computation complexity, only the WGS mapping files of 491 individuals from five diverse populations (CEU, CHS, LWK, PEL, and YRI) were used. These individuals were sequenced to a high depth (30×), and were good resources for detecting retroCNVs using sideRETRO.

The initial list of retroCNVs was further filtered with the following criteria: 1) retroCNVs that were not located on autosomes and the X chromosome were removed. 2) If the insertion site of a retroCNV and its parental gene were on the same chromosome, they were excluded from further analysis. 3) retroCNVs with a missing rate greater than 20% were discarded.

To detect possible positive selection on retroCNVs, we calculated three population genetic statistics for each retroCNV that had a frequency ≥ 50% in any of the five populations: 1) the average nucleotide diversity (*π*) of 100 kb flanking regions centered at the insertion sites of retroCNV carriers over that of non-carriers (*π*_*ratio*_) (Schrider et al. 2013). 2) average Tajima’s D of 100 kb flanking regions centered at the insertion sites of retroCNV carriers (Llopart et al. 2002). 3) average LD (r^2^) of 10 kb flanking regions centered at the insertion sites of retroCNV carriers (Cardoso-Moreira et al. 2016). These statistics were computed for individual populations using matching polymorphism data from the 1000 Genomes Project. In order to evaluate the statistical significance of the estimates, we randomly generated “pseudo” insertion site in the genome, while keeping the same setup of carriers and noncarriers as the focal retroCNV. A total of 200 samplings were carried out and the Wilcoxon rank-sum test was used to determine how likely to get a smaller *π*_*ratio*_, a smaller Tajima’s D, or a greater LD from “pseudo” insertion sites as that from a true retroCNV insertion (Zhang and Tautz 2022).

## Data Availability

No new data were generated or analysed in support of this research. The custom written scripts used in this study are available at GitHub (https://github.com/yanyubin/Retrocopy_Evolution). The sources of public data and their corresponding genome assembly versions are summarized in supplementary table S7.

## Acknowledgements

We thank members of the Yang laboratory for their discussion. This work was supported by the Fund of Northwest A&F University (Z111021404) and the “100-Talent Program” of Shaanxi Province of China (A289021612) to R.Y.

## Author Contributions

R.Y. conceived and supervised the project. Y.Y. and R.Y. designed the study. Y.Y., Y.T., Z.W., and K. Z. performed data analysis. Y.Y. wrote the manuscript with input from R.Y. and Y.T. All authors reviewed and approved the manuscript.

## Supplementary Material

Supplementary tables S1-S7 and supplementary figures S1-S45 are available as separate files.

