## supplementary fig. for "Interchromosomal Colocalization with Parental Genes Is Linked to the Function and Evolution of Mammalian Retrocopies"

**Supplementary figures**

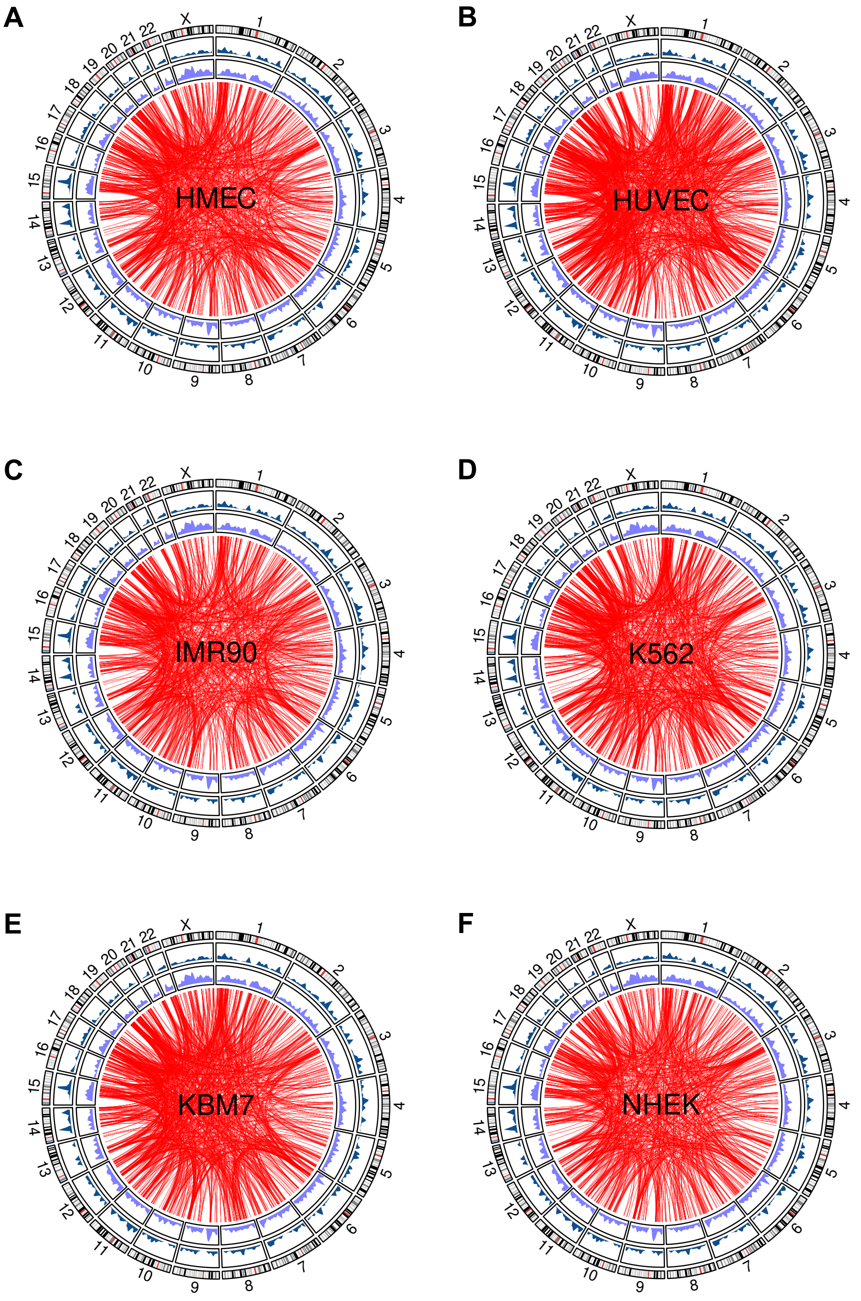


**Figure S1.** Circos plots highlighting the spatial colocalization between retrocopies and their parental genes in the HMEC (A), HUVEC (B), IMR90 (C), K562 (D), KBM7 (E), and NHEK (F) cell lines. From outer to inner are the ideogram of human chromosomes, the density of parental genes, the density of retrocopies, and significant chromatin interactions between retrocopies and parental genes.

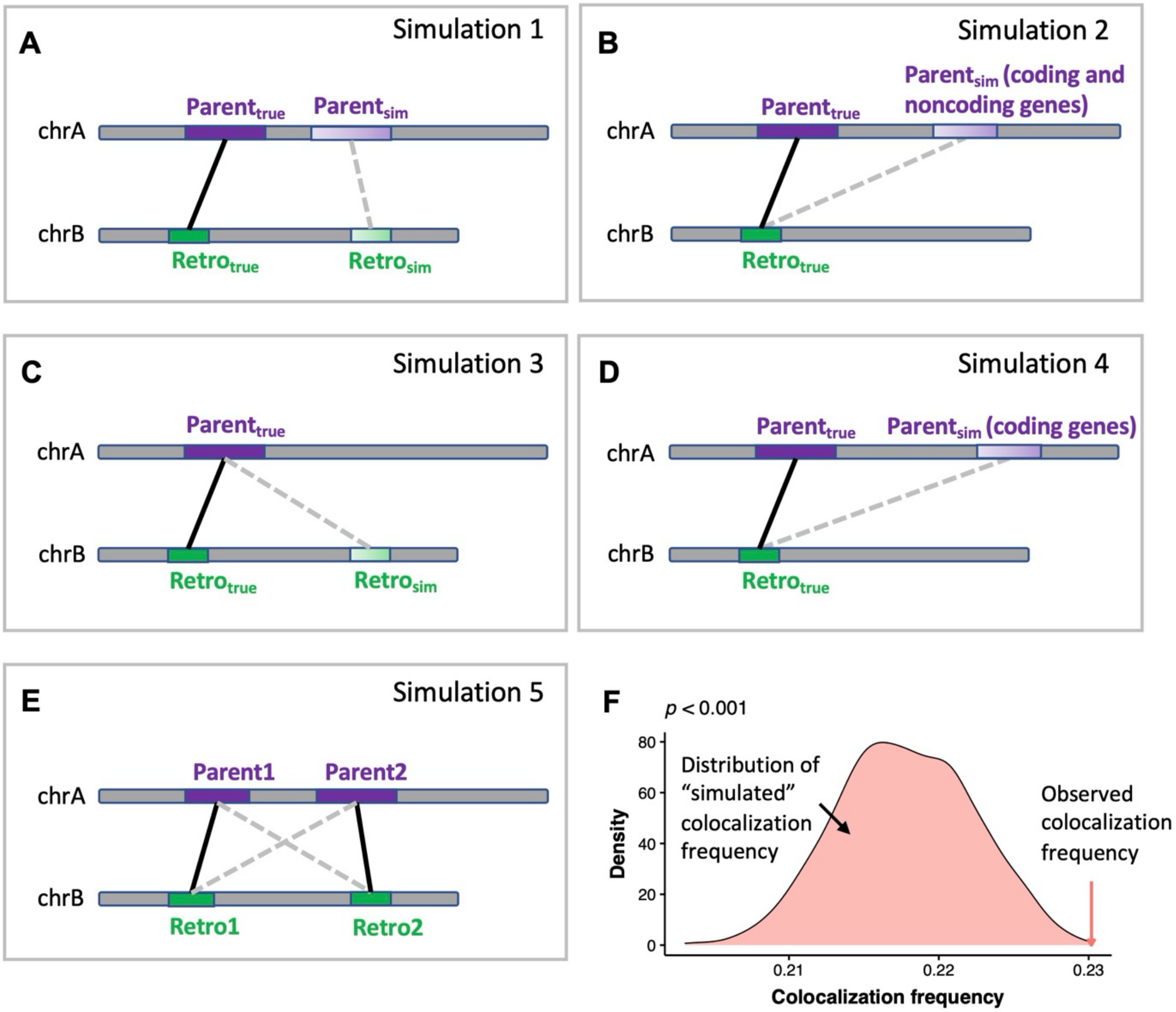


**Figure S2.** Schematic illustrations of the five simulations carried out to assess the significance of colocalization between retrocopies and parental genes. (A) Simulation 1: random fragment pairs with matching chromosomes and sequence lengths as the retrocopy-parent pairs were generated to represent simulated chromatin pairs. (B) Simulation 2: the information of retrocopies was kept unchanged while sampling random genes (including coding and noncoding ones) with matching chromosomes as parental genes to form simulated chromatin pairs. (C) Simulation 3: the information of parental genes was kept unchanged while sampling random fragments with matching chromosomes and sequence lengths as retrocopies to be simulated chromatin pairs. (D) Simulation 4: similar to the second simulation, but the randomly sampled genes were restricted to protein-coding genes only. (E) Simulation 5: the pairing relationship between retrocopies and parental genes was randomly shuffled. (F) For each simulation, 1000 random samplings with identical number of chromatin pairs as true retrocopy-parent pairs were carried out, and the distribution of simulated colocalization frequency was determined using Hi-C data of matching cell lines. The *p*-value was calculated as the fraction of simulated data sets that exhibited an equal or higher colocalization than that between true retrocopies and their parental genes.

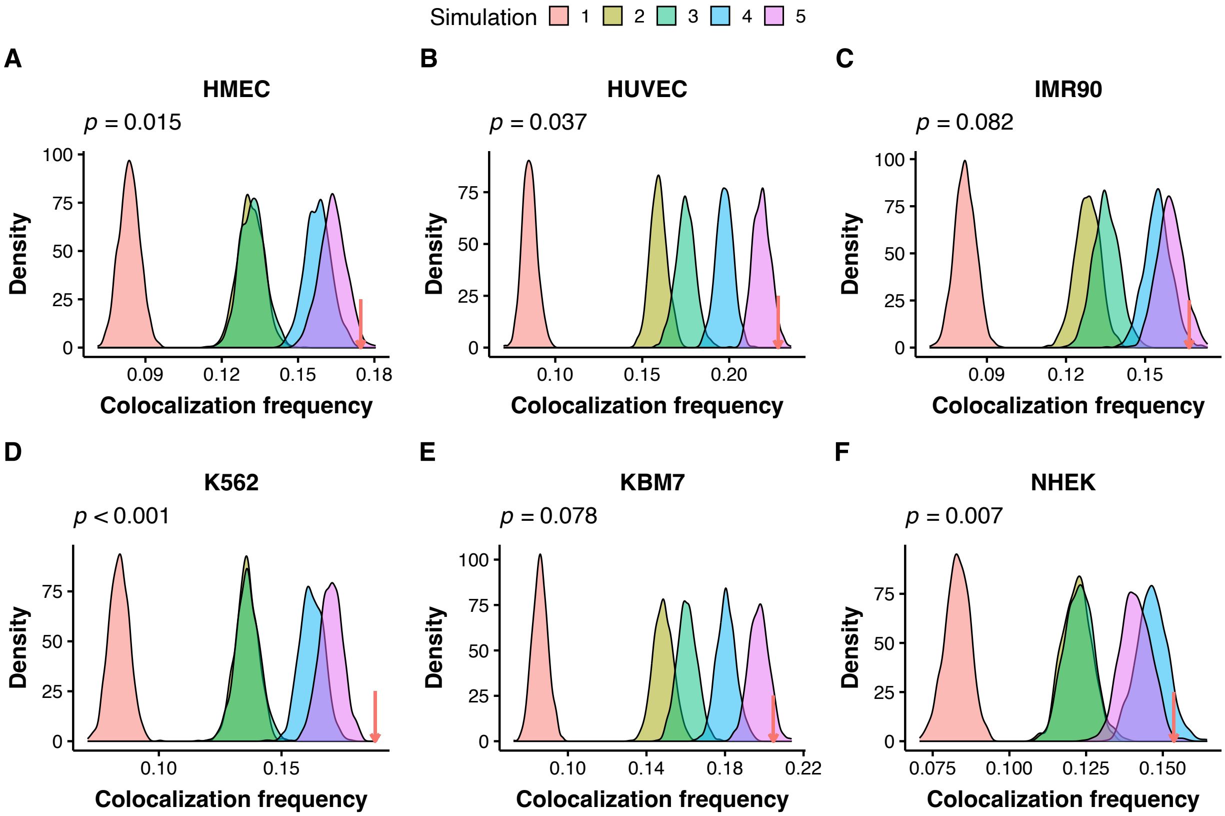


**Figure S3.** Density plots showing the distribution of colocalization frequency between chromatin pairs under different null models in HMEC (A), HUVEC (B), IMR90 (C), K562 (D), KBM7 (E), and NHEK (F). The red arrow denotes the colocalization frequency of true retrocopies and their parental genes in respective cell lines. *P*-value is indicated for simulation 5 in each plot.

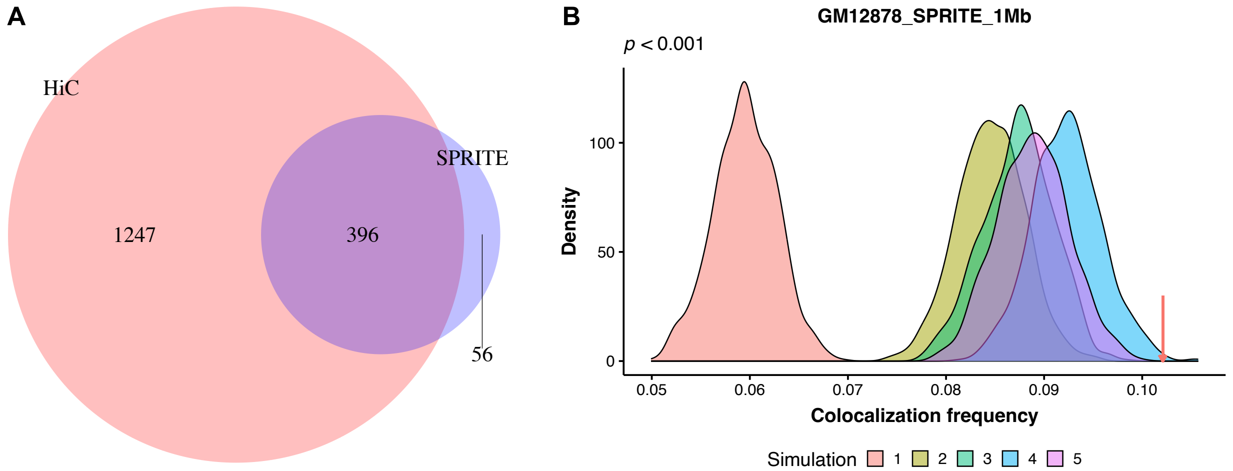


**Figure S4.** For the human GM12878 cell line, (A) displaying the overlap of colocalized retrocopy-parent pairs between Hi-C and SPRITE chromatin interaction data and (B) showing the distribution of colocalization frequency between chromatin pairs under different null models when using chromatin contact map derived from SPRITE. The red arrow denotes the colocalization frequency of true retrocopies and their parental genes. *P*-value is indicated for simulation 5. The resolution of both Hi-C and SPRITE is 1 Mb.

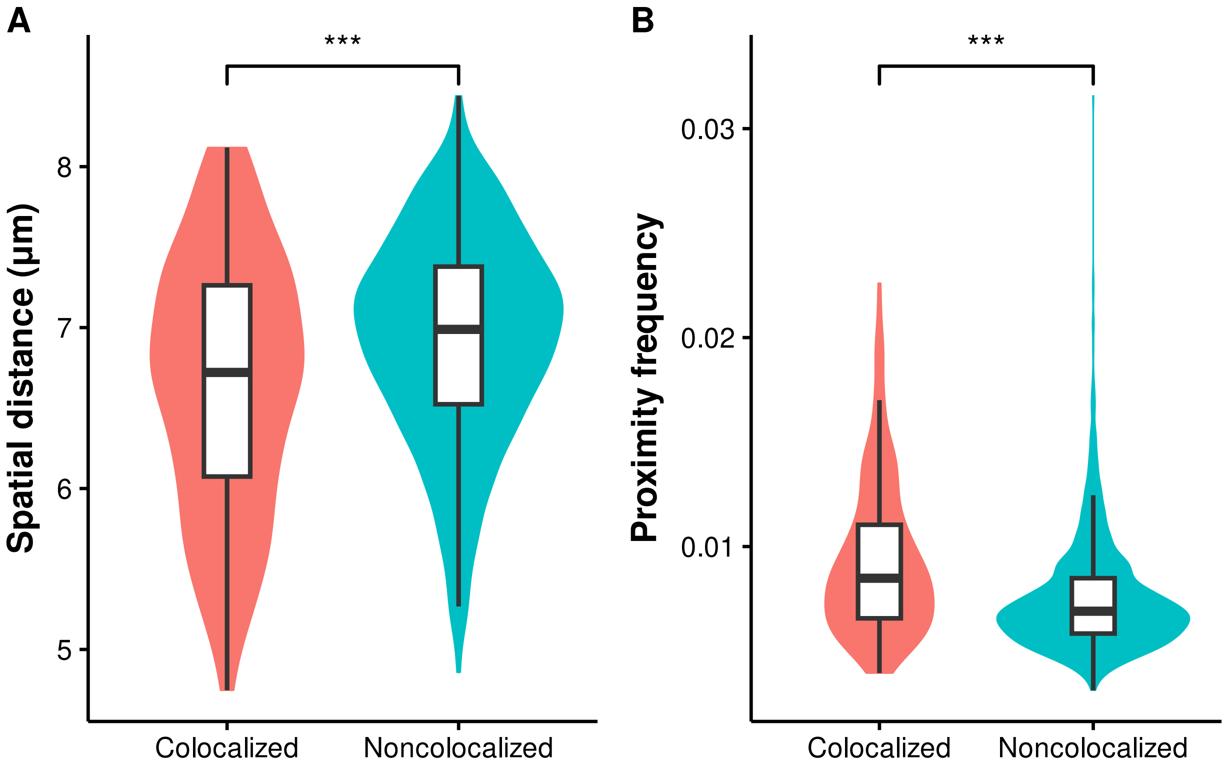


**Figure S5.** Comparison of median spatial distance (A) and proximity frequency (B) detected by MERFISH between colocalized and noncolocalized retrocopy-parent pairs in IMR90 cells. Proximity is defined when two loci is < 1 µm apart. ***$p < 0.001$; two-tailed Wilcoxon test.

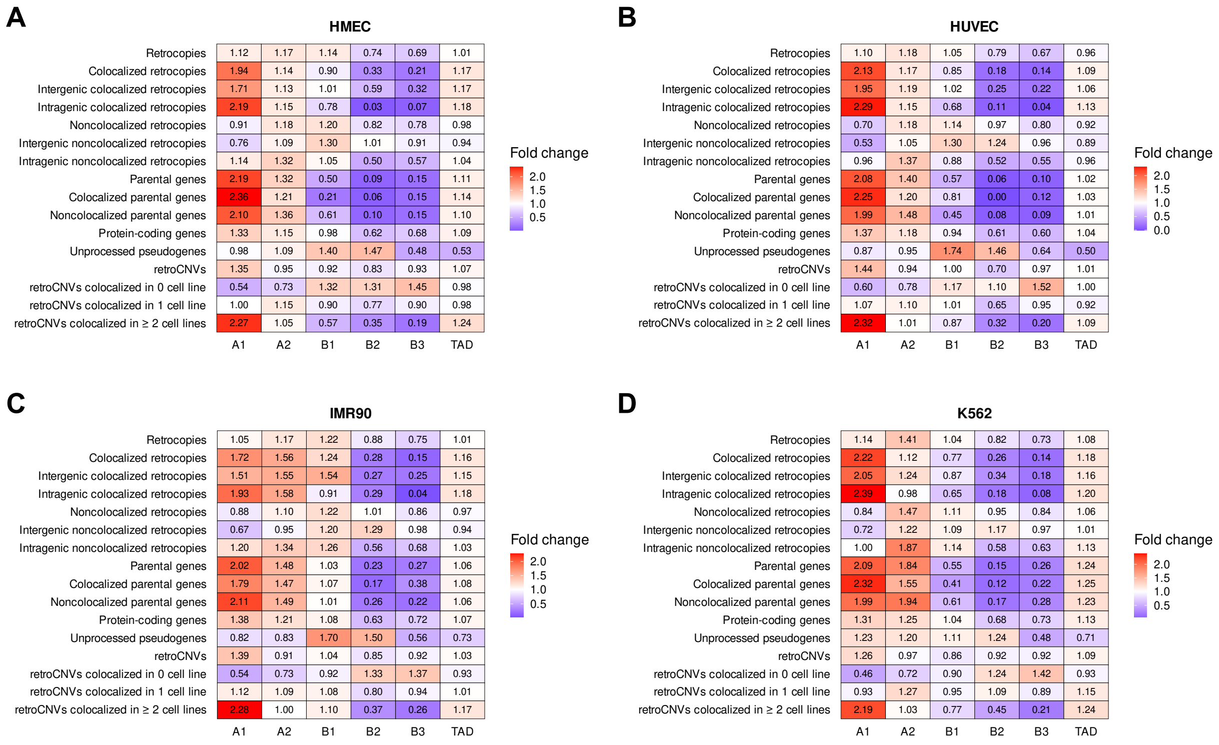


**Figure S6.** Enrichments of retrocopies, parental genes, unprocessed pseudogenes, and retroCNVs in five primary subcompartments and TAD in the HMEC (A), HUVEC (B), IMR90 (C), and K562 (D) cell lines.

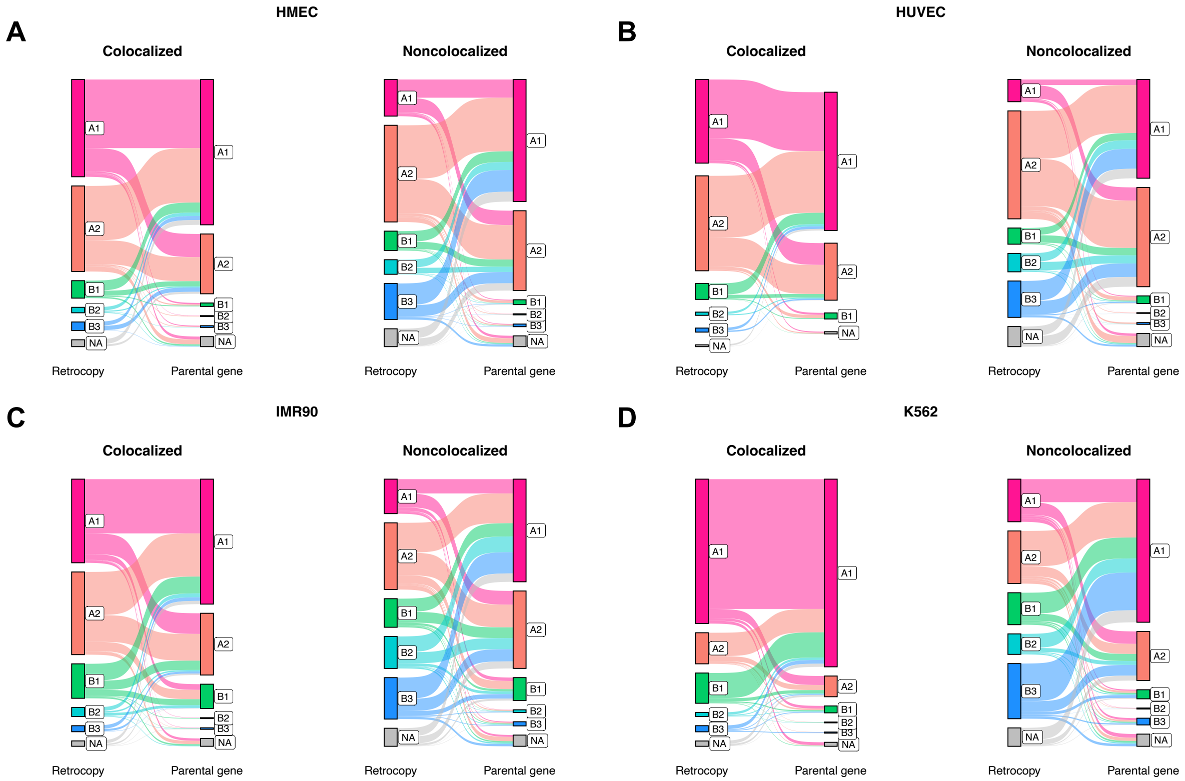


**Figure S7.** Sankey diagrams showing the connection of colocalized (left panel in each plot) and noncolocalized (right panel in each plot) retrocopy-parent pairs in the context of subcompartment annotations in HMEC (A), HUVEC (B), IMR90 (C), and K562 (D). The height of bar indicates the proportion of loci that fall in respective categories. Lines connect retrocopies and their corresponding parent genes. NA denotes a fragment is not annotated as any of the five subcompartments.

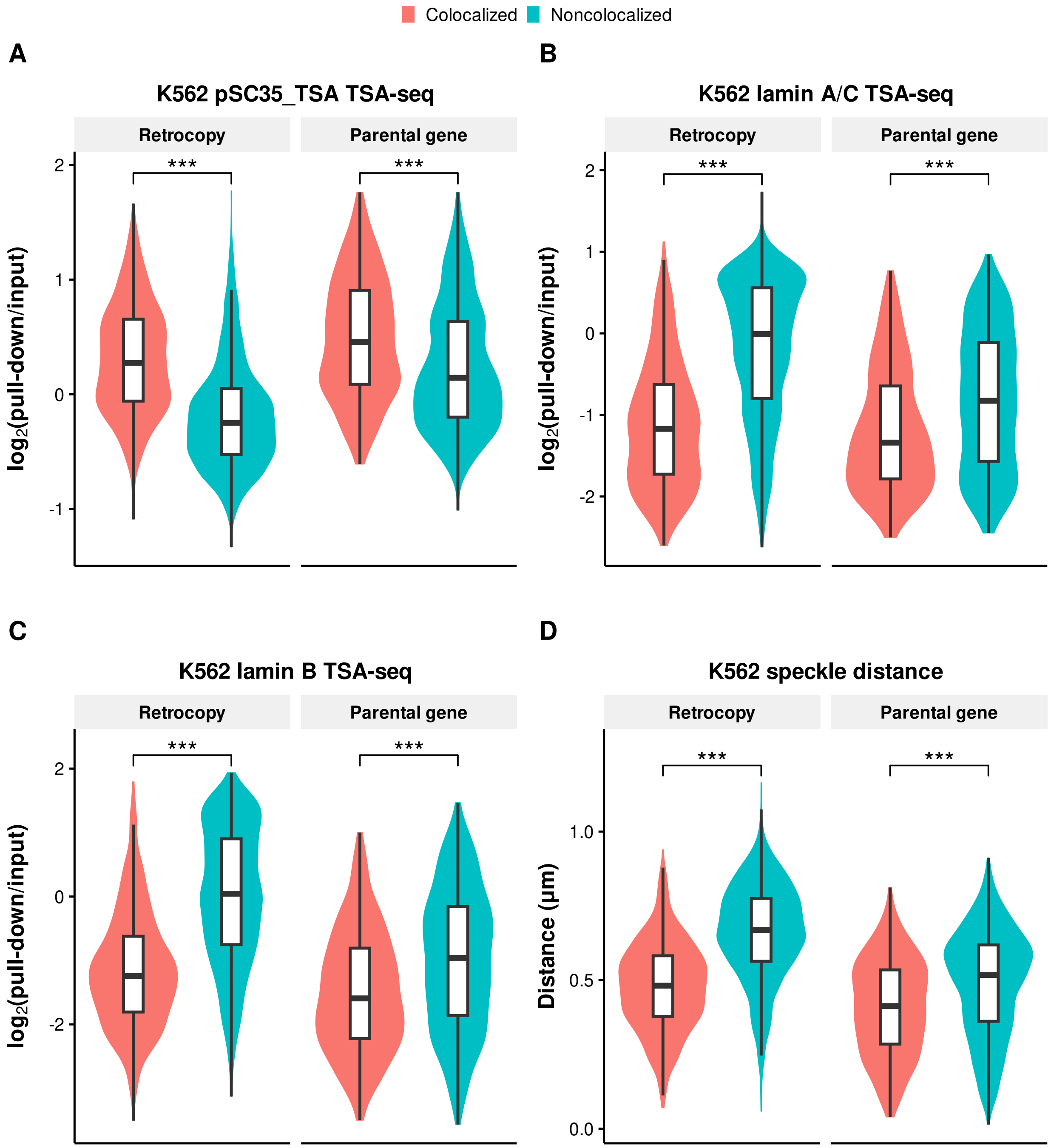


**Figure S8.** Comparison of pSC35 TSA-seq signal (A), lamin A/C TSA-seq signal (B), lamin B TSA-seq signal (C), and speckle distance (D) between colocalized and noncolocalized retrocopies/parental genes in K562. $^{***}p < 0.001$; two-tailed Wilcoxon test.

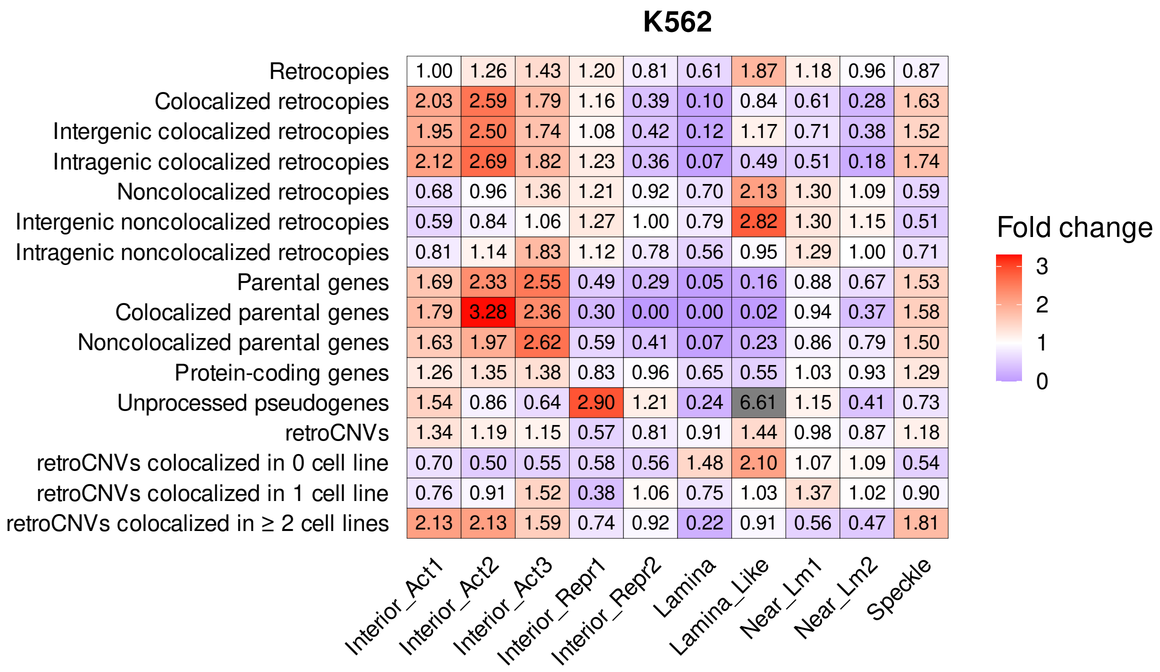


**Figure S9.** Enrichments of retrocopies, parental genes, unprocessed pseudogenes, and retroCNVs in 10 different spatial compartmentalization states relative to nuclear speckles, lamina, and nucleoli defined by SPIN. The 10 SPIN states are: Interior Active 1, 2, 3 (Interior_Act1, Interior_Act2, Interior_Act3), Interior Repressive 1, 2 (Interior_Repr1, Interior_Repr2), Lamina, Lamina_Like, Near Lamina 1, 2 (Near_Lm1, Near_Lm2), and Speckle.

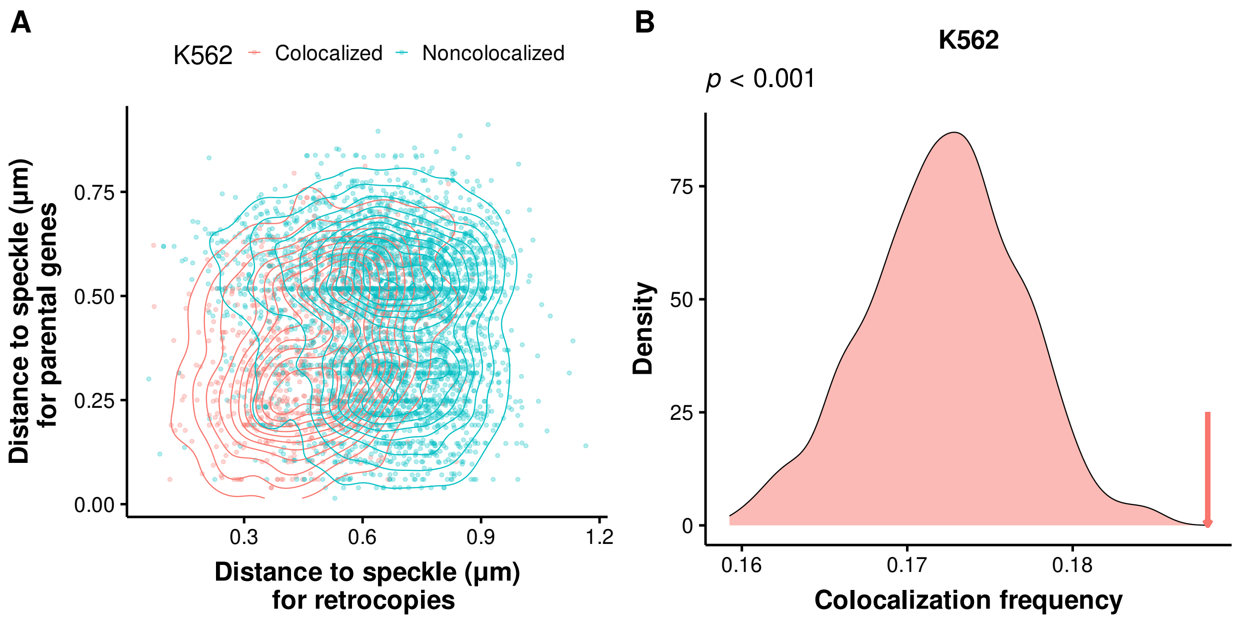


**Figure S10.** (A) Scatter plot showing the distances to speckles for colocalized (red dots and contours) and noncolocalized (blue dots and contours) retrocopy-parent pairs in K562 cells. (B) The distribution of colocalization frequency between retrocopies and randomly sampled protein coding genes with matching speckle distances (± 0.01 μm away) as true parental genes. The red arrow denotes the colocalization frequency of true retrocopies and their parental genes in the K562 cell line.

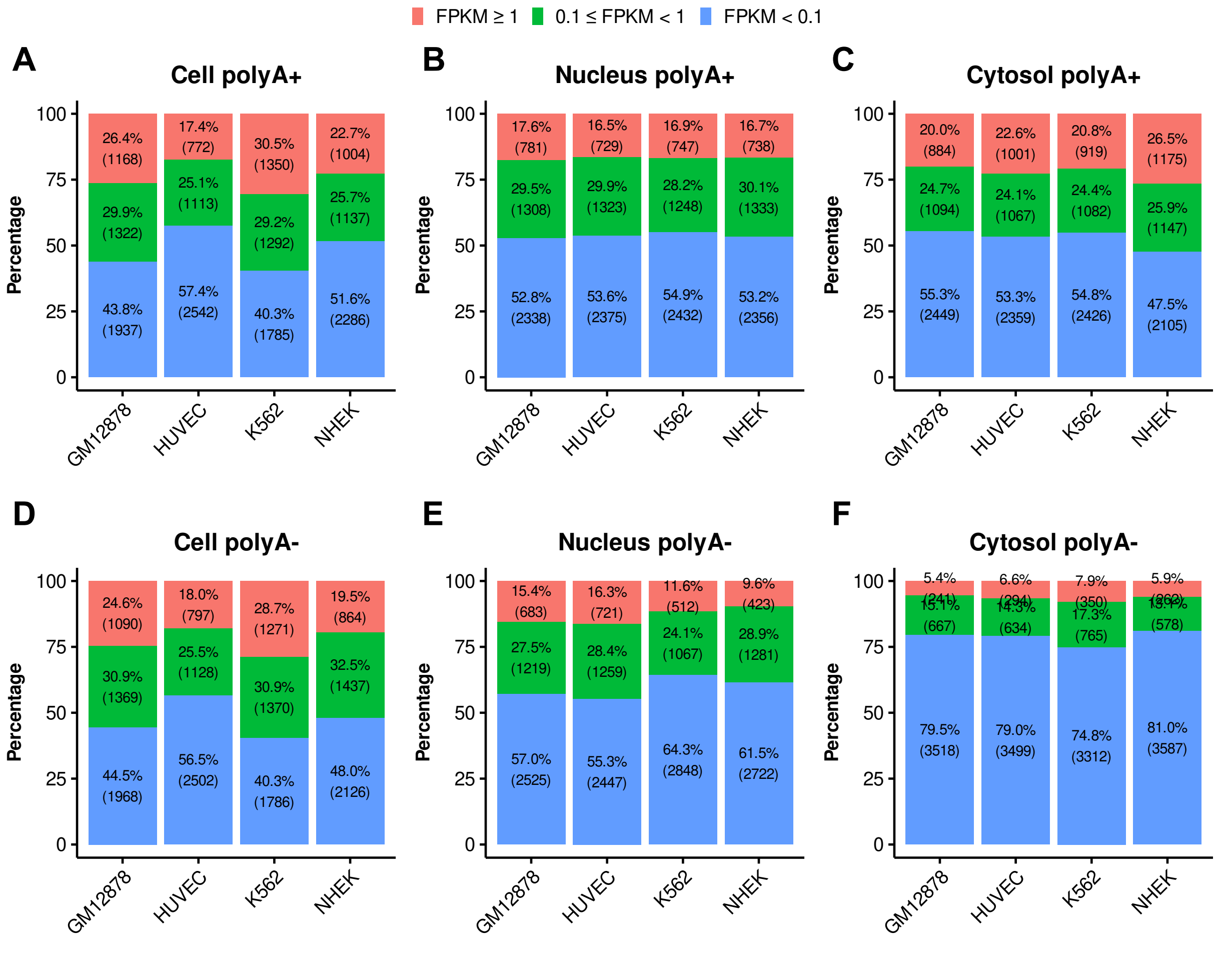


**Figure S11.** Proportion of retrocopies that expressed with FPKM ≥ 1 (red), 0.1 ≤ FPKM < 1 (green), and FPKM ≤ 0.1 (blue) in GM12878, HUVEC, K562, and NHEK. RNA expression levels derived from different cell compartments and fractions are plotted separately. (A) Polyadenylated (polyA+) RNAs from the whole cell; (B) PolyA+ RNAs from nucleus; (C) PolyA+ from cytosol; (D) Non-polyadenylated (polyA-) RNAs from the whole cell; (E) PolyA- RNAs from nucleus; (F) PolyA- from cytosol.

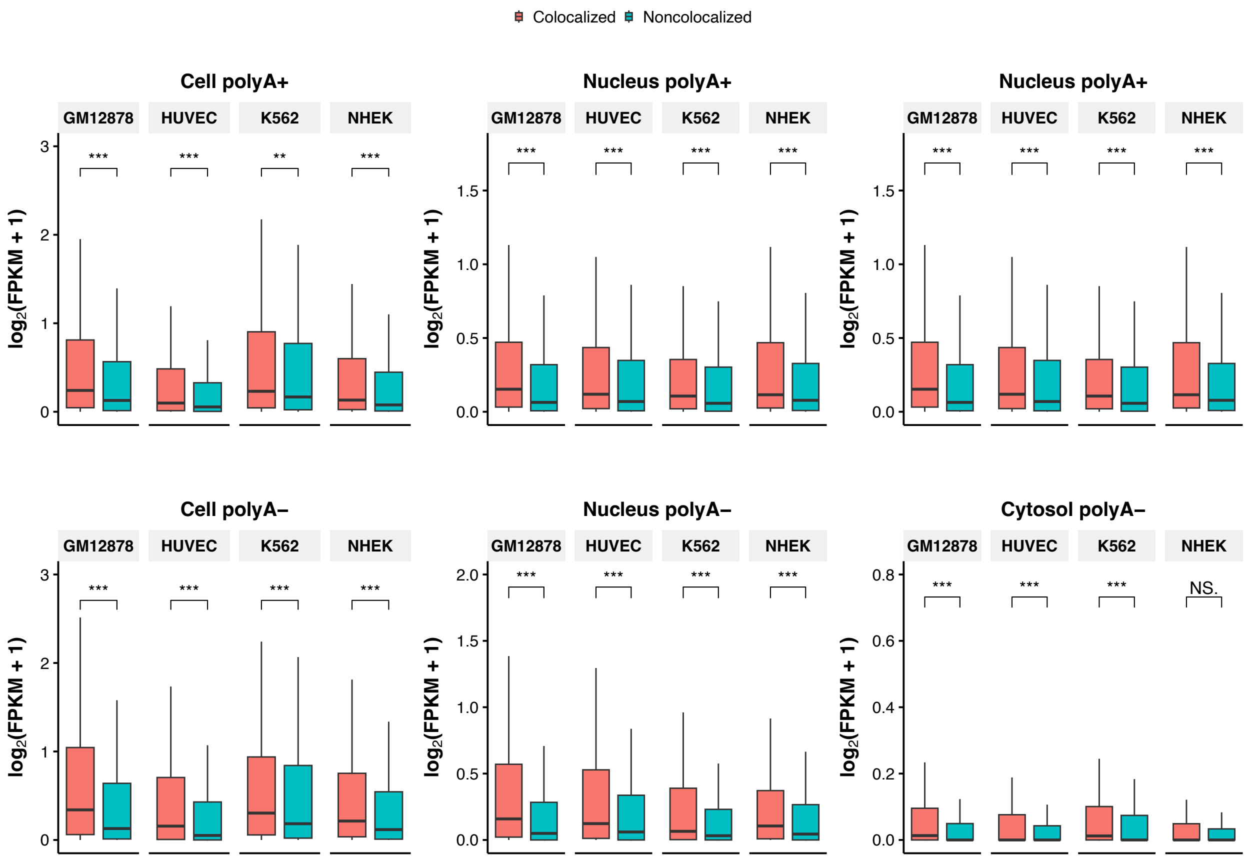


**Figure S12.** Comparison of expression level between colocalized and nonlocalized retrocopies in GM12878, HUVEC, K562, and NHEK, for RNAs derived from different cell compartments and fractions. Cell polyA+: polyadenylated (polyA+) RNAs from the whole cell; Nucleus polyA+: polyA+ RNAs from nucleus; Cytosol polyA+: polyA+ RNAs from cytosol; Cell polyA-: non-polyadenylated (polyA-) RNAs from the whole cell; Nucleus polyA-: polyA- RNAs from nucleus; Cytosol polyA-: polyA- RNAs from cytosol. $^{***}p < 0.001$, $^{**}p < 0.01$, NS. not significant; two-tailed Wilcoxon test.

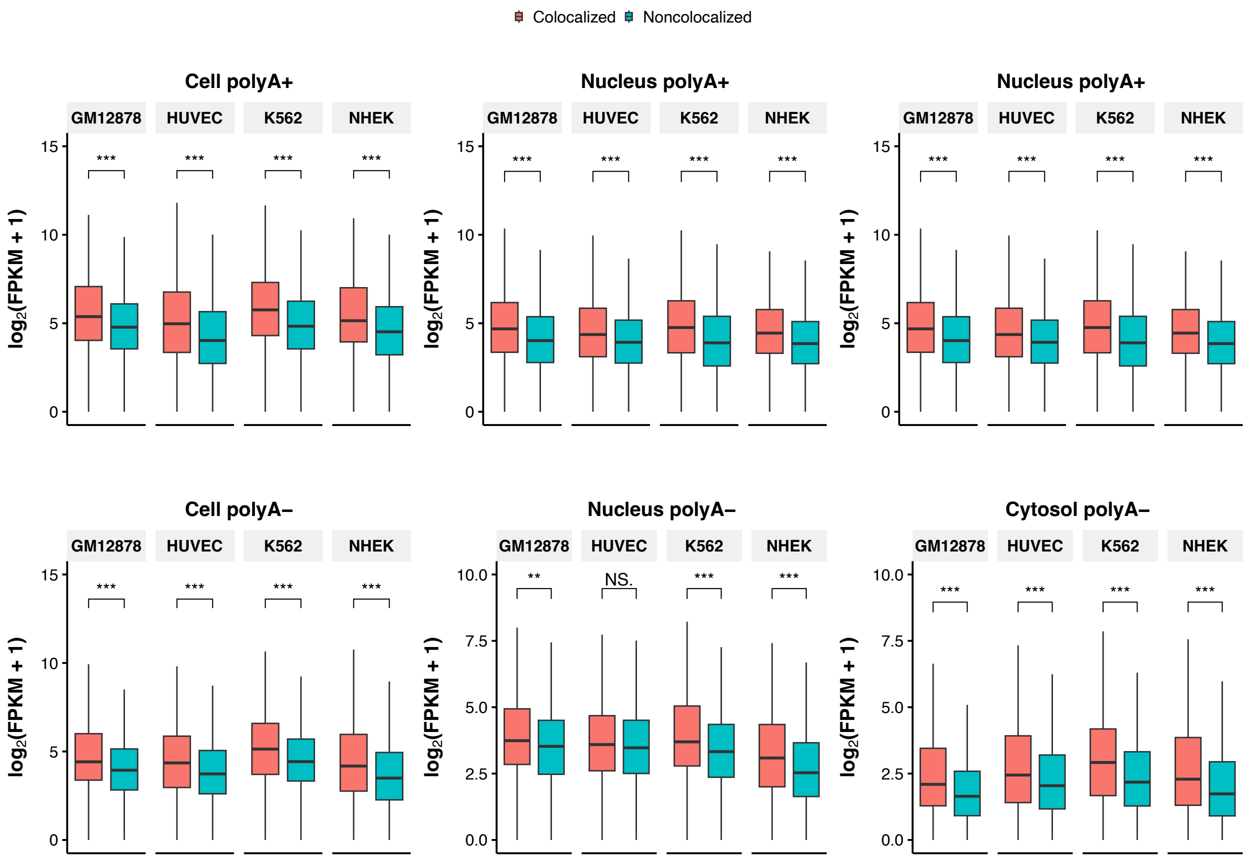


**Figure S13.** Comparison of expression level between colocalized and nonlocalized parental genes in GM12878, HUVEC, K562, and NHEK, for RNAs derived from different cell compartments and fractions. Cell polyA+: polyadenylated (polyA+) RNAs from the whole cell; Nucleus polyA+: polyA+ RNAs from nucleus; Cytosol polyA+: polyA+ RNAs from cytosol; Cell polyA-: non-polyadenylated (polyA-) RNAs from the whole cell; Nucleus polyA-: polyA- RNAs from nucleus; Cytosol polyA-: polyA- RNAs from cytosol. $^{***}p < 0.001$, $^{**}p < 0.01$, NS. not significant; two-tailed Wilcoxon test.

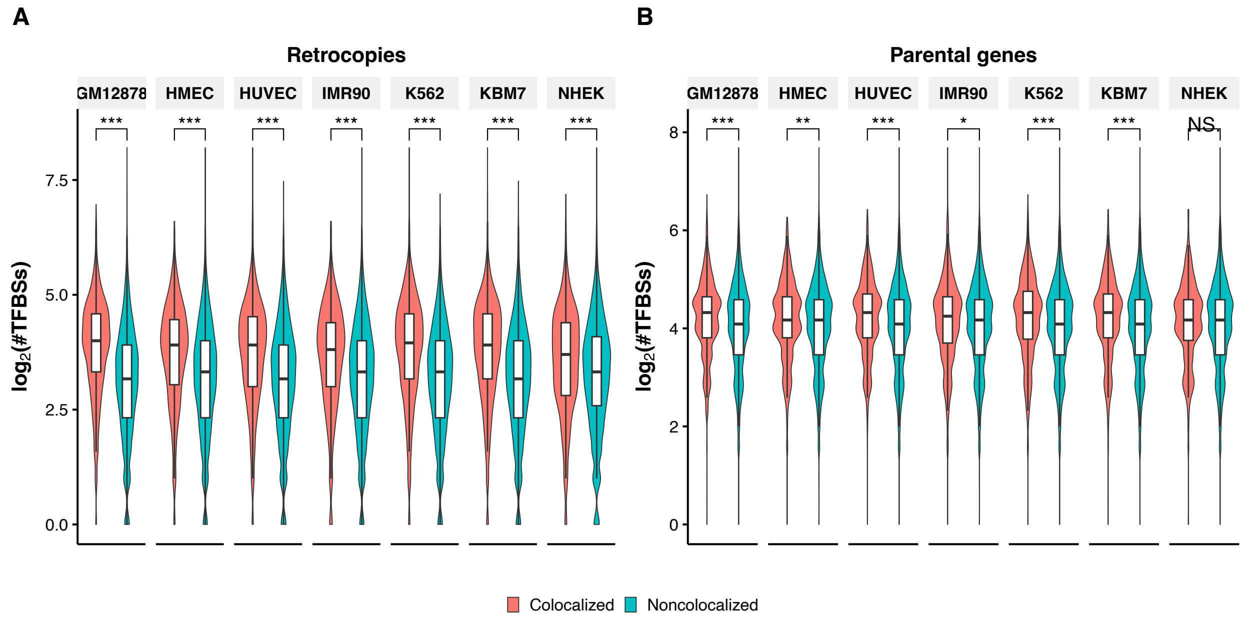


**Figure S14.** Comparison of the number of harbored TFBSs between colocalized and nonlocalized retrocopies (A) and between colocalized and noncolocalized parental genes (B). The number of TFBSs were based on the flanking regions (putative TSS ± 3 kb) of retrocopies and parental genes. $^{***}p < 0.001$, $^{**}p < 0.01$, $^{*}p < 0.05$, NS. not significant; two-tailed Wilcoxon test.

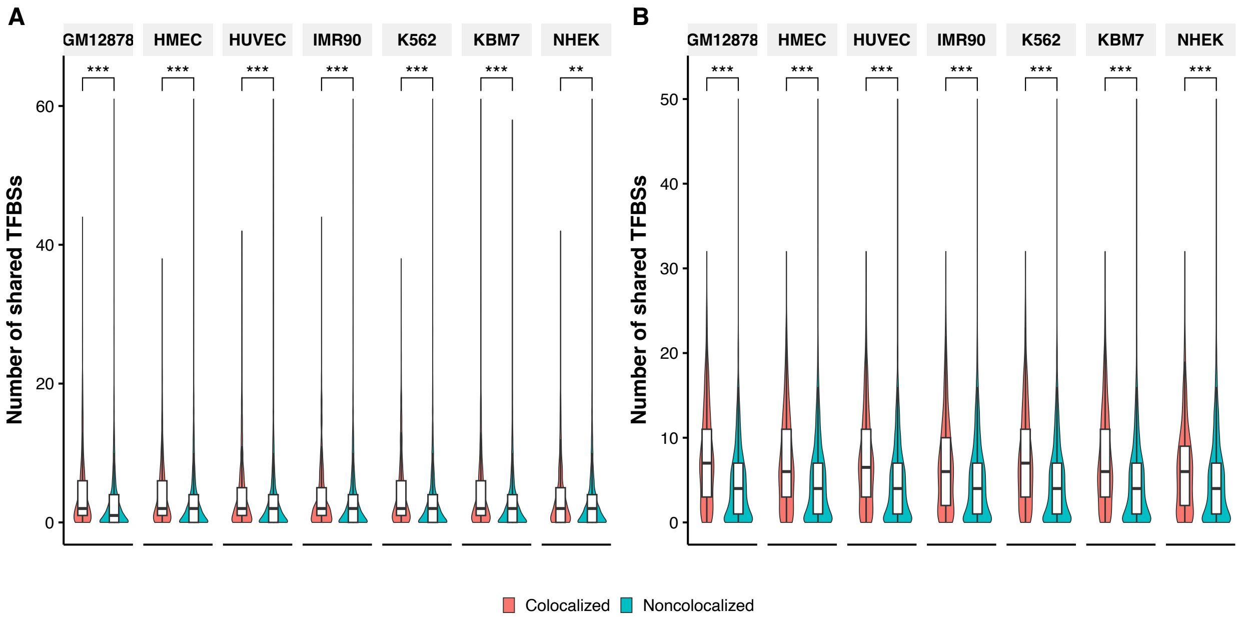


**Figure S15.** Comparison of the number of shared TFBSs between colocalized and noncolocalized retrocopy-parent pairs for regions of upstream and downstream 1 kb (A) and upstream 3 kb (B) of putative TSSs. $^{***}p < 0.001$, $^{**}p < 0.01$; two-tailed Wilcoxon test.

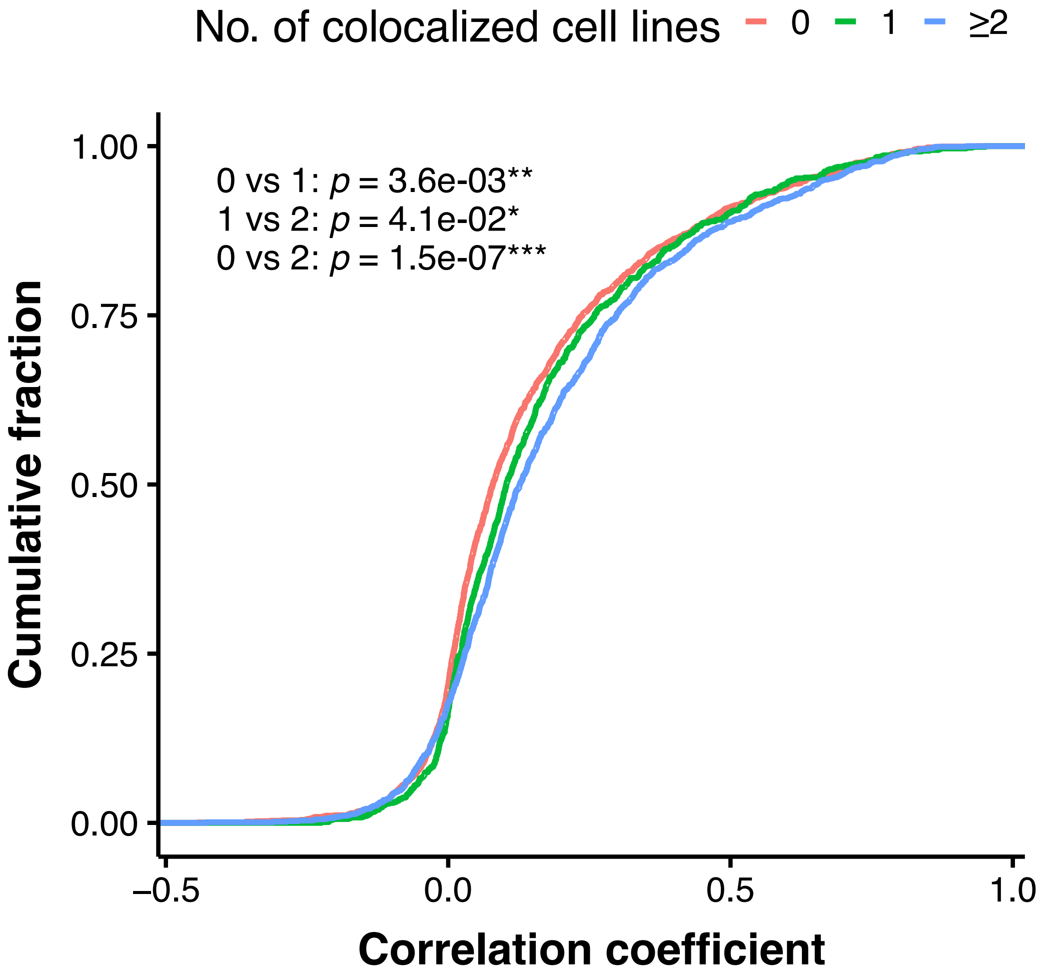


**Figure S16.** The cumulative fraction of correlation coefficient across GTEx samples for individual retrocopies that are colocalized in 0, 1, or ⩾ 2 cell lines.

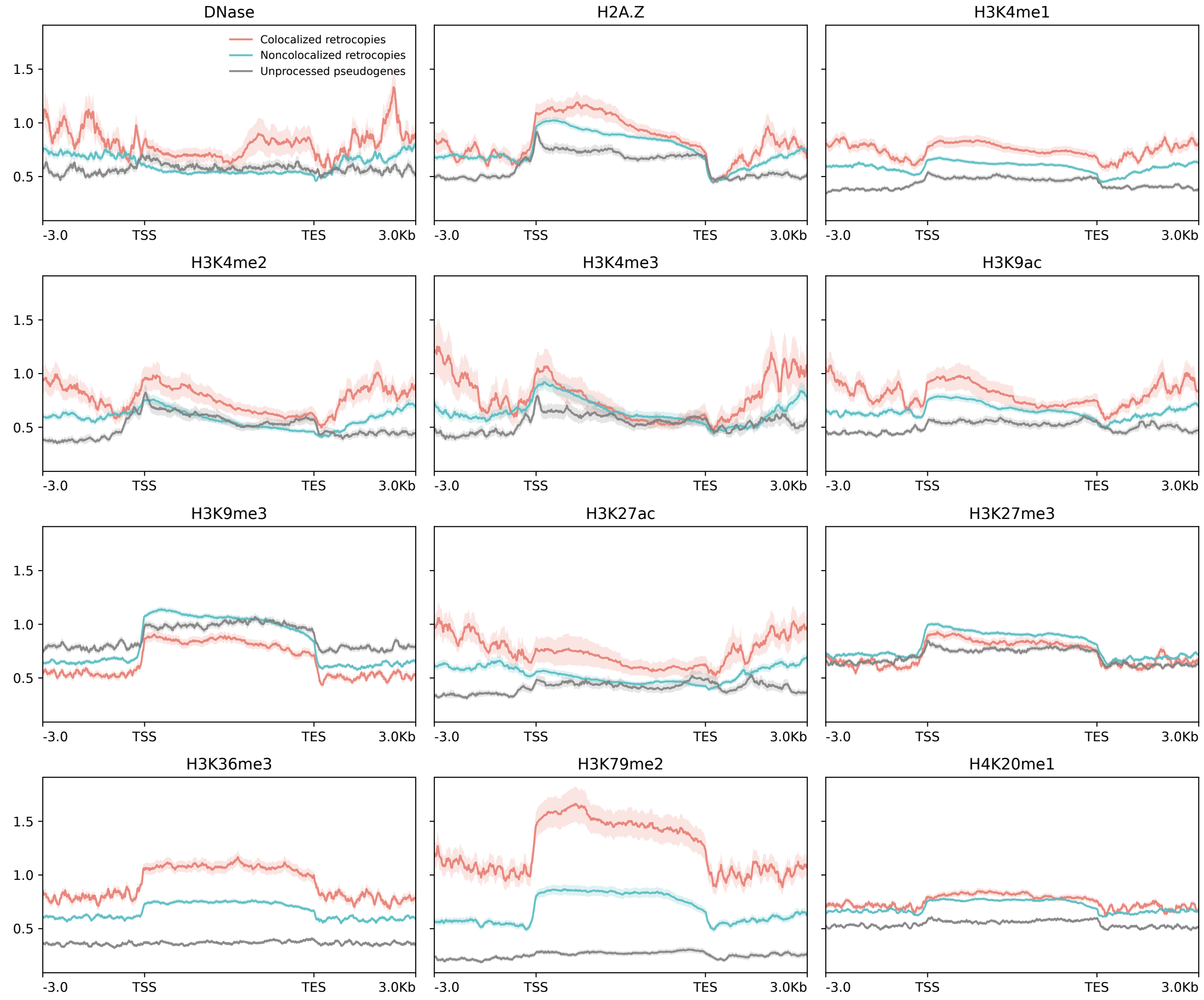


**Figure S17.** Epigenetic signals for DNase I hypersensitivity, histone variant H2A.Z, and 10 histone modifications along colocalized retrocopies (red lines), noncolocalized retrocopies (blue lines), and unprocessed pseudogenes (gray lines) and their flanking regions (± 3 kb) in the HMEC cell line. Solid lines represent the mean fold change signal, shadowed areas denote the standard error. TSS: putative transcription start site; TES: putative transcription end site.

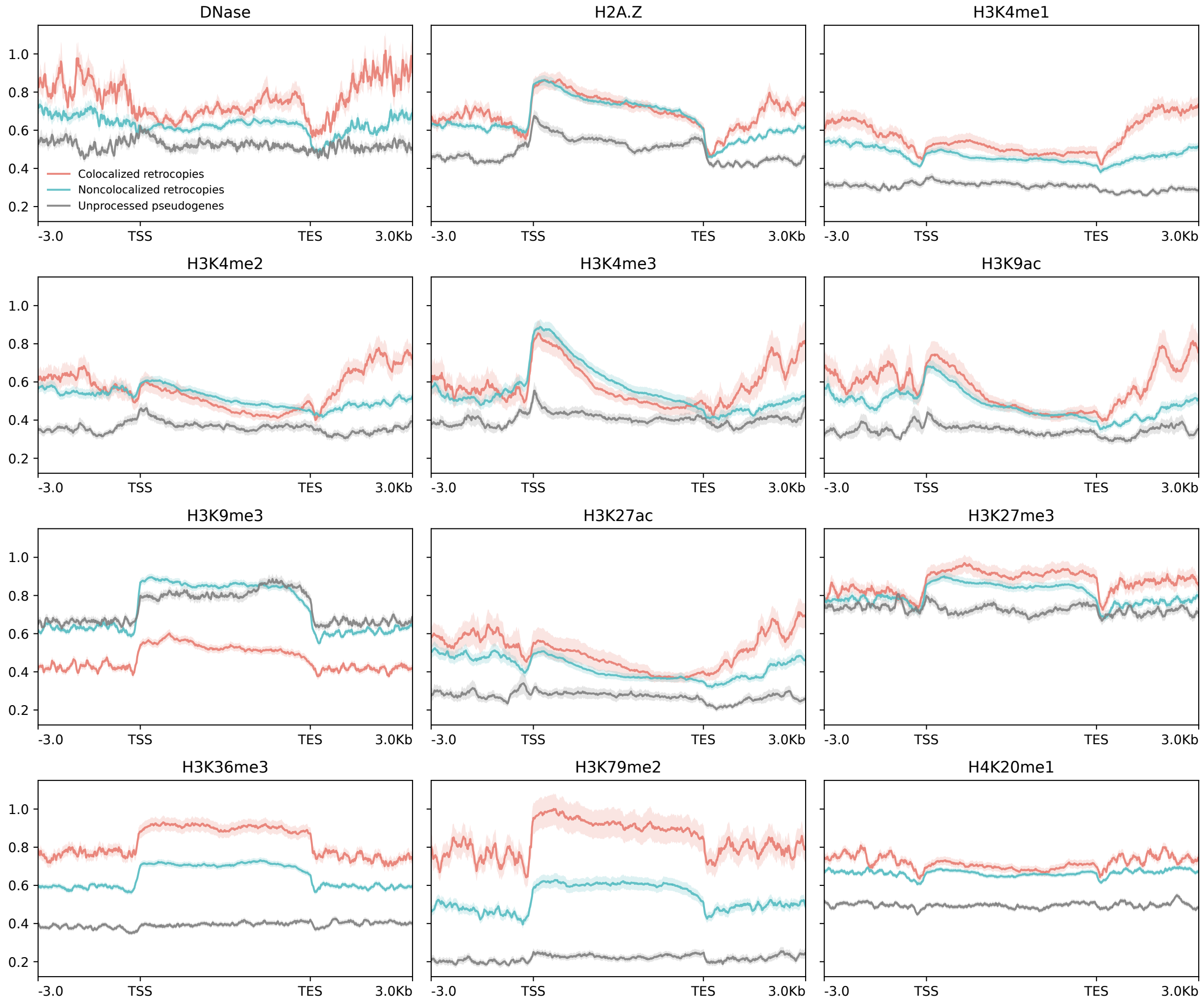


**Figure S18.** Epigenetic signals for DNase I hypersensitivity, histone variant H2A.Z, and 10 histone modifications along colocalized retrocopies (red lines), noncolocalized retrocopies (blue lines), and unprocessed pseudogenes (gray lines) and their flanking regions (± 3 kb) in the HUVEC cell line. Solid lines represent the mean fold change signal, shadowed areas denote the standard error. TSS: putative transcription start site; TES: putative transcription end site.

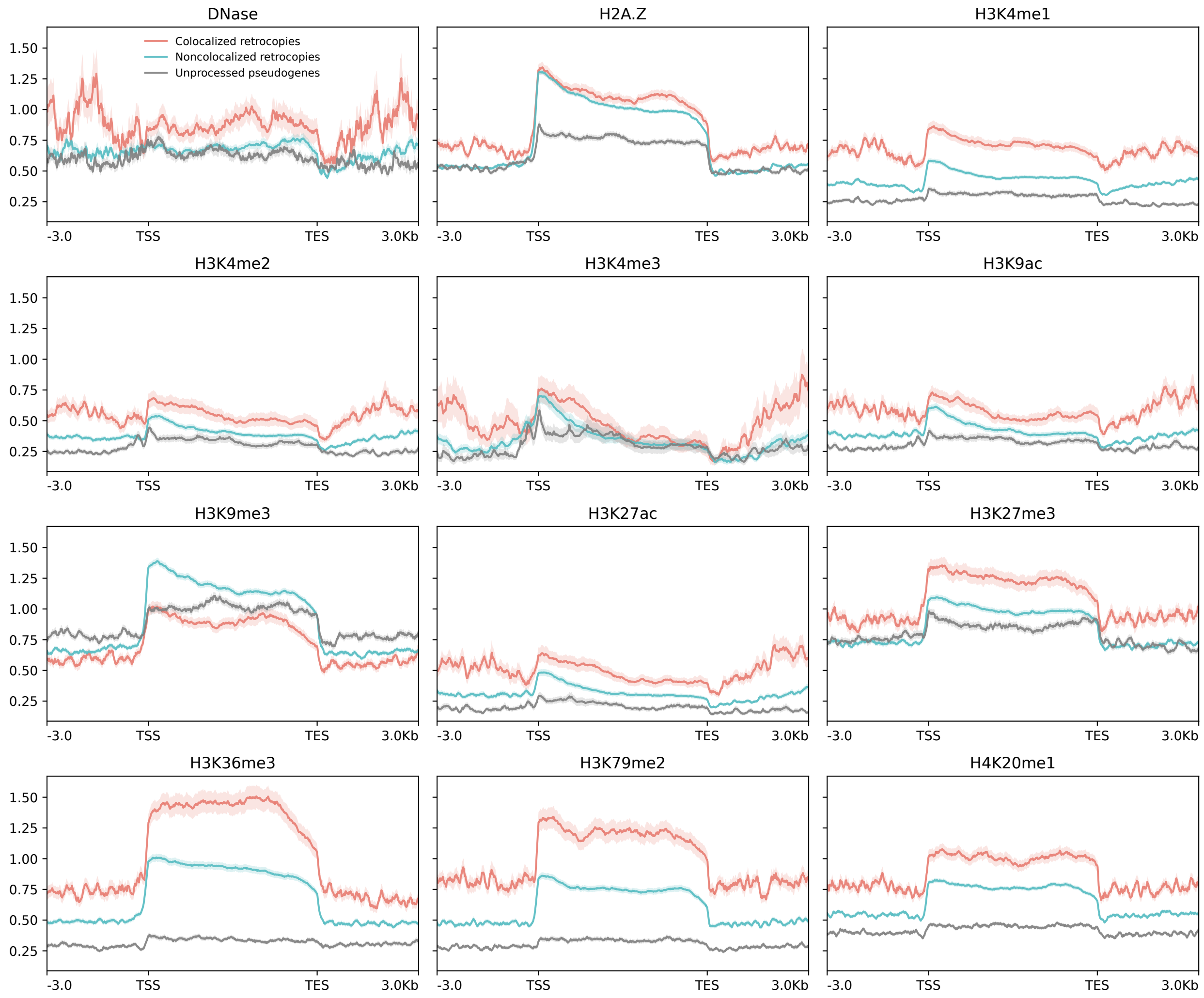


**Figure S19.** Epigenetic signals for DNase I hypersensitivity, histone variant H2A.Z, and 10 histone modifications along colocalized retrocopies (red lines), noncolocalized retrocopies (blue lines), and unprocessed pseudogenes (gray lines) and their flanking regions (± 3 kb) in the IMR90 cell line. Solid lines represent the mean fold change signal, shadowed areas denote the standard error. TSS: putative transcription start site; TES: putative transcription end site.

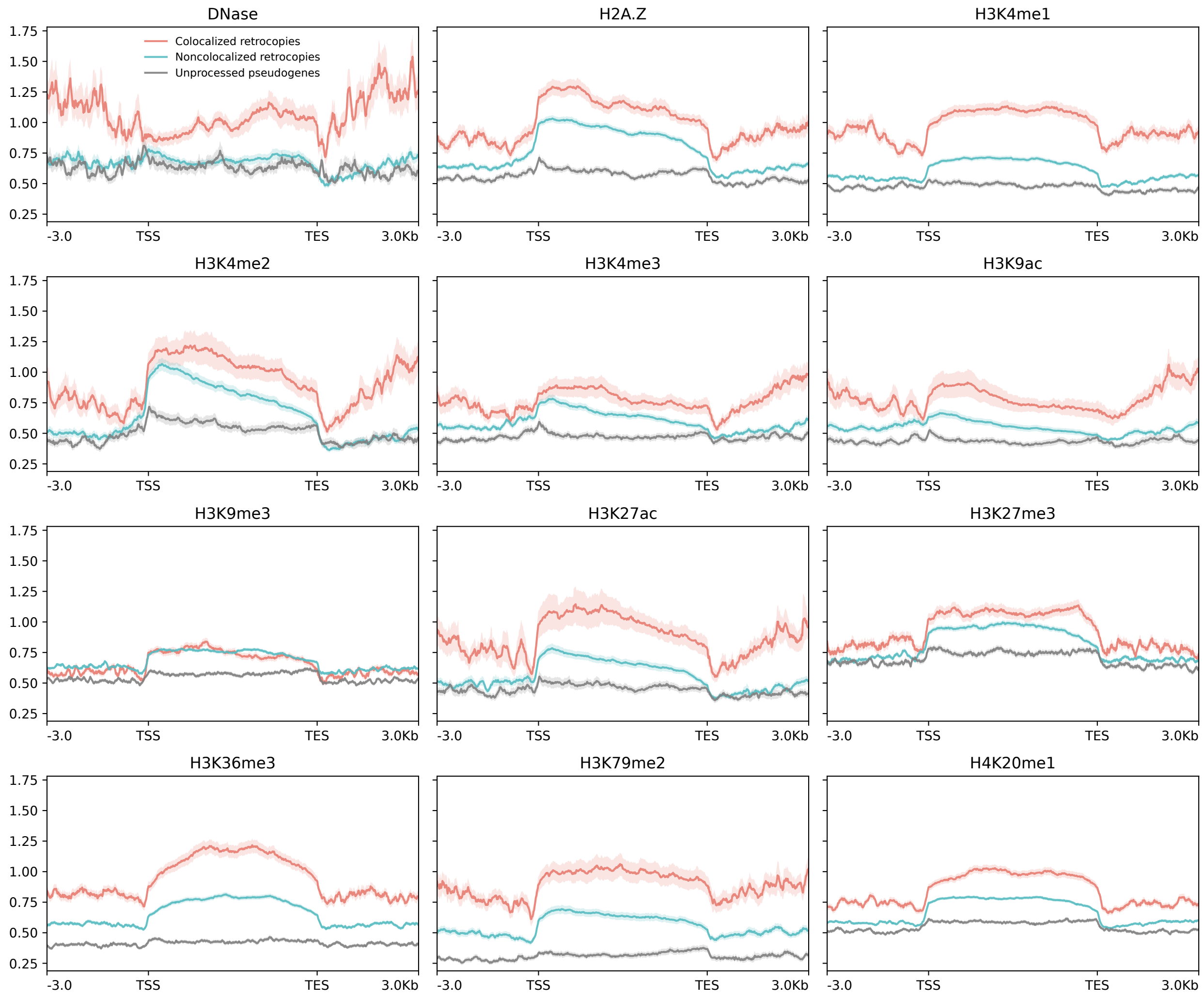


**Figure S20.** Epigenetic signals for DNase I hypersensitivity, histone variant H2A.Z, and 10 histone modifications along colocalized retrocopies (red lines), noncolocalized retrocopies (blue lines), and unprocessed pseudogenes (gray lines) and their flanking regions (± 3 kb) in the K562 cell line. Solid lines represent the mean fold change signal, shadowed areas denote the standard error. TSS: putative transcription start site; TES: putative transcription end site.

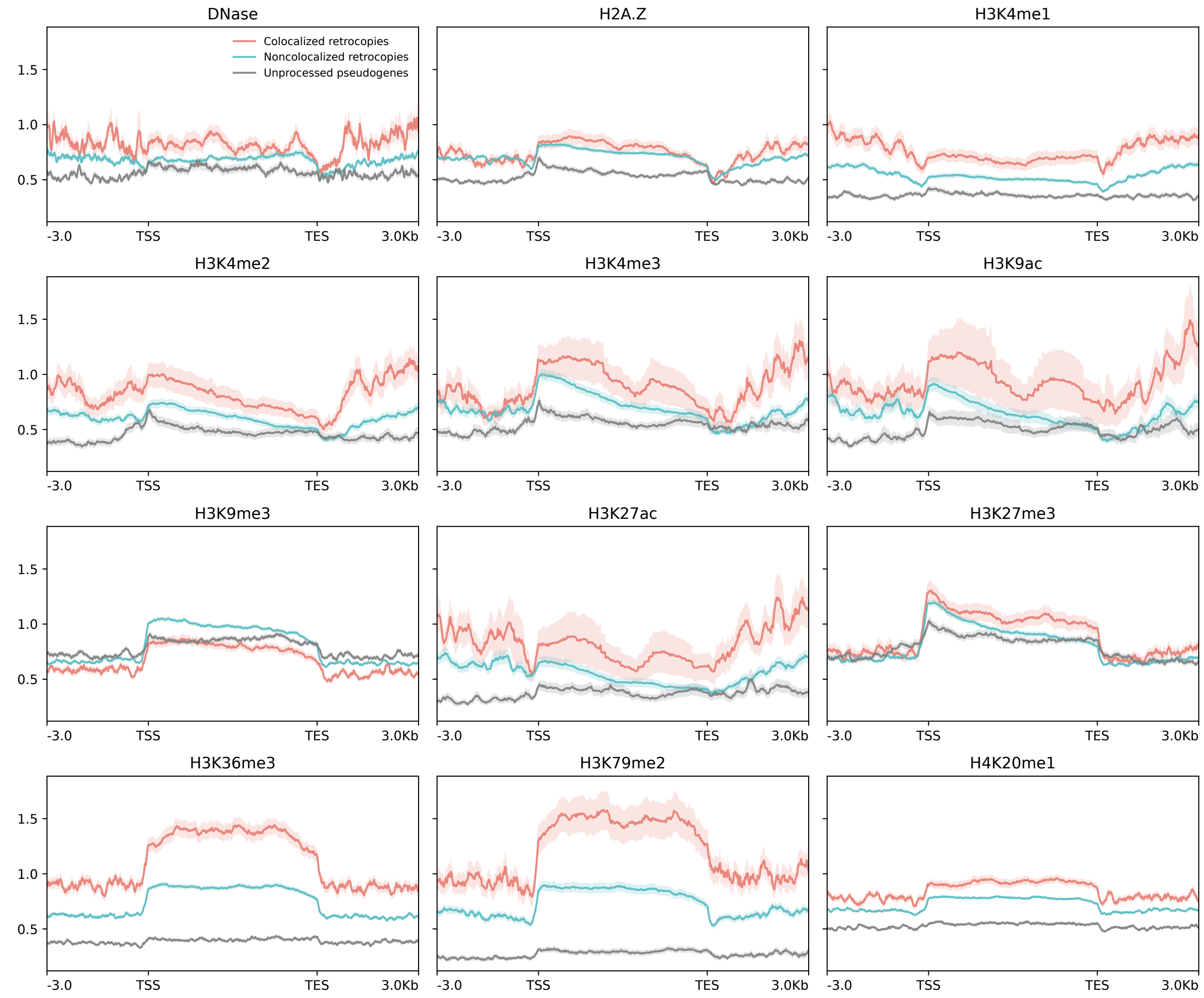


**Figure S21.** Epigenetic signals for DNase I hypersensitivity, histone variant H2A.Z, and 10 histone modifications along colocalized retrocopies (red lines), noncolocalized retrocopies (blue lines), and unprocessed pseudogenes (gray lines) and their flanking regions (± 3 kb) in the NHEK cell line. Solid lines represent the mean fold change signal, shadowed areas denote the standard error. TSS: putative transcription start site; TES: putative transcription end site.

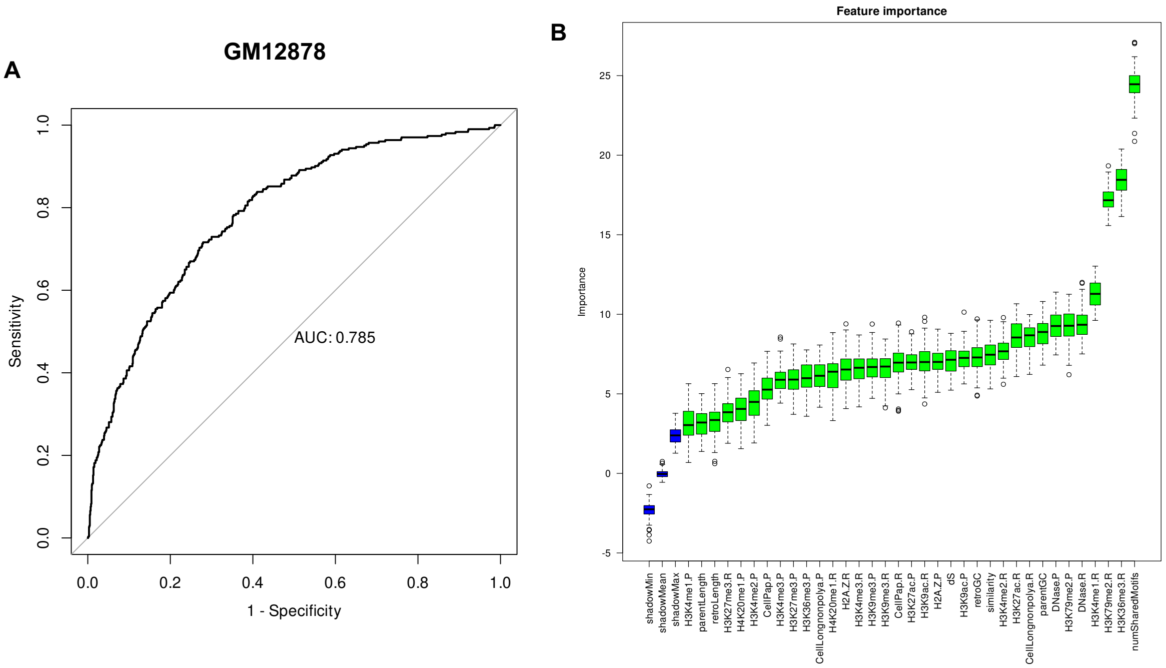


**Figure S22.** Performance and feature importance for the logistic regression model in GM12878. (A) Area under the receiver operating characteristic curve (AUC) of the model. (B) The relative importance of 35 genetic and epigenetic features of retrocopies and parental genes used in the model. numSharedMotifs: number of shared motifs; similarity: the percentage of identical sequence between a retrocopy and its parental gene; dS: synonymous substitution rate between a retrocopy and its parental gene; CellPap: FPKM of polyadenylated RNAs in the whole cell; CellLongnonpolya: FPKM of non-polyadenylated RNAs in the whole cell. The suffix ".R" means it's a feature of retrocopies and the suffix ".P" denotes it's a feature of parental genes. See supplementary table S2 for a detailed description of all features.

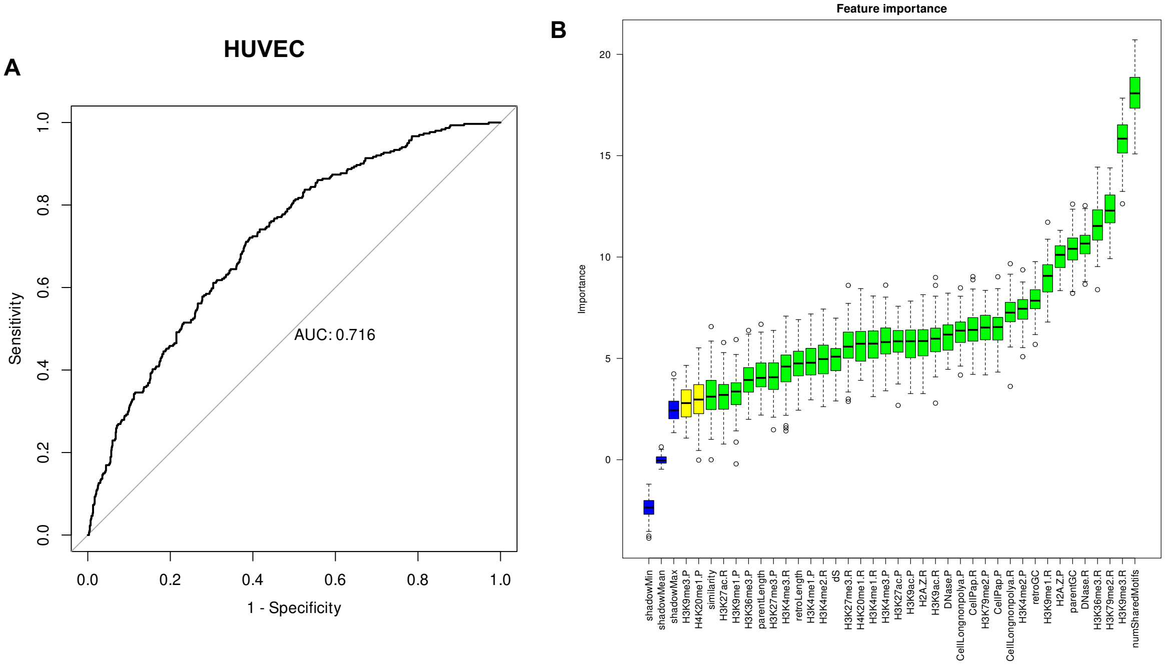


**Figure S23.** Performance and feature importance for the logistic regression model in HUVEC. (A) Area under the receiver operating characteristic curve (AUC) of the model. (B) The relative importance of 35 genetic and epigenetic features of retrocopies and parental genes used in the model. numSharedMotifs: number of shared motifs; similarity: the percentage of identical sequence between a retrocopy and its parental gene; dS: synonymous substitution rate between a retrocopy and its parental gene; CellPap: FPKM of polyadenylated RNAs in the whole cell; CellLongnonpolya: FPKM of non-polyadenylated RNAs in the whole cell. The suffix ".R" means it's a feature of retrocopies and the suffix ".P" denotes it's a feature of parental genes. See supplementary table S2 for a detailed description of all features.

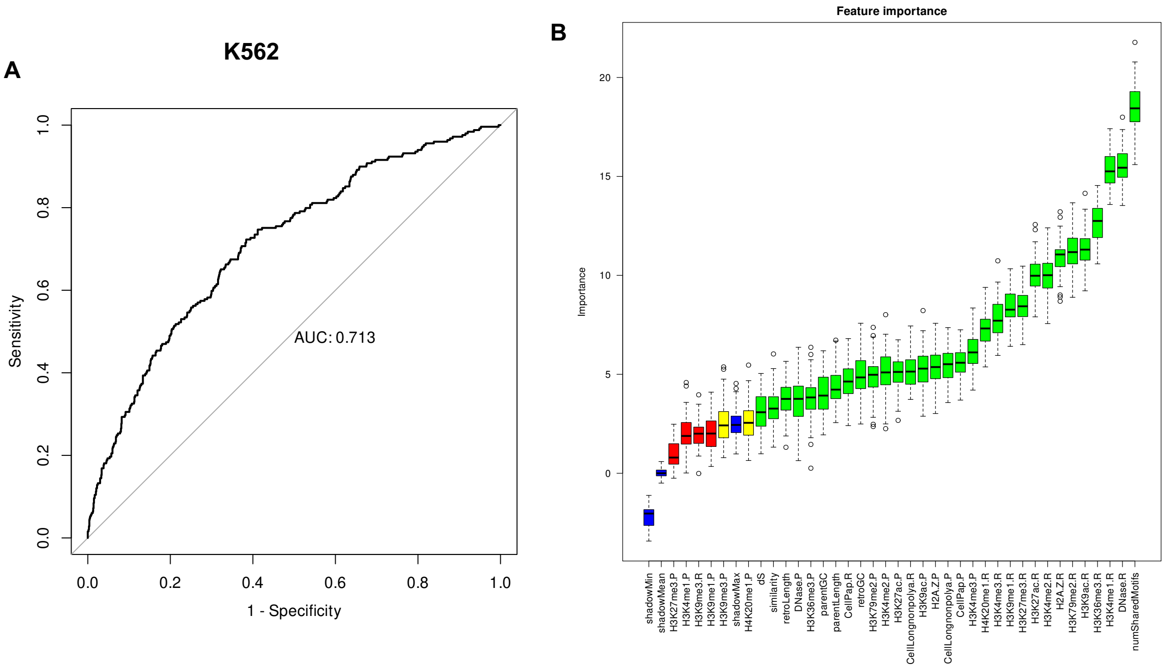


**Figure S24.** Performance and feature importance for the logistic regression model in K562. (A) Area under the receiver operating characteristic curve (AUC) of the model. (B) The relative importance of 35 genetic and epigenetic features of retrocopies and parental genes used in the model. numSharedMotifs: number of shared motifs; similarity: the percentage of identical sequence between a retrocopy and its parental gene; dS: synonymous substitution rate between a retrocopy and its parental gene; CellPap: FPKM of polyadenylated RNAs in the whole cell; CellLongnonpolya: FPKM of non-polyadenylated RNAs in the whole cell. The suffix ".R" means it's a feature of retrocopies and the suffix ".P" denotes it's a feature of parental genes. See supplementary table S2 for a detailed description of all features.

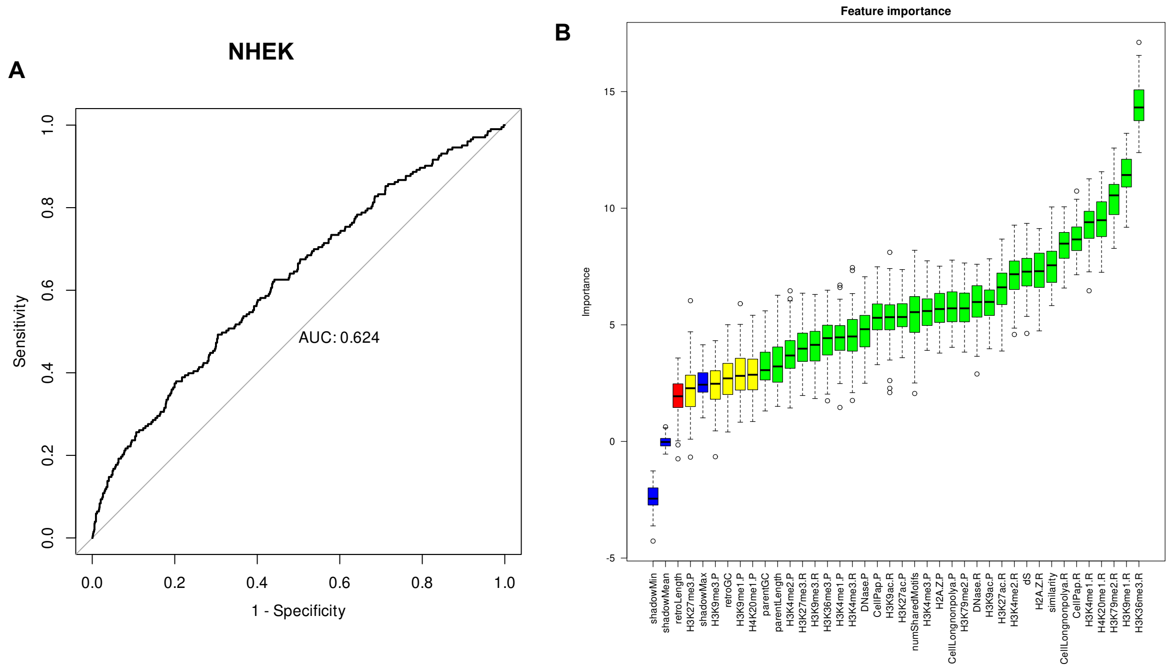


**Figure S25.** Performance and feature importance for the logistic regression model in NHEK. (A) Area under the receiver operating characteristic curve (AUC) of the model. (B) The relative importance of 35 genetic and epigenetic features of retrocopies and parental genes used in the model. numSharedMotifs: number of shared motifs; similarity: the percentage of identical sequence between a retrocopy and its parental gene; dS: synonymous substitution rate between a retrocopy and its parental gene; CellPap: FPKM of polyadenylated RNAs in the whole cell; CellLongnonpolya: FPKM of non-polyadenylated RNAs in the whole cell. The suffix ".R" means it's a feature of retrocopies and the suffix ".P" denotes it's a feature of parental genes. See supplementary table S2 for a detailed description of all features.

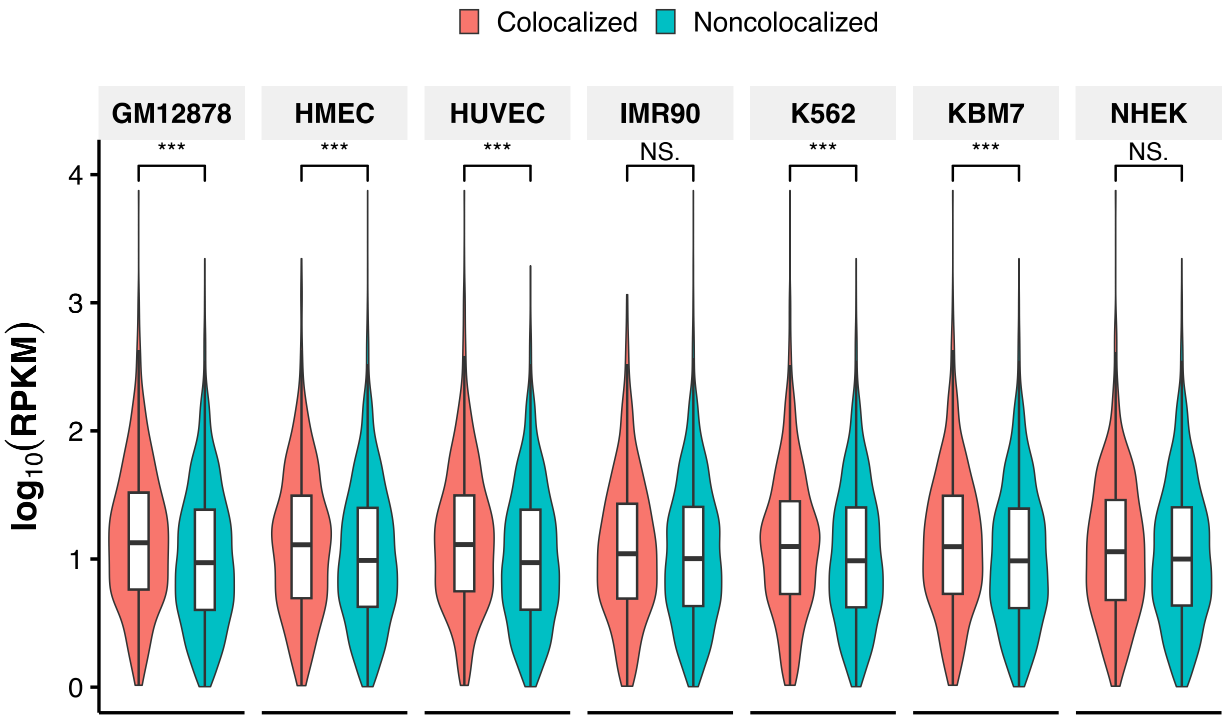


**Figure S26.** Comparison of ribo-seq RPKM values between colocalized and noncolocalized retrocopies in individual cell lines. Only retrocopies with RPKM ≥ 1 were included. $^{***}p < 0.001$, NS. not significant; two-tailed Wilcoxon test.

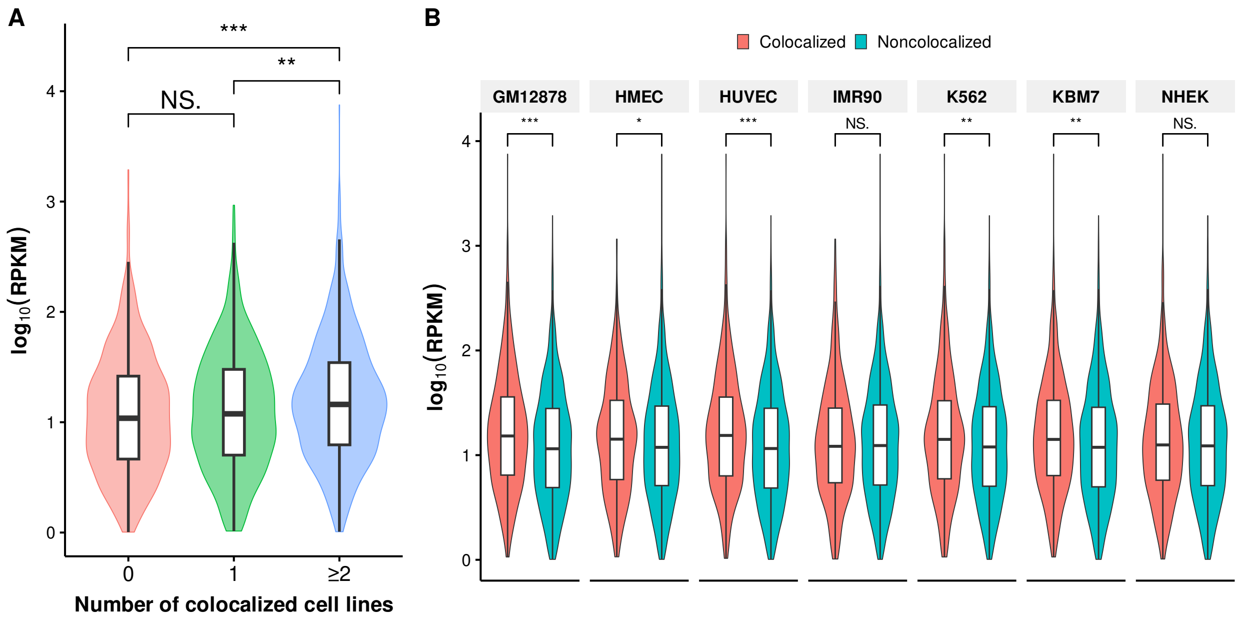


**Figure S27.** Comparison of ribo-seq RPKM values among retrocopies that are colocalized in 0, 1, and ≥ 2 cell lines (A) and between colocalized and noncolocalized retrocopies in individual cell lines (B). Only retrocopies with RPKM ≥ 1 and with at least one detected ORFs were included. $^{***}p < 0.001$, $^{**}p < 0.01$, $^{*}p < 0.05$, NS. not significant; two-tailed Wilcoxon test.

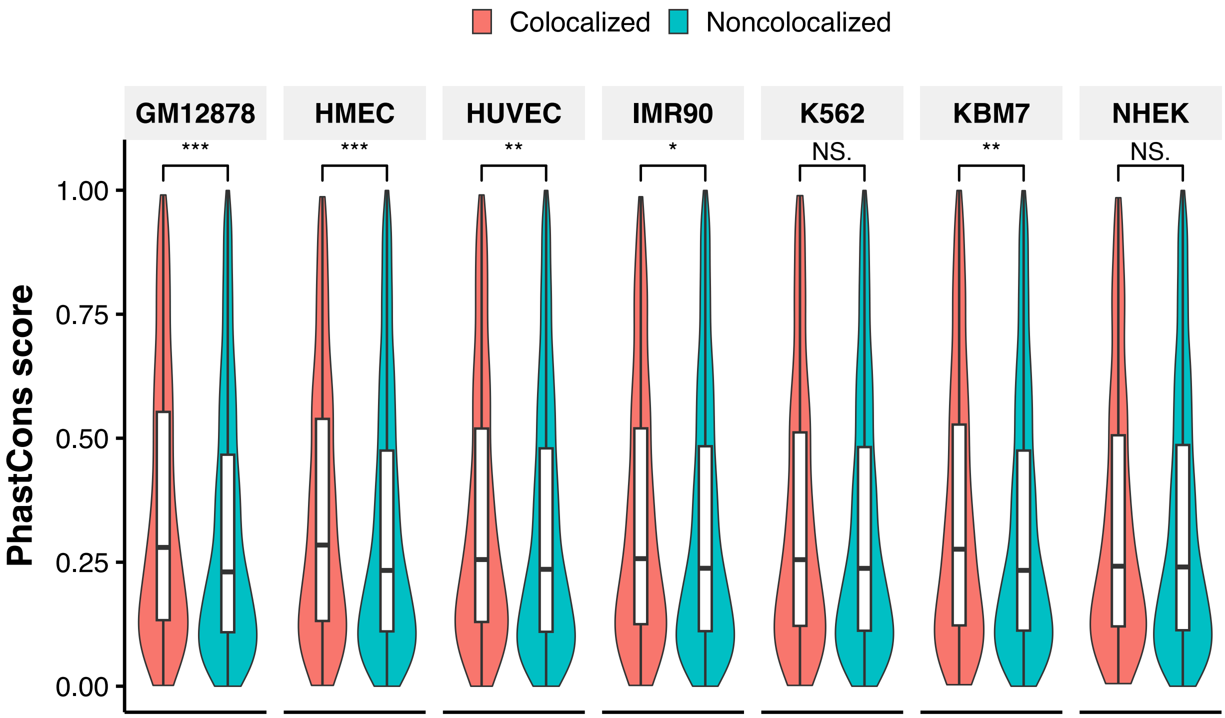


**Figure S28.** Comparison of phastCons scores between colocalized and noncolocalized retrocopies in individual cell lines. $^{***}p < 0.001$, $^{**}p < 0.01$, $^{*}p < 0.05$, NS. not significant; two-tailed Wilcoxon test.

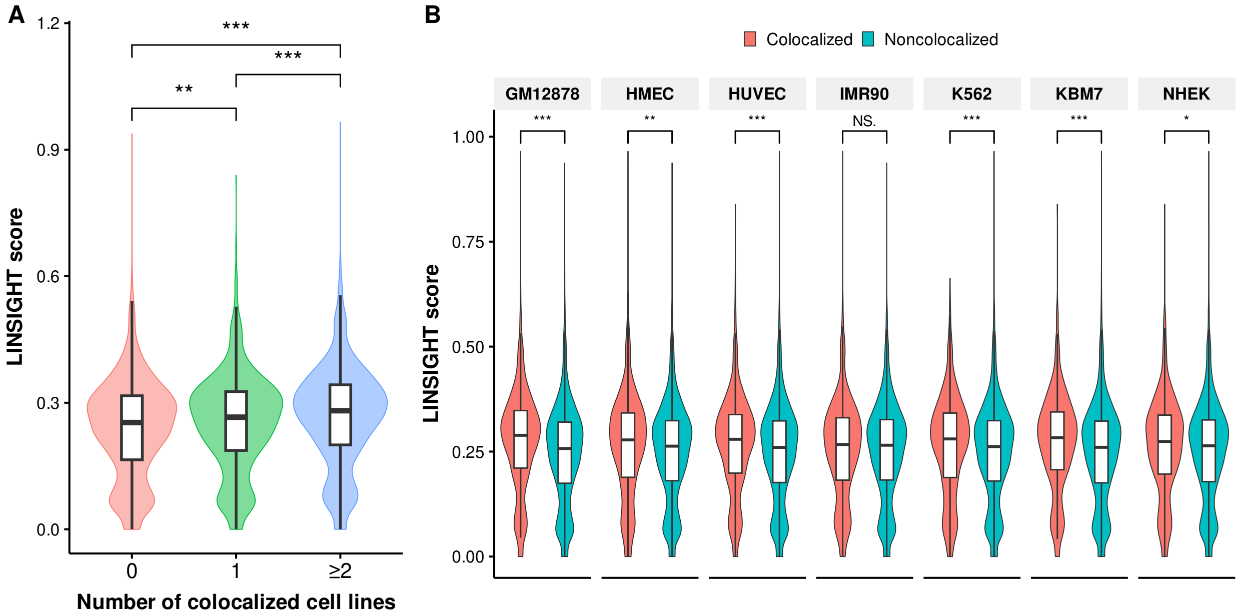


**Figure S29.** Comparison of LINSIGHT scores among retrocopies that are colocalized in 0, 1, and ≥ 2 cell lines (A) and between colocalized and noncolocalized retrocopies in individual cell lines (B). $^{***}p < 0.001$, $^{**}p < 0.01$, $^{*}p < 0.05$, NS. not significant; two-tailed Wilcoxon test.

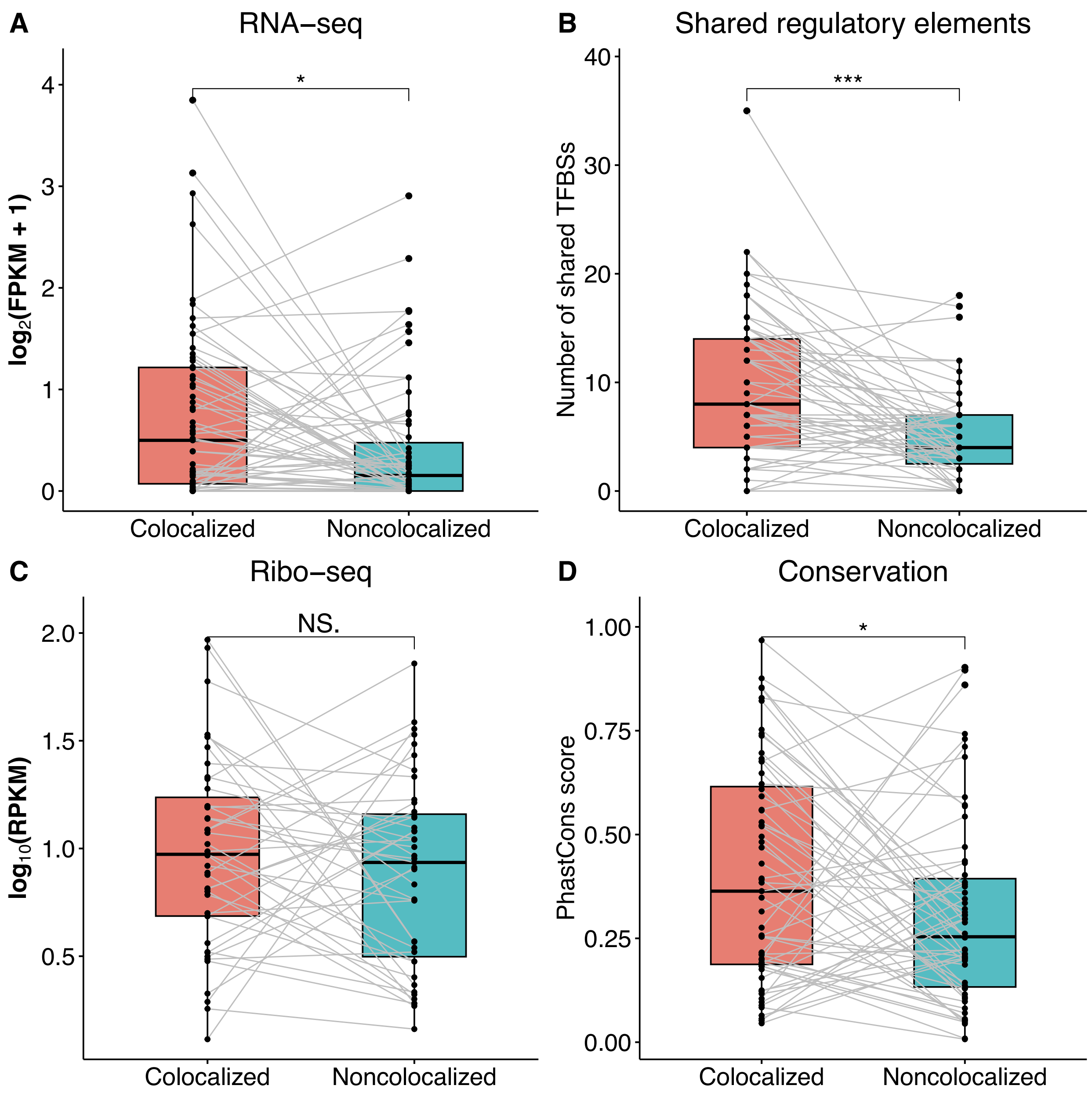


**Figure S30.** Comparison of RNA expression levels (A), number of shared motifs (B), ribo-seq RPKM values (C), and conservation scores for paired colocalized and noncolocalized retrocopies derived from the same parental genes. The colocalization status, levels of transcription and translation are based on data in the GM12878 cell line. These parental genes have exactly two retrocopies, one is colocalized with parent and the other is not colocalized with parent. $^{***}p <$ 0.001, $^{*}p < 0.05$, NS. not significant; paired t-test.

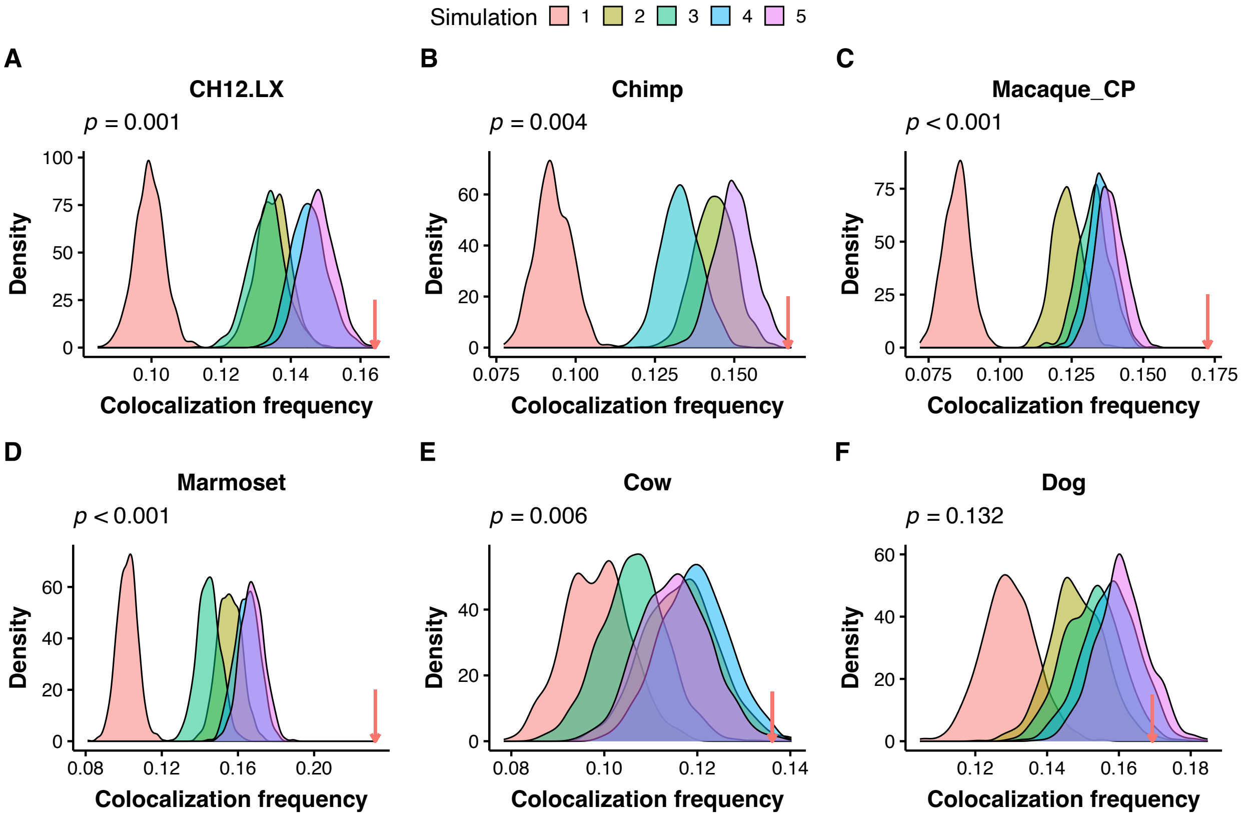


**Figure S31.** Spatial colocalization between retrocopies and their parental genes in mouse CH12-LX cell line (A), chimpanzee iPSC (B), rhesus macaque cortex plate (C), marmoset iPSC (D), cow skin ear tissue (E), and dog skin ear tissue (F). For each species, the density plot showing the distribution of colocalization frequency between chromatin pairs under different null models, while the red arrow denotes the colocalization frequency between true retrocopy-parent pairs. For chimpanzee, due to the lack of Ensembl annotation for panTro6 and the NCBI annotation containing no information of the gene type, we did not carry out simulation 4 in this species. *P*-values are given for simulation 5.

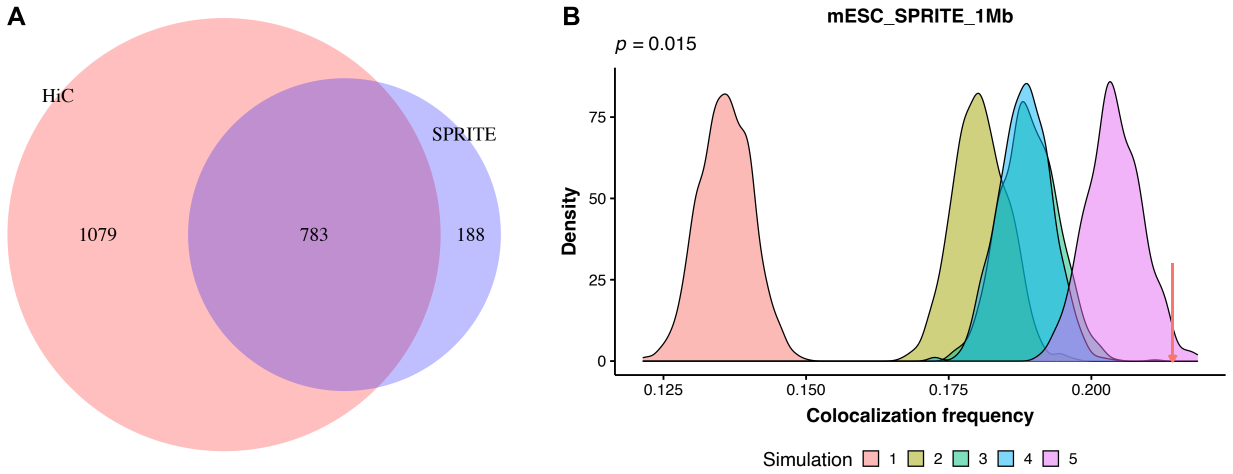


**Figure S32.** For the mouse mESC cell line, (A) displaying the overlap of colocalized retrocopy-parent pairs that are derived from significant Hi-C and SPRITE chromatin interactions and (B) showing the distribution of colocalization frequency between chromatin pairs under different null models when using chromatin contact map derived from SPRITE. The red arrow denotes the colocalization frequency of true retrocopies and their parental genes. *P*-value is indicated for simulation 5. The resolution of both Hi-C and SPRITE is 1 Mb.

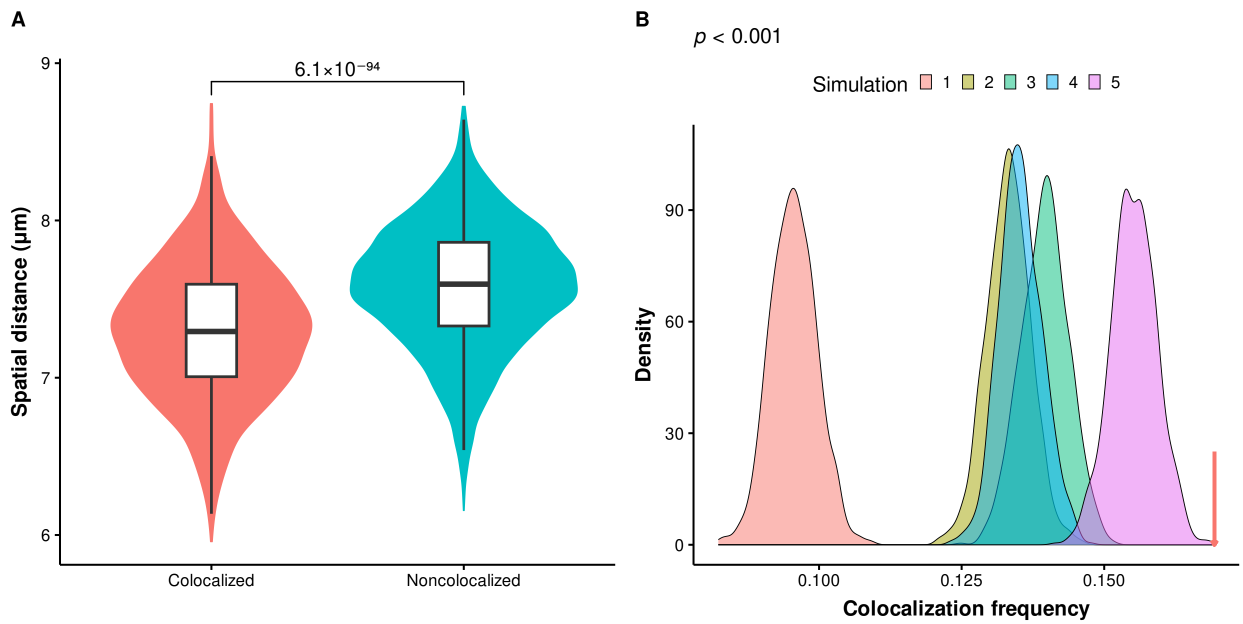


**Figure S33.** For the mouse mESC cell line, (A) showing the comparison of mean spatial distance determined by seqFISH+ between colocalized and noncolocalized retrocopy-parent pairs detected by Hi-C. The *p*-value of two-tailed Wilcoxon test is indicated. (B) illustrating the distribution of colocalization frequency between chromatin pairs under different null models when the top 10% interchromosomal chromatin pair with the shortest spatial distances is regarded as significant colocalization. The red arrow denotes the colocalization frequency of true retrocopies and their parental genes. The resolution of seqFISH+ chromatin contact map is 200 kb. *P*-values < 0.001 in all simulations.

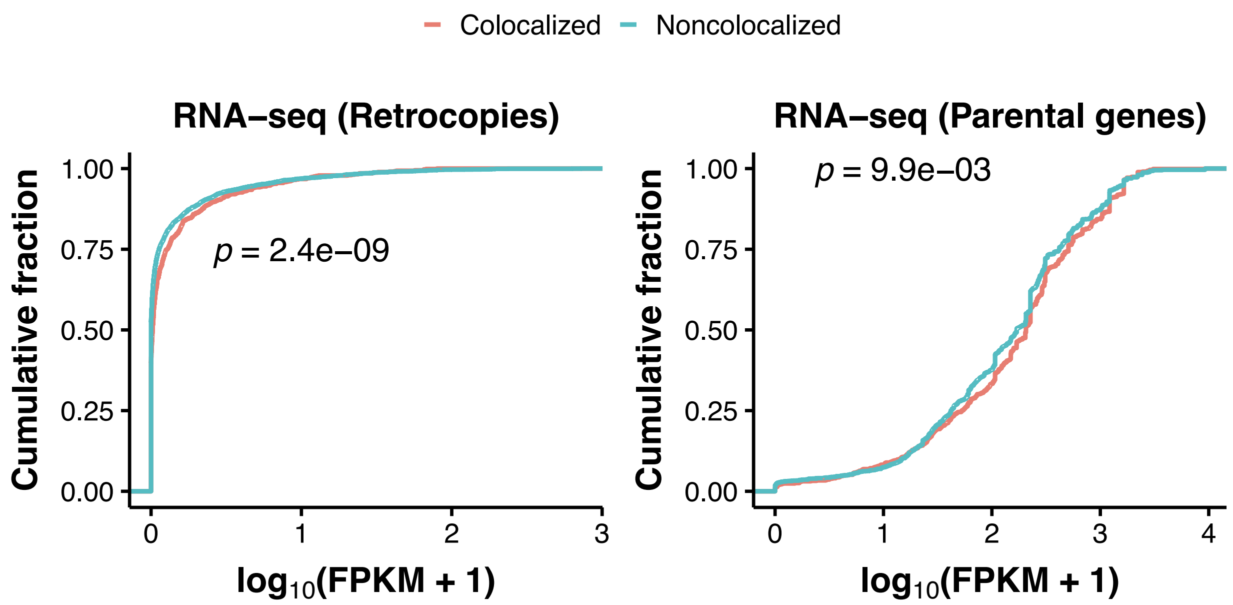


**Figure S34.** Plots of cumulative fraction of expression of retrocopies (left panel) and parental genes (right panel) in the mouse CH12-LX cell line. Expression level was measured from polyadenylated (polyA+) RNAs from the whole cell. Two-tailed Wilcoxon tests were carried out to compare the expression differences between colocalized retrocopies/parental genes and their noncolocalized counterparts, and the associated *p*-values are indicated in each plot.

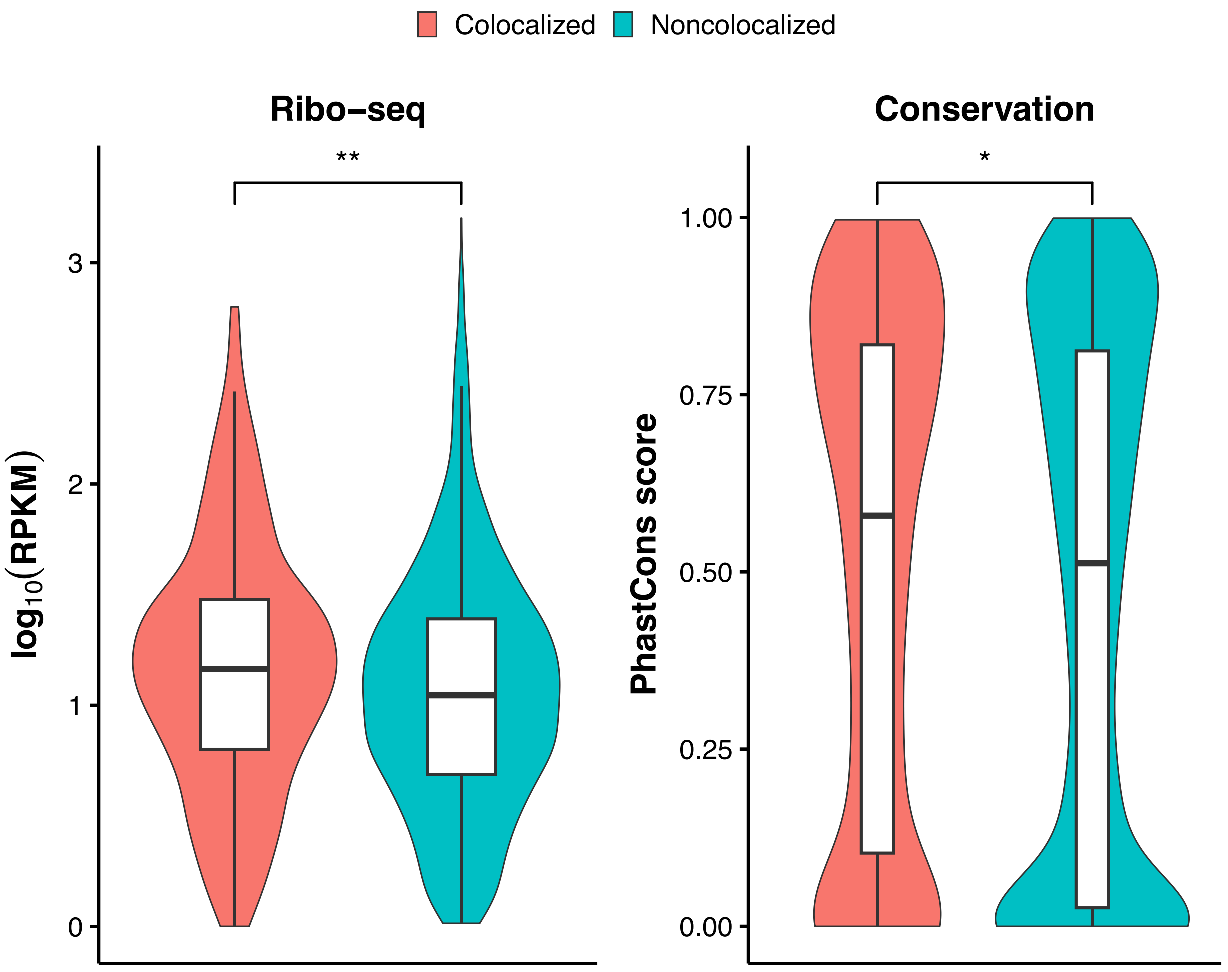


**Figure S35.** Comparison of ribo-seq RPKM values (left panel) and phastCons scores (right panel) between colocalized and noncolocalized retrocopies in the mouse CH12-LX cell line. For ribo-seq data, only retrocopies with RPKM ≥ 1 were included. $^{**}p < 0.01$, $^{*}p < 0.05$; two-tailed Wilcoxon test.

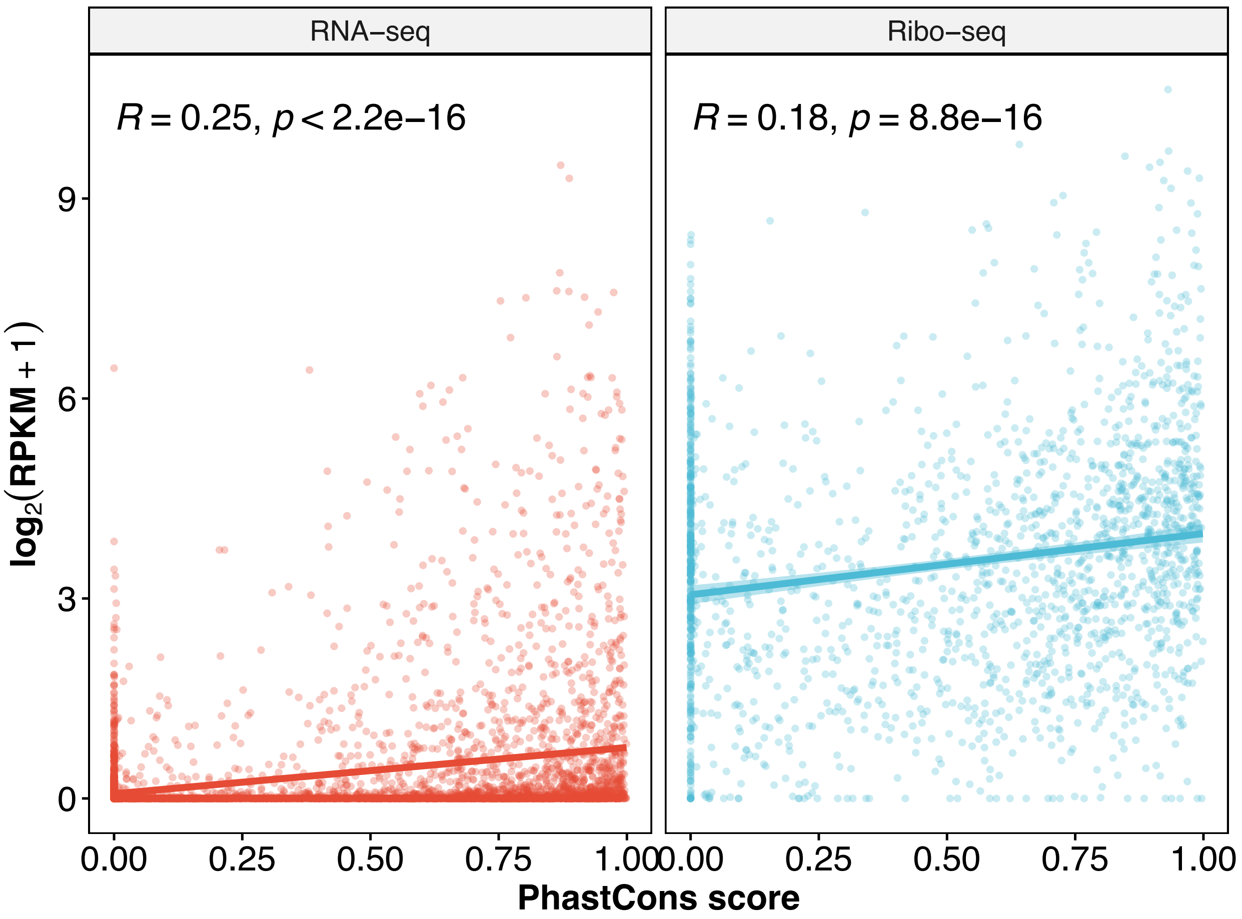


**Figure S36.** Correlation between phastCons score and RNA expression level (left panel) and between phastCons score and ribo-seq signal (right panel) of retrocopies in mouse. Solid line and shadowed area in each plot denote the linear regression line and its confident interval. Pearson correlation coefficient and its associated *p*-value is given at the top left corner of each plot.

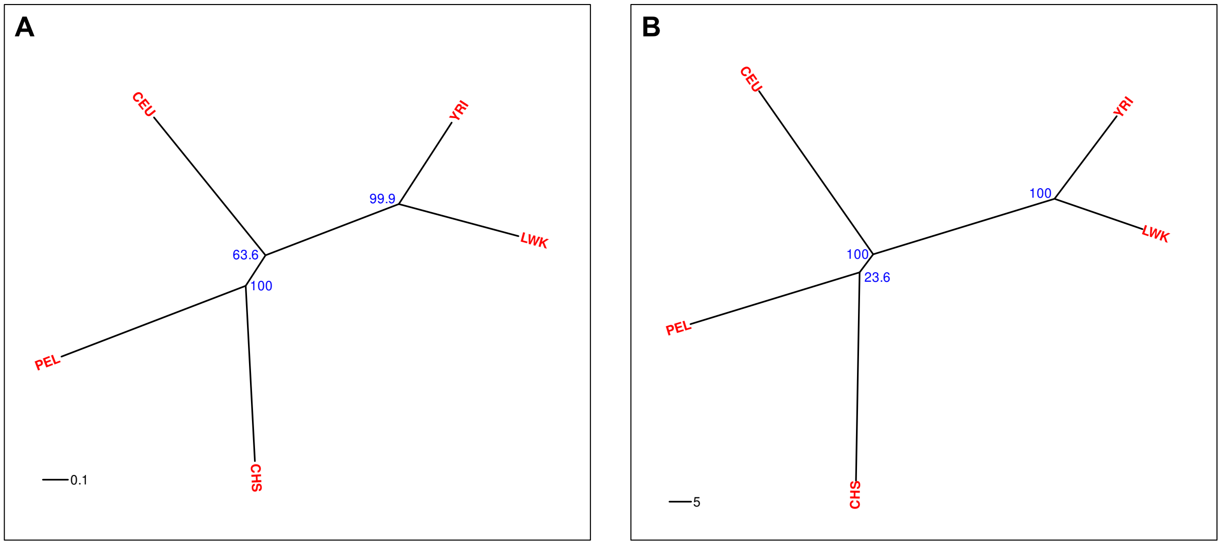


**Figure S37.** Neighbor-joining trees built on the basis of the presence fraction spectrum of retroCNVs within each population (A) and presence fraction spectrum of SNPs (B) in the same set of populations from the 1000 Genomes Project. Bootstrap support value is given for each node.

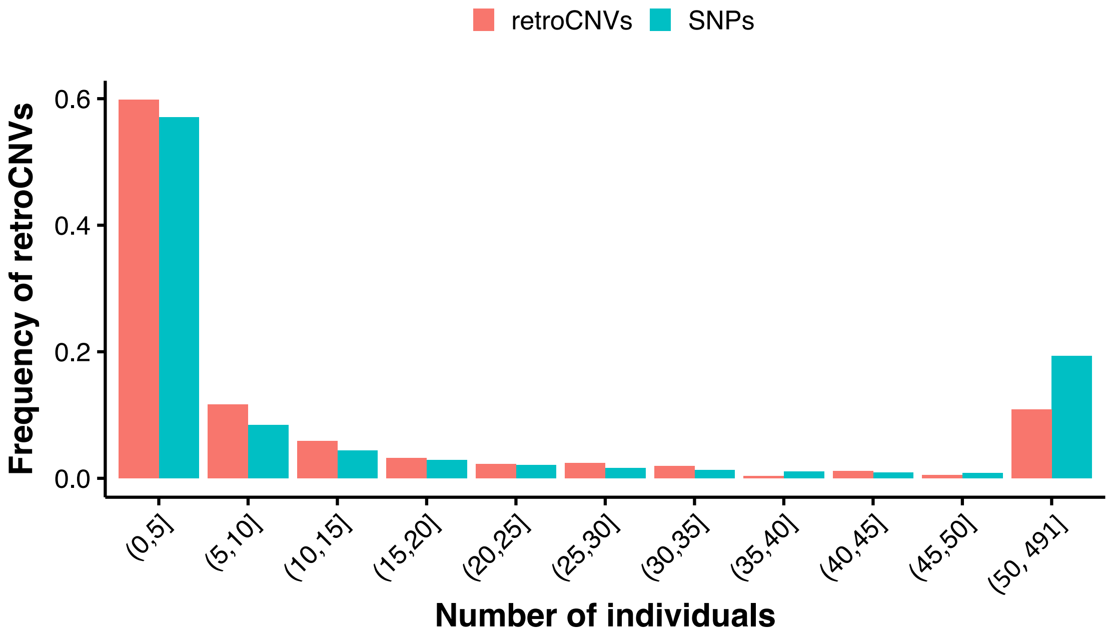


**Figure S38.** Presence frequency spectra of 625 retroCNVs (red) and 38,631,377 SNPs in 491 individuals of 5 human populations. For both retroCNVs and SNPs, only the presence and absence of an allele was accounted, i.e., without distinguishing homozygous and heterozygous status.

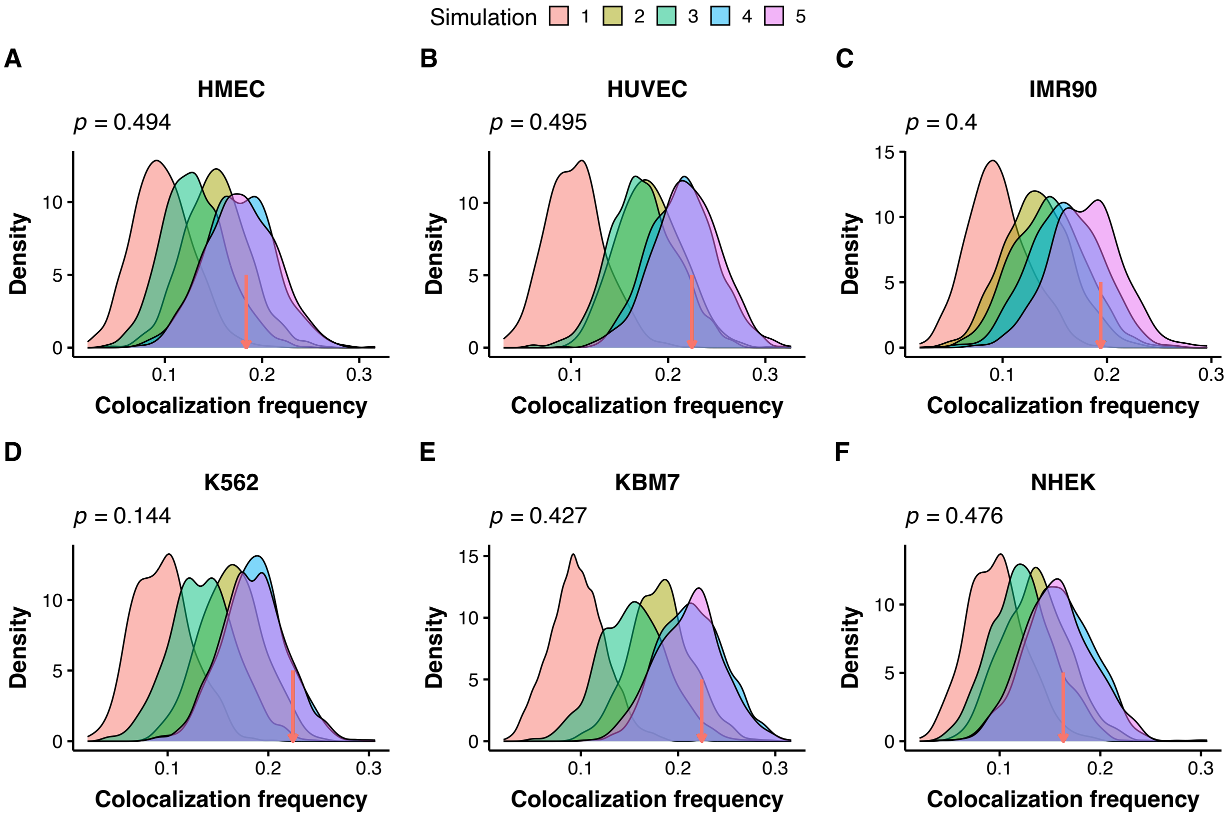


**Figure S39.** Density plots showing the distribution of colocalization frequency between chromatin pairs under different null models for singleton retroCNVs. Red arrows denote the colocalization frequency between true retroCNVs and their parental genes. Simulations and true frequency are based on Hi-C data in HMEC (A), HUVEC (B), IMR90 (C), K562 (D), KBM7 (E), and NHEK (F). *P*-value of simulation 5 is shown at the top left in each plot.

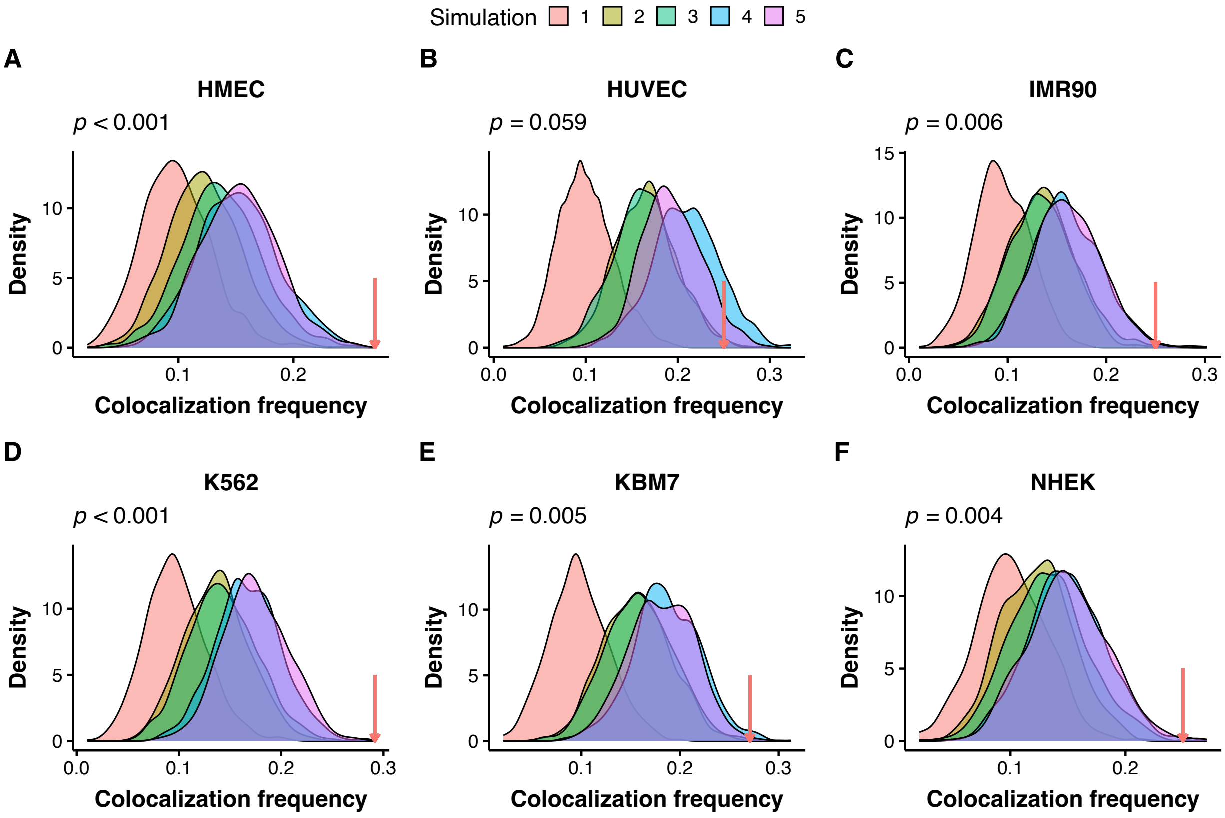


**Figure S40.** Density plots showing the distribution of colocalization frequency between chromatin pairs under different null models for high-frequency retroCNVs. Red arrows denote the colocalization frequency between true retroCNVs and their parental genes. Simulations and true frequency are based on Hi-C data in HMEC (A), HUVEC (B), IMR90 (C), K562 (D), KBM7 (E), and NHEK (F). *P*-value of simulation 5 is shown at the top left in each plot.

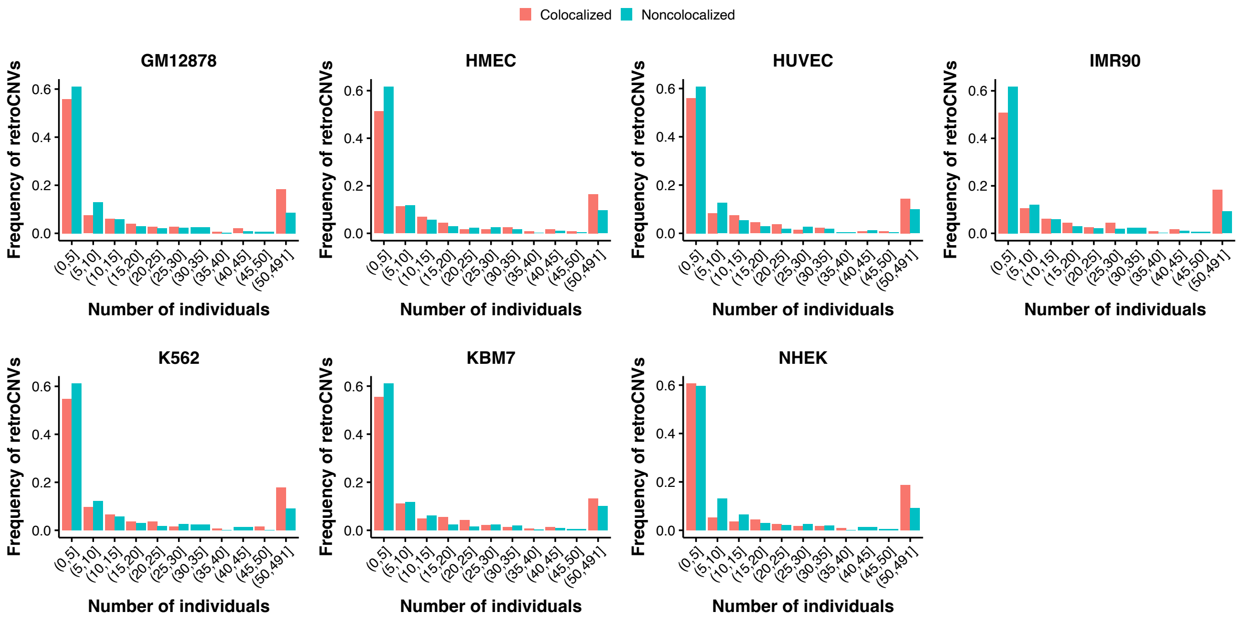


**Figure S41.** Presence frequency spectrum for retroCNVs that are colocalized and noncolocalized in individual cell lines.

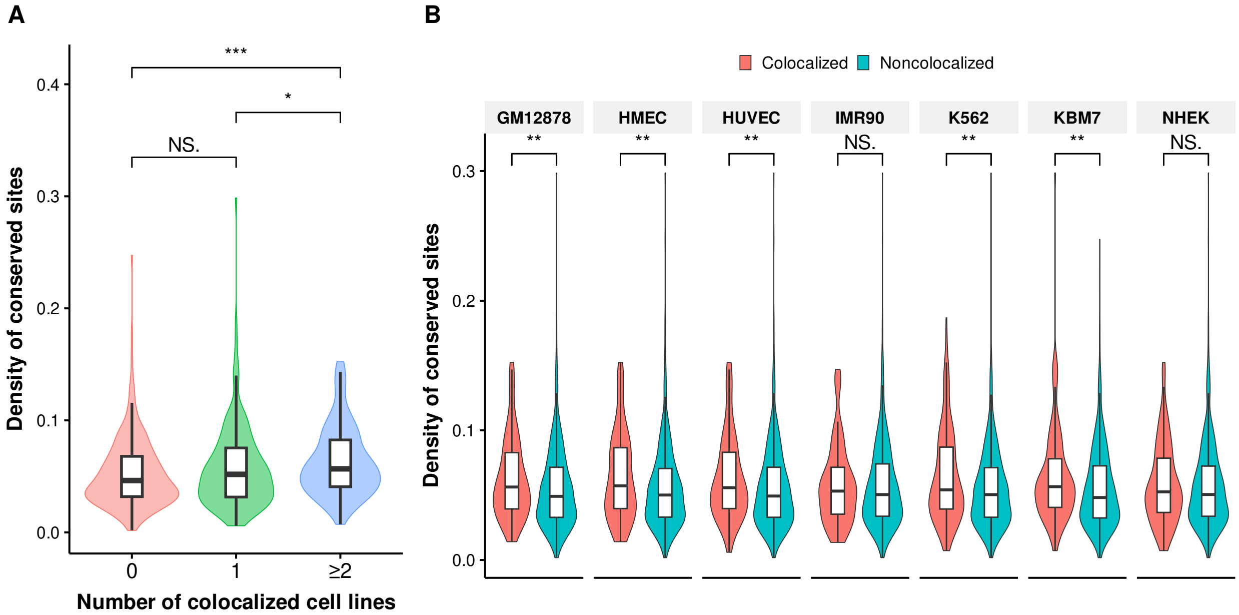


**Figure S42.** Comparison of the density of conserved sites (phastCons score > 0.5) in flanking regions (insertion site ± 50 kb) among retroCNVs that are colocalized in 0, 1, and ≥ 2 cell lines (A) and between colocalized and noncolocalized retrocopies in individual cell lines (B). $^{***}p <$ 0.001, $^{**}p < 0.01$, $^{*}p < 0.05$, NS. not significant; two-tailed Wilcoxon test.

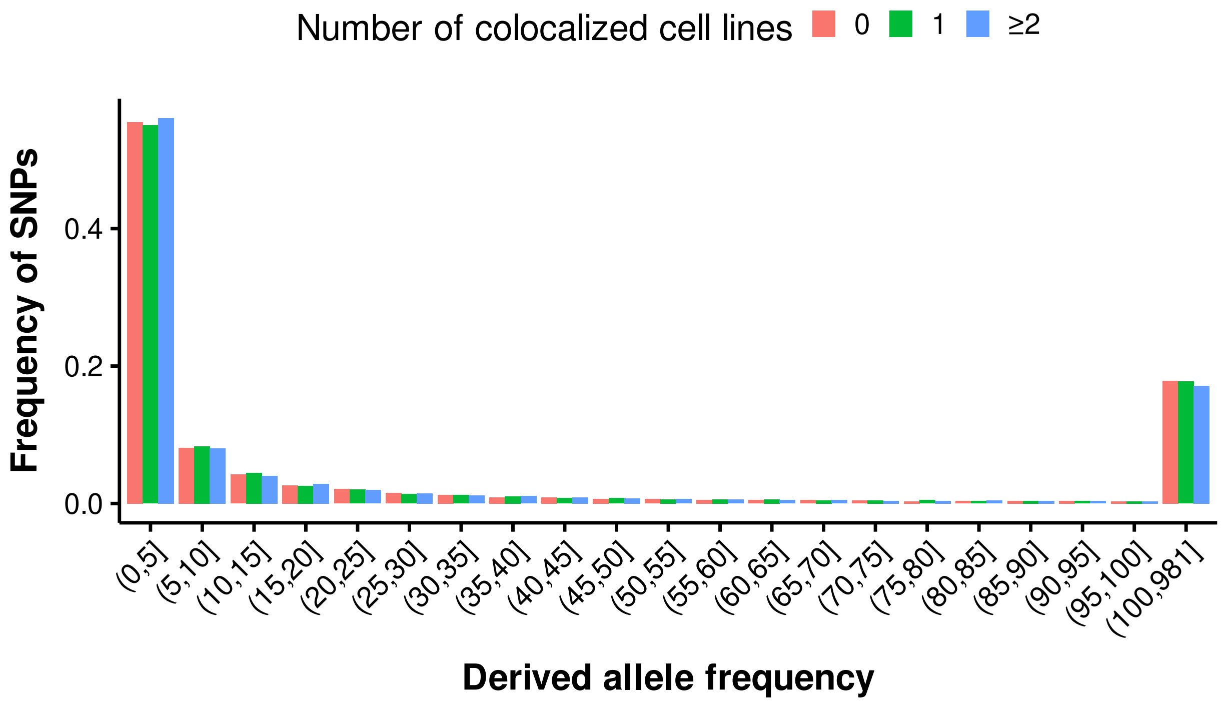


**Figure S43.** Unfolded site frequency spectra for SNPs flanking (insertion site ± 50 kb) retroCNVs that are colocalized in 0, 1, or ≥ 2 cell lines.

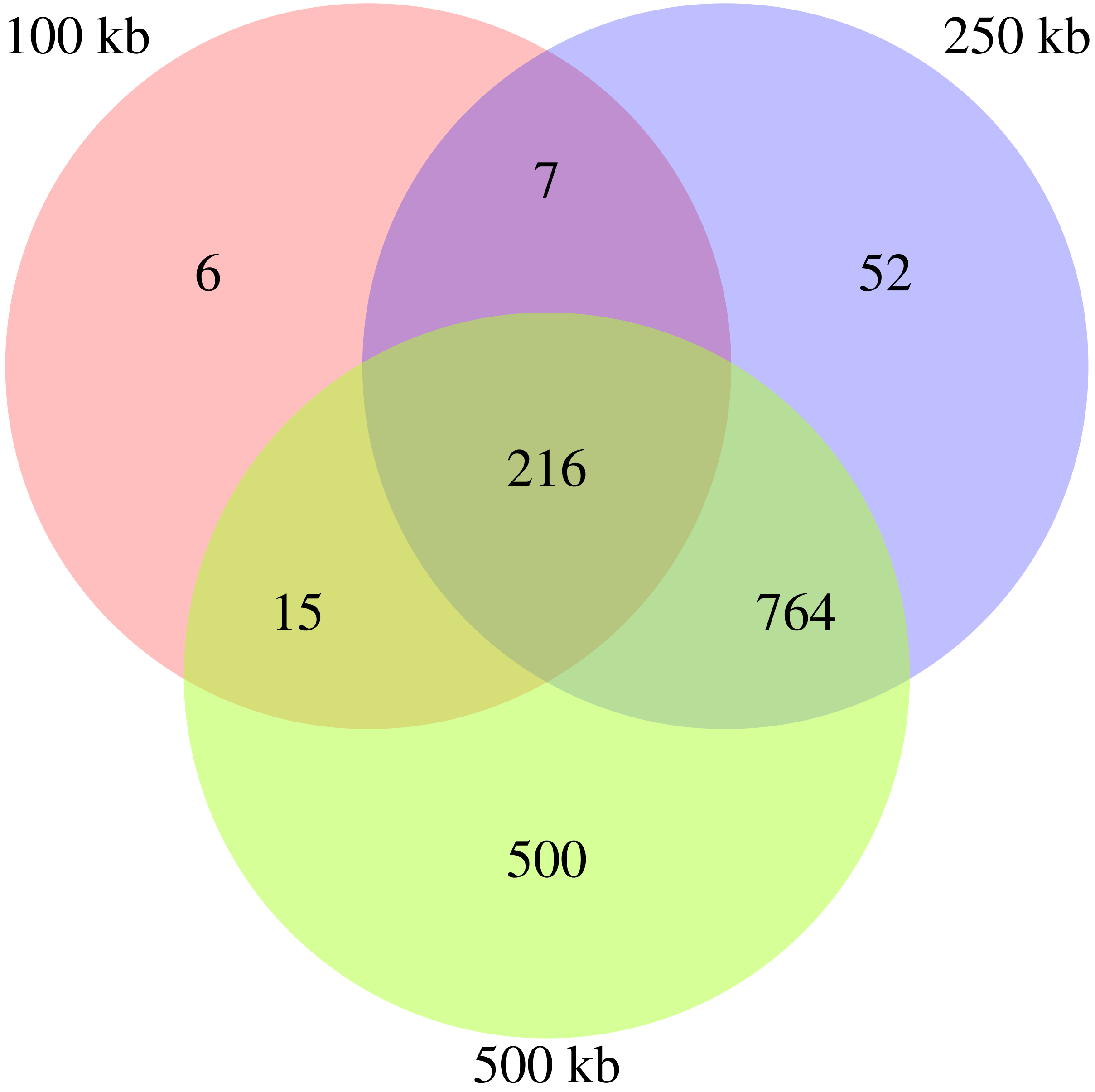


**Figure S44.** Venn diagram showing the overlap of colocalized retrocopy-parent pairs under different resolutions (100-kb, 250-kb, and 500-kb) in the GM12878 cell line.

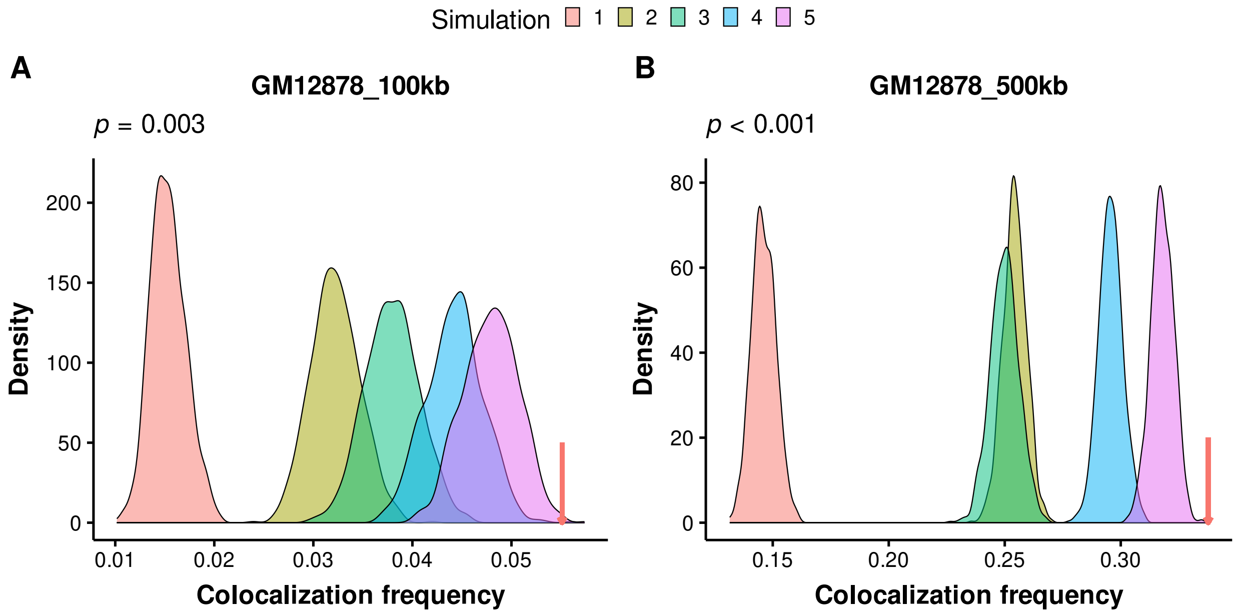


**Figure S45.** Density plots showing the distribution of colocalization frequency between chromatin pairs under different null models for GM12878 at 100-kb (A) and 500-kb (B) resolutions, while the red arrow denotes the colocalization frequency of true retrocopy-parent pairs. *P*-value of simulation 5 is indicated in each plot; *p*-values < 0.001 for the other four simulations in both plots.
